# Single-Cell Integration of Chromatin Accessibility and Transcriptomics Reveals Regulatory Networks in Ovarian Tumor-Infiltrating Adaptive NK Cells

**DOI:** 10.1101/2025.09.13.676040

**Authors:** Yizhe Sun, Meng Wan, Solrun Kolbeinsdottir, Kang Wang, Shruti Khare, Okan Gultekin, Sahar Salehi, Kaisa Lehti, Ramanuj Dasgupta, Theodoros Foukakis, Martin Enge, Dhifaf Sarhan

## Abstract

Natural killer (NK) cells are traditionally recognized for their rapid, non-specific responses against virus-infected or malignantly transformed cells, functioning as key effectors of innate immunity. However, a distinct subset known as adaptive NK (aNK) cells has been shown to acquire memory-like properties following viral infections, indicating their capacity for antigen-specific immune recall. Intriguingly, aNK cells have also been identified within the tumor microenvironment, where they can mediate tumor-specific recall responses. Yet, the regulatory mechanisms governing their function in tumor-infiltrating aNK cells remain largely undefined. In this study, we integrated publicly available multiomics datasets from ovarian cancer, including single-cell chromatin accessibility (scATAC-seq) and single-cell RNA sequencing (scRNA-seq), to identify chromatin-accessible regions and construct transcription factors (TF)-gene regulatory networks. To validate and extend these findings, we performed Smart-seq3 on NK cells isolated from ovarian tumors and applied SCENIC analysis to identify TF-driven gene regulation. By integrating results from both analyses, we identified PRDM1 and STAT2 as key TFs, along with their downstream targets *CRCP* and *MTFP1*, specifically enriched in tumor-infiltrating aNK cells. The expression levels of *CRCP* and *MTFP1* positively correlated with NK cell infiltration in ovarian cancer tissues, suggesting their potential functions in supporting tumor-specific NK cell memory responses. In addition, external validation using data from the PROMIX clinical trial demonstrated that *PRDM1* and *STAT2* expression levels are positively associated with both overall survival and aNK cell-associated transcriptional features.

**Graphical abstract:** 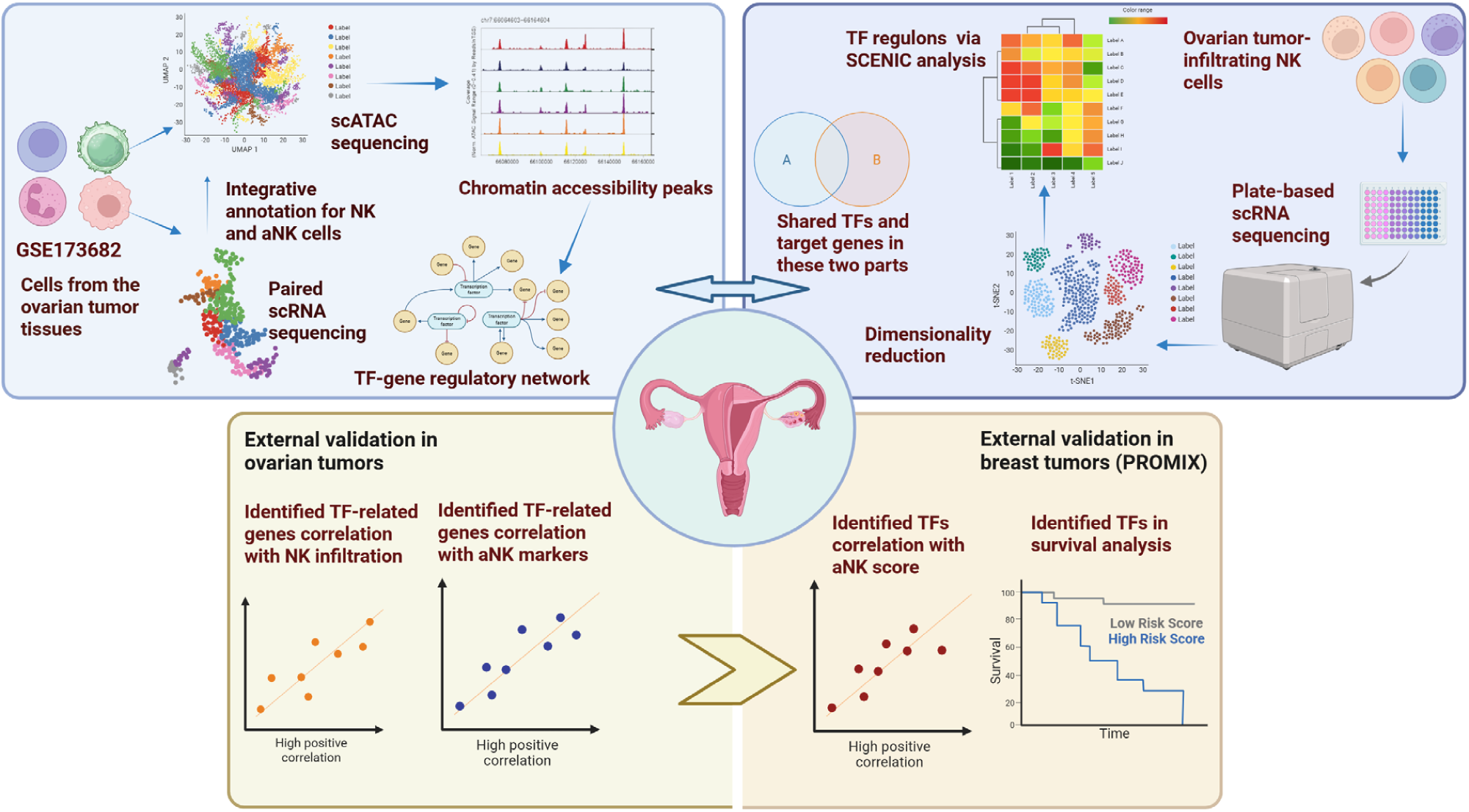

## Introduction

Natural killer (NK) cells are cytotoxic lymphocytes and key components of the innate immune system, capable of mounting rapid responses against virally infected, transformed, or stressed cells^[1]^. Unlike T lymphocytes, NK cells function independently of antigen-specific receptors and prior sensitization, relying instead on a finely balanced system of activating and inhibitory signals conveyed through germline-encoded receptors^[2]^.

Inhibitory receptors, such as killer-cell immunoglobulin-like receptors (KIRs) and the heterodimeric CD94/NKG2A, are essential for maintaining self-tolerance by recognizing MHC class I molecules^[3]^. KIRs specifically engage polymorphic epitopes on classical MHC-I molecules, including HLA-C1, HLA-C2, and HLA-Bw4^[4]^, while NKG2A recognizes the non-classical MHC-I molecule HLA-E^[5]^, which presents conserved leader peptides derived from classical MHC-I molecules. These inhibitory pathways prevent dysfunctional NK cell activation under physiological conditions. In contrast, activating receptors such as NKG2D and the natural cytotoxicity receptors (NKp30, NKp44, NKp46) recognize stress-induced ligands that are upregulated upon infection, malignancy, or cellular distress^[6]^. Additionally, CD16 (FcγRIIIa) enables NK cells to mediate antibody-dependent cellular cytotoxicity (ADCC) by binding the Fc region of IgG antibodies, thereby bridging innate and adaptive immune responses^[7]^.

However, this binary classification of innate versus adaptive immunity has been increasingly challenged by accumulating evidence that NK cells can develop memory-like properties under specific conditions. Most notably, during viral infections such as human cytomegalovirus (HCMV), subsets of NK cells can persist long-term, undergo clonal-like expansion, and mount antigen-specific recall responses, hallmarks of adaptive immunity^[8–10]^. These discoveries redefine NK cells as not merely transient effectors but also potential contributors to long-lived immune surveillance. This paradigm shift is particularly relevant in the context of cancer, where the tumor microenvironment (TME) may similarly imprint long-lasting changes on NK cell identity and function. Understanding whether, and how, NK cells acquire such memory-like features in tumors remains a critical and unresolved question^[11–14]^.

Adaptive NK (aNK) cells induced by HCMV infection exhibit reduced expression of inhibitory NKG2A and increased expression of NKG2C, along with durable changes in chromatin accessibility and metabolism^[10–14]^. These features reflect underlying epigenetic reprogramming, including the enrichment of AP-1 motifs and a reduction in DNA methylation at key effector genes (e.g., *IFNG, GZMB, PRF1*). Studies have identified transcription factors (TFs) such as *Fli1*, acting in coordination with *STAT5*^[15]^, and *Bcl11b*^[16]^, as key regulators closely associated with aNK cell identity. Such memory-like NK cells are functionally distinct, displaying increased IFN-γ production and augmented cytotoxicity upon reactivation^[17–19]^.

While these phenomena have been well-characterized in viral contexts, their relevance to solid tumors is less clear. Recent pan-cancer single-cell atlases have identified tumor-infiltrating NK cell populations expressing adaptive markers, suggesting memory-like features may also emerge within the TME^[15, 16]^. Moreover, our previous work demonstrated that expanded aNK cells within ovarian tumors exhibited tumor-specific recall responses and cytotoxicity, and that naïve NK cells could be trained *ex-vivo* to recognize primary ovarian tumor antigens ^[20]^. These findings support the possibility that aNK cell states may also arise in tumors, though the regulatory mechanisms remain undefined.

We hypothesized that tumor-infiltrating NK cells may undergo specific epigenetic reprogramming to adopt an adaptive, memory-like state, enabling them to mount persistent and antigen-specific anti-tumor responses.

To address this gap, we performed an integrative multiomic analysis focused on aNK cells in the ovarian cancer microenvironment. We re-analyzed publicly available scATAC-seq and RNA-seq data to define chromatin-accessible regions and NK cell subpopulations, and we generated full-length transcriptomes of tumor-infiltrating NK cells using Smart-seq method. Using SCENIC, we reconstructed regulatory networks defining aNK identity and identified *PRDM1* and *STAT2* as key transcription factors, supported by both chromatin accessibility and regulon activity. *CRCP* and *MTFP1* emerged as putative downstream targets, exhibiting elevated expression in aNK cells and association with increased NK infiltration across independent datasets. Moreover, *PRDM1* and *STAT2* correlated with improved prognosis and aNK-like transcriptional signature in HER2-negative breast cancer. These results suggest a conserved transcriptional circuit driving memory-like NK states in tumors with potential relevance for immunotherapy.

## Methods and Materials

### Workflow of scATACseq data Analysis

scATAC-seq data from six human samples were processed using the ArchR package(v1.0.3) ^[21]^. Fragment files were converted into Arrow files using createArrowFiles, with both TileMatrix and GeneScoreMatrix generated. The mitochondrial (chrM) and sex chromosomes (chrX and chrY) were excluded from downstream analysis. An ArchRProject was initialized using the hg38 human genome reference, and the entire workflow was carried out within this project.

To ensure high-quality data, we applied doublet detection using addDoubletScores followed by filtering with filterDoublets. For each sample, Gaussian Mixture Models **(GMMs)** were fitted separately to log-transformed fragment counts (nFrags) and Transcription Start Site (TSS) enrichment scores using the mclust package. Cells classified as high-quality in both distributions, with classification uncertainty ≤ 0.05, were retained. The intersection of cells passing both filters was used for subsequent analysis.

Dimensionality reduction was performed using iterative Latent Semantic Indexing (**LSI**) via addIterativeLSI, with four iterations, 25,000 variable features, and 50 components retained. Clustering was performed using the Seurat Louvain method (addClusters) with resolution set to 0.7. UMAP embedding was computed based on the LSI dimensions using cosine distance (addUMAP, n_neighbors = 30, min_dist = 0.3), and cluster distribution was visualized in UMAP space.

To assign cell type identities to the scATAC-seq clusters, we performed label transfer from a matched scRNA-seq dataset using addGeneIntegrationMatrix. Gene activity profiles from the GeneScoreMatrix were aligned with a reference Seurat object containing scRNA-seq data, and cell type labels were transferred. Predicted cell types (predictedGroup) were used to annotate the ATAC clusters by determining the cell identity across all the clusters. Simultaneously The expression of NK cell markers (e.g., *NCAM1, KLRC1, FCGR3A, KLRK1*) was also visualized in UMAP space using the imputed gene expression matrix.

To identify reproducible peaks across NK cell subtypes, we first grouped cells by their final annotations and generated pseudo-bulk replicates using addGroupCoverages, requiring a minimum of 50 cells per group. Peak calling was performed using addReproduciblePeakSet, with MACS2 (v2.2.9.1) ^[22]^ as the backend peak caller and a minimum threshold of 500 peaks per cell. The resulting unified peak set was used to construct the PeakMatrix for each cell.

To identify marker peaks specific to each cluster, we used getMarkerFeatures on the PeakMatrix, controlling for potential bias from TSS enrichment and sequencing depth. Marker peaks were defined using Wilcoxon testing with thresholds of FDR ≤ 0.1 and log2 fold change ≥ 0.25. Peaks from selected clusters (e.g., “Adaptive_NK_cells” and “Conventional_NK_cells”) were retained and further processed. All marker peak results were consolidated, filtered, and exported as both .rds and .csv files for downstream interpretation.

To investigate the genomic distribution of cis-regulatory elements, we annotated the identified peaks using the ChIPseeker R package (v1.44.0) ^[23]^ and GenomicRanges package (v1.60.0) ^[24]^. Unique peaks defined by chromosomal coordinates (seqnames, start, end) were first converted into a GRanges object. Peak annotation was performed with the annotatePeak function, using TxDb.Hsapiens.UCSC.hg38.knownGene as the transcript database and org.Hs.eg.db for gene annotation. Promoter regions were defined as ±3 kb from the TSS. The resulting annotations were classified into functional categories, including promoter (±3 kb), 5′ UTR, 3′ UTR, exon, intron, downstream, and distal intergenic regions, to determine the genomic distribution of peaks across cis-regulatory elements.

We then identified peak-to-gene regulatory relationships by computing correlations between peak accessibility and gene expression using addPeak2GeneLinks, with a Pearson correlation cutoff of 0.4 and a genomic window of ±250 kb. Peaks with significant correlations were linked to target genes, and the results were visualized using plotBrowserTrack.

To identify potential regulatory TFs, we integrated motif annotation data (Motif-Matches-In-Peaks.rds, which was generated via addMotifAnnotations with the CIS-BP database, and the binary motif occurrence matrix was created via addMotifMatrix, resulting in the file Motif-Matches-In-Peaks.rds stored under the ArchR project’s *Annotations* directory) with the peak-to-gene links. Logical motif match matrices were filtered to retain only those peaks linked to genes, and motif matches were aggregated per peak. Each peak was annotated with a list of matched TFs and merged with its associated gene and cluster identity. This allowed us to generate a regulatory TF–peak–gene–cluster map. Frequency analysis of motif occurrences across clusters revealed distinct TF motif enrichment patterns in aNK cell populations.

### Workflow for pySCENIC analysis

Gene regulatory network inference and regulon activity analysis were performed using pySCENIC (v0.11.2) ^[25]^. First, co-expression modules were inferred from the filtered single-cell expression matrix using the GRNBoost2 algorithm via the pyscenic grn command, with a curated list of human transcription factors (**allTFs_hg38.txt**) as input. The resulting adjacency matrix was then used for motif enrichment analysis using pyscenic ctx, with the hg38 motif ranking database (**hg38 refseq- r80 10kb_up_and_down_tss.mc9nr.genes_vs_motifs.rankings.feather)** and motif annotation table (**motifs-v9-nr.hgnc-m0.001-o0.0.tbl**). This step generated a set of high-confidence regulons stored in output csv file. Finally, regulon activity scores were computed per cell using pyscenic aucell, resulting in a SCENIC-annotated loom file for downstream visualization and clustering.

Regulon specificity scores **(RSS)** were computed for each cluster to identify key TFs, and visualizations included heatmaps, RSS ranking plots, and cluster-specific TF activity UMAPs. Z-score–normalized regulon activity was used to further explore TF specificity across clusters using heatmaps and violin plots. The final integrated dataset allowed the visualization and comparison of candidate regulons (e.g., STAT2(+), PRDM1(+), BCL11A(+), SP1(+), etc.) across NK cell subtypes.

### Pre-processing and annotations for scATAC-seq-matched scRNA dataset

Count matrices were processed with Seurat (v4.3.0). Cells with fewer than 3 detected genes were discarded, and quality control was applied by filtering out outliers in library size, gene number, and mitochondrial content (>2 MAD from the median, scater). After filtering, data were log-normalized, 2,000 variable genes were identified using the variance-stabilizing method, and expression values were scaled. Dimensionality reduction was performed with PCA, followed by graph-based clustering (resolution = 0.7) and UMAP visualization.

For automated cell annotation, the Seurat object was converted to a SingleCellExperiment and analyzed with SingleR (v2.10.0)^[26]^. Reference datasets included the Human Primary Cell Atlas (HPCA) and BlueprintEncode (BED) from the Celldex package (v1.18.0)^[26]^. SingleR predictions were pruned to the main label level. To define NK cells, we integrated both references: cells annotated as NK by either HPCA or BlueprintEncode were classified as “NK cells,” while all remaining cells were categorized as “Other.”

For aNK cells annotation, NK cell subsets were annotated using the scType framework^[27]^. A custom NK reference gene set described in the results part was processed with gene_sets_prepare() to generate positive and negative markers, and enrichment scores were calculated on the scaled RNA expression matrix (sctype_score()). Cluster-level scores were aggregated across Seurat clusters, with the highest-scoring type assigned as the annotation; clusters with low-confidence scores (<25% of cluster size) were labeled as Unknown.

### Enzymatic digestion of IPLA tumors and single-cell sample production

Ovarian tumor samples were obtained from patients enrolled in the Phase III clinical trial Intra- Peritoneal Local Anesthetics in Ovarian Cancer (IPLA-OVCA; NCT04065009), which included patients undergoing upfront cytoreductive surgery for stage III or IV epithelial high-grade serous ovarian cancer (HGSOC) ^[28]^. All participants provided written informed consent prior to inclusion, in accordance with the Declaration of Helsinki. The study protocol was approved by the local Ethics Committee and Institutional Review Board (Approval numbers: Dnrs 2019-05149/updated as 2020-02888).

To get the single cell suspension from ovarian tumors, sections were cut into small pieces and digested using a harvest medium containing DNase I (100 µg/mL) (Sigma Alrich, Cat.#4716728001), and Liberase (150 µg/mL) (Sigma Aldrich, Cat. #5401127001) in RPMI medium at 37°C for 30 minutes. After digestion, the cell suspension was collected by passing the tissue through a 70 µm strainer. Red blood cell lysis was then performed, followed by two PBS washes, resulting in a tumor tissue suspension. All tumor suspensions were cryopreserved in 90% FBS and 10% DMSO.

### Smart-seq2 for ovarian tumor-derived NK cells

For Smart-seq2, ovarian tumor samples were enzymatically dissociated into single-cell suspensions, and dead cells were removed using the Dead Cell Removal Kit (Miltenyl Biotech, Cat.# 130-090-101). CD45 staining (BD biosciences, Cat. #5555483) was applied to separate CD45⁺ and CD45⁻ populations, which were individually sorted into 384-well plates, with two plates prepared per patient sample. The initial sample processing steps were performed according to the Direct nuclear tagmentation and RNA-sequencing (DNTR-seq) protocol (steps 1–4 and post-step 12)^[29]^. Cytosolic and nuclear fraction were separated, and only the cytosolic fraction was used in this paper. Primer annealing, reverse transcription and cDNA amplification were performed. cDNA was purified, tagmented and barcoded. Libraries were sequenced with paired-end sequencing, 2x150bp on a novaseq6000.

For Smart-seq2 library pre-processing, the raw sequencing reads were trimmed using Cutadapt (v3.2) ^[30]^ to remove adapter sequences and low-quality bases. Reads were then deduplicated with Picard MarkDuplicates (v2.22.0). Alignment to the reference genome was performed using STAR (v2.7.7a), and transcript counting was done with HTSeq (v 0.9.1).

Quality control thresholds were applied as follows: cells were retained only if they exhibited at least 500 features and at least 1,000 total counts. Additionally, an ACTB quantile cutoff at 0.005 was used to filter out low-quality or outlier cells. Finally, the processed Smart-seq2 data were ready for downstream analyses, such as normalization, integration and clustering using Seurat package (v4.3.0)^[31]^. NK cells were annotated across all clusters using SingleR (v2.10.0) , with the HPCA reference obtained via the Celldex (v1.18.0), and subsequently extracted for integration with the Smart-seq3 dataset.

### Smart-seq3 for ovarian tumor-derived NK cells

For Smart-seq3, ovarian tumor samples were enzymatically dissociated into single-cell suspensions, and NK cells were isolated by fluorescence-activated cell sorting **(FACS)** based on surface marker expression of CD45⁺CD3⁻CD56⁺.

Sorted NK cells were processed for full-length single-cell RNA sequencing using the Smart- seq3 protocol^[32]^, performed at the Single Cell Core Facility of Flemingsberg Campus (SICOF), Karolinska Institutet.

Individual NK cells were index-sorted into 384-well plates preloaded with lysis buffer containing oligo-dT primers, dNTPs, RNase inhibitors, and ERCC spike-ins. Reverse transcription was carried out using a template-switching oligonucleotide (**TSO**), enabling capture of full-length transcripts along with incorporation of unique molecular identifiers (**UMIs**) for absolute quantification. After reverse transcription and pre-amplification, cDNA was tagmented using Tn5 transposase and subjected to PCR amplification with indexed primers to generate sequencing-ready libraries.

Libraries were sequenced on an Illumina NovaSeq 6000 system using paired-end mode, targeting a sequencing depth of approximately 0.5–1 million reads per cell. Raw sequencing data were processed by the core facility using the standard Smart-seq3 pipeline. Briefly, reads were demultiplexed, and UMI-aware alignment was performed using STAR (v2.7.7a) to the human reference genome (hg38). Gene-level quantification was conducted using HTSeq (v 0.9.1), taking into account transcript start sites and UMI counts to reduce amplification bias. Cells with low library complexity, excessive mitochondrial gene expression, or low total UMI counts were excluded from downstream analysis. The final expression matrix was used for dimensionality reduction, clustering, and regulatory network analysis.

### Integration of Smart-seq2 and Smart-seq3 NK cell datasets

Tumor-infiltrating NK cells profiled by Smart-seq2 were integrated with Smart-seq3 datasets using Seurat (v4.3.0). Metadata were harmonized by adding patient labels and unique cell barcodes. Datasets were merged into a single Seurat object and processed with SCTransform normalization (5,000 variable features, regressing out mitochondrial gene percentage). Principal component analysis (PCA) was performed, and sample-specific differences were assessed by dimensionality plots and violin plots. To account for platform- and patient-related batch effects, Harmony was applied (θ = 2.5), and corrected embeddings were used for downstream analyses. Based on the top 13 Harmony dimensions, we constructed neighborhood graphs and performed non-linear dimensionality reduction with UMAP. Finally, unsupervised clustering was carried out using graph-based methods (resolution = 1.1), allowing joint characterization of NK cell heterogeneity across Smart-seq2 and Smart-seq3 datasets.

### Neoadjuvant chemotherapy (NAC) trial

The PROMIX trial (NCT00957125) enrolled patients with locally advanced (tumor >20 mm) HER2-negative breast cancer scheduled to receive six cycles of neoadjuvant chemotherapy (NAC) with epirubicin and docetaxel. Bevacizumab was added in cycles 3-6 for patients without a clinical complete response (cCR) after two cycles. After surgery, adjuvant therapy followed Swedish national guidelines. Details of the clinical trial have been previously reported by Kimbung et al (PMID: 28940389). Primary endpoints were early objective response and pathological complete response (pCR, absence of invasive tumor in breast and nodes; residual DCIS allowed). Recurrence-free survival (RFS) was a secondary endpoint. RNA was extracted from pre-treatment biopsies and surgical specimens, yielding 125 samples. Expression profiling was performed on the Illumina Human HT-12 v4.0 BeadChip (Illumina, San Diego, CA), as previously described (GSE87455) (PMID: 28940389). The clinical study and correlative analyses were approved by the Ethics Committee at Karolinska University Hospital (2007/1529–31/2). All patients provided written informed consent for participation in the trial and for associated translational research.

## Code availability

All custom scripts used in this study for single-cell ATAC-seq and RNA-seq processing, integration, and downstream analyses are available from the corresponding author upon reasonable request. scATAC-seq data analysis was performed using ArchR (v1.0.1), and transcriptional regulatory network inference was conducted with pySCENIC (v0.11.2). Single-cell RNA-seq data processing and visualization were carried out using Seurat (v4.3.0) and Scanpy (v1.9.1) in R and Python environments, respectively. All analysis scripts have been deposited at https://github.com/yizhesuncode/scATAC_pySCENIC_aNK and are publicly available under the MIT license.

## Data availability

All raw and processed single-cell ATAC-seq and RNA-seq data generated in this study are available from the corresponding author upon reasonable request. Publicly available datasets used in the analysis, including TCGA-HGSOC bulk RNA-seq were obtained via the cBio Cancer Genomics Portal. All custom analysis scripts have been deposited at GitHub: https://github.com/yizhesuncode/scATAC_pySCENIC_aNK and are publicly available under the MIT license.

## Results

### scATAC-seq analysis revealed distinct accessible chromatin landscapes in ovarian adaptive NK cells

To dissect the regulatory features of tumor-infiltrating aNK cells in ovarian cancer, we implemented a multilayer integrative analysis pipeline.

We re-analyzed a publicly available ovarian cancer dataset by integrating scATAC-seq and scRNA-seq to annotate NK cell subpopulations and identify chromatin-accessible regions, from which we reconstructed TF-gene regulatory networks. In parallel, Smart-seq2 and Smart-seq3 transcriptomic profiling was performed on sorted cells to characterize TF regulon activity at single-cell resolution for tumor-infiltrating NK cells. By combining the chromatin accessibility and transcriptional layers, we pinpointed shared TFs and inferred putative target genes specifically associated with the aNK cluster. This multi-omics workflow **(Figure 1A)** provided a framework to connect epigenetic potential with transcriptional output in defining functional regulators of aNK cell identity.

**Figure 1.**
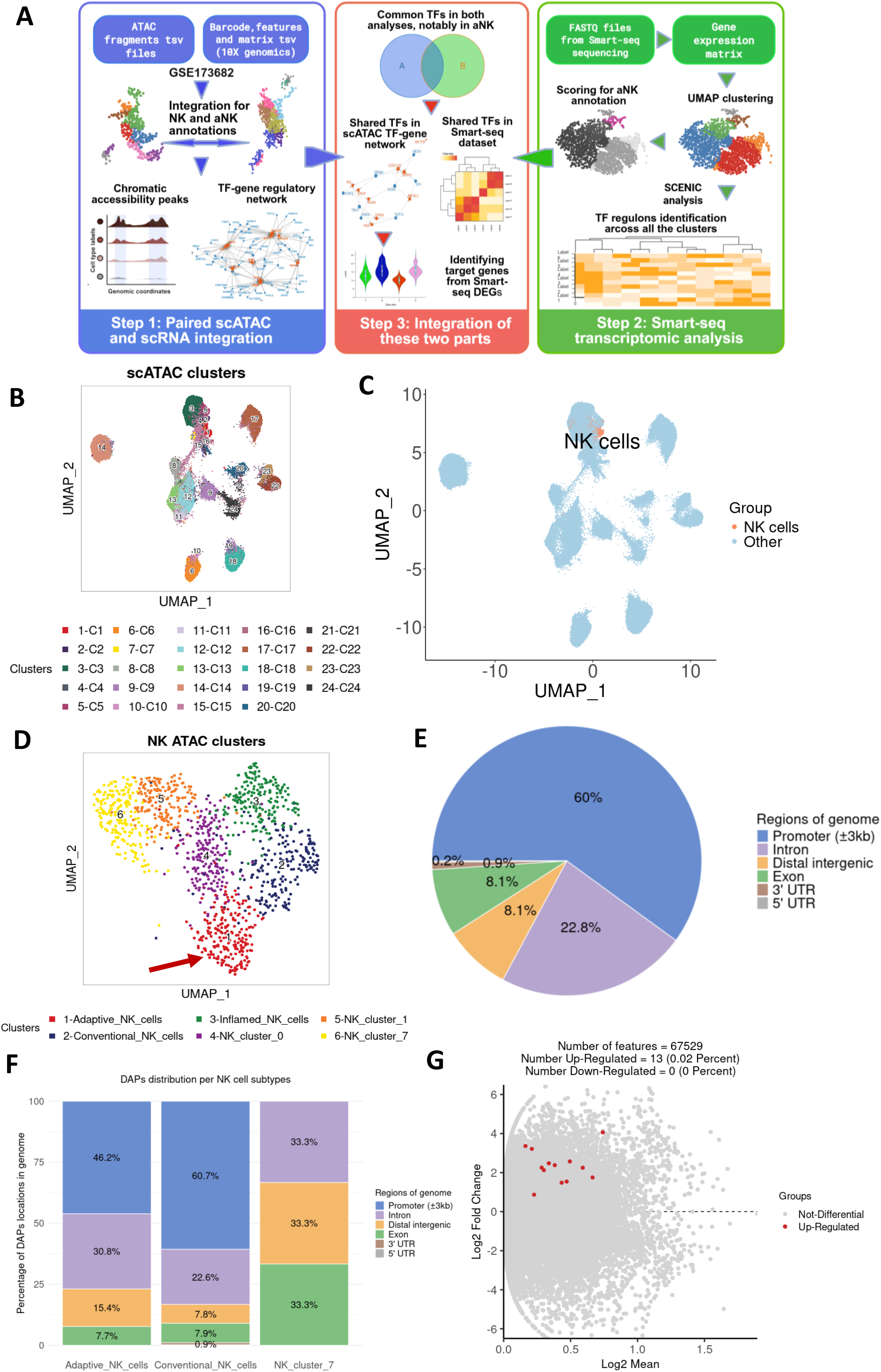
scATAC-seq identifies distinct chromatin accessibility peaks in adaptive NK (aNK) cells. **A,** Schematic workflow of the overall analysis **B.** UMAP visualization of scATAC-seq data showing all identified cell clusters. **C.** NK cell populations were annotated via integration with matched scRNA-seq data. **D.** Finalized UMAP plot displaying NK-specific clusters extracted from the complete dataset. **E.** Pie chart showing genomic distribution of peaks across promoter, intronic, intergenic, exonic, and UTR regions. **F**. Stacked bar plot showing the proportion of different genomic regions containing differentially accessible peaks (DAPs) in adaptive NK cells, conventional NK cells, and NK_cluster_7**. G**. Volcano map showing 13 chromatin peaks were found to be significantly enriched in the aNK cluster.

To test our approach, we first analyzed a publicly available multiomic dataset (GSE173682)^[33]^, which includes paired single-cell RNA sequencing (scRNA-seq) and scATAC-seq data from patients with gynecological malignancies. This dataset includes tumor samples from two anatomical sites, endometrium and ovary. To focus specifically on ovarian cancer, we included only tumor samples derived from ovarian sites, representing six patients (Patients 6–11).

Unsupervised clustering based on scATAC-seq profiles revealed 24 distinct clusters **(Figure 1B)**. Differential accessibility analysis (FDR ≤ 0.01 and Log₂FC ≥ 1), identified NK cell-associated genes including *CD69, KLRD1, KLRC1, KLRC2, NCR2, and NCR3*, predominantly present in clusters 1, 2, and 3 **(Supplementary Figure 1A)**. To refine transcriptomic annotation of NK cells, we performed unsupervised clustering on the scRNA-seq data from 6 ovarian tumors, identifying 31 transcriptomic clusters **(Supplementary Figure 1B)**. Cell types were annotated using the HPCA (Celldex) and BlueprintEncode reference datasets. Cells classified as NK cells in either reference were labeled accordingly and visualized on the UMAP projection **(Supplementary Figure 1C)**. Integration of scRNA-seq and scATAC-seq data enabled the transfer of these annotations, allowing precise identification of NK cells within the full scATAC-seq dataset **(Figure 1C)**.

Following NK cell annotation, we extracted the NK cell population from the complete scRNA-seq dataset. This subset was resolved into eight transcriptionally distinct subclusters **(Supplementary Figure 1D)**. To annotate these NK subpopulations, we established a reference framework comprising aNK cells, conventional NK (cNK) cells, inflamed NK cells, and terminal NK cells **(Supplementary Table 1)**. Specifically, the marker genes for aNK cells were derived from two sources. The primary source was bulk RNA sequencing data comparing late-mature aNK cells (CD3⁻ CD56^dim^ CD57⁺ NKG2C⁺) to early-mature cNK cells (CD3⁻ CD56^dim^ CD57⁻ NKG2C⁻) from human peripheral blood^[34, 35]^. Early mature cNK cells were selected as the reference population for two principal reasons: (1) their developmental distance from aNK cells enhances the detection of differentially expressed genes (DEGs), and (2) they provide a clear contrast for identifying terminal NK cells within our annotation strategy. The second source of aNK cell marker genes was transcriptomic data from NK cells infiltrating ovarian tumors ^[20]^. To further annotate inflamed NK cells and terminal NK cell subsets, we additionally incorporated signatures from NK cells isolated from bone marrow and peripheral blood^[36]^.

Using the aNK cell gene set derived from our own and others’ discoveries (**Supplementary Table 1)**, we calculated module scores across all clusters and identified cluster 4 as having the highest aNK signature **(Supplementary Figure 1E)**. Next, to annotate all of the NK cell clusters, we applied the scType^[27]^ function to score each NK cell and assign annotations accordingly **(Supplementary Table 2)**. If the annotation score for a given cluster was lower than one-quarter of the total number of cells in that cluster, we designated it as "undefined" and retained its original cluster label in the format "NK_cluster (original number)". Based on the annotation results, we defined cluster 4 as aNK cells, cluster 5 as cNK cells, and cluster 3 as inflamed NK cells **(Supplementary Figure 1F)**. Based on the scType cluster annotation, although cluster 7 exhibited a module score comparable to that of cluster 4, it was classified as transitional NK cells rather than aNK cells. Furthermore, cluster 7 was located far from cluster 4 in the embedding space, suggesting clear differences between the two. Consequently, cluster 4 was ultimately annotated as aNK cells.

We then applied a similar integration strategy as used for the scATAC–scRNA alignment to define NK cell subpopulations within the scATAC dataset. For scATAC NK cell clustering, the original clusters prior to quality control (QC) filtering are shown **(Supplementary Figure 1G)**. This multimodal integration approach enabled the identification of an aNK-enriched cluster among scATAC-derived NK cells. Low-quality clusters lacking consistent transcriptional correspondence to the scRNA-seq reference, suggestive of mixed or ambiguous cell identities, were excluded from downstream analyses. Specifically, C5 in the original clusters corresponded to both Adaptive_NK_cells and NK_cluster_0 to a similar extent, resulting in an ambiguous classification **(Supplementary Figure 1F)**. For C1, although it matched Adaptive_NK_cells, its match rate was lower than that of C8 and comparable to C5. Moreover, its UMAP position was clearly separated from C8, indicating distinct differences. Therefore, we retained C8 as aNK cells and excluded C1 **(Supplementary Figure 1H)**.

Following the exclusion of clusters C1 and C5, we re-ordered and finalized the clusters **(Figure 1D)**: C8 from the original clusters, classified as aNK cells, was designated as cluster 1; C7, classified as cNK cells, as cluster 2; and C4, classified as inflamed NK cells, as cluster 3.

Following the annotation of the aNK cell subset, we first profiled the global chromatin accessibility landscape of NK cell subsets and quantified the total number of peaks within each group **(Supplementary Figure 2A)**. We then identified differentially accessible peaks (DAPs) between NK cell subsets (FDR ≤ 0.1 and Log₂FC ≥ 0.25). Overall, promoter peaks accounted for 60% of all DAPs, followed by intronic (22.8%) and distal intergenic (8.1%) regions. Subset-specific analysis revealed that cNK cells had the highest proportion of promoter-associated DAPs (60.7%), whereas aNK cells exhibited a larger fraction of intronic DAPs (30.8%) and distal intergenic regions (aNK: 15.4 % vs. cNK: 7.8%). In contrast, NK_cluster_7 displayed a more balanced distribution of DAPs among promoter, intronic, and distal intergenic regions **(Figure 1E and 1F)**. These patterns suggest that cNK cells predominantly regulate gene expression through promoter accessibility, consistent with rapid activation of effector programs. In contrast, aNK cells exhibit greater accessibility at both intronic and distal intergenic regions, indicative of enhancer-driven regulation and long-range chromatin interactions that underpin the stable transcriptional programs associated with adaptive or memory-like functions.

Subsequently, we selected the top 13 peaks exhibiting the highest accessibility, as determined by inserted fragment counts, across all NK clusters **(Supplementary Figure 2B)**, with particular enrichment observed in the aNK cluster **(Figure 1G)**.

### The TF motif–gene regulatory network was identified in the aNK cluster

To further characterize the transcriptional regulatory landscape of aNK cells, we analyzed TF motif enrichment within the 13 most DAPs regions identified in scATAC-seq data. This analysis identified a diverse repertoire of enriched motifs within the 13 aNK DAP regions, with *DMRT2*, *ZSCAN4*, and *RUNX1* ranking among the most prevalent **(Figure 2A, 2B)**. Several TFs known to be involved in lymphocyte development and tissue residency, including *RUNX1*, *PRDM1*, and *ZNF683* were also enriched, supporting their potential involvement in orchestrating aNK cell differentiation, migration, and long-term persistence in the tumor microenvironment. Notably, *STAT2*, a mediator of immune activation, was also enriched, suggesting a transcriptional link between immune responsiveness and aNK cell identity. Additionally, multiple members of the *SOX* family, namely *SOX3, SOX5, SOX6,* and *SOX13*, were consistently enriched, highlighting a potential function in shaping or maintaining aNK cell feature.

**Figure 2.**
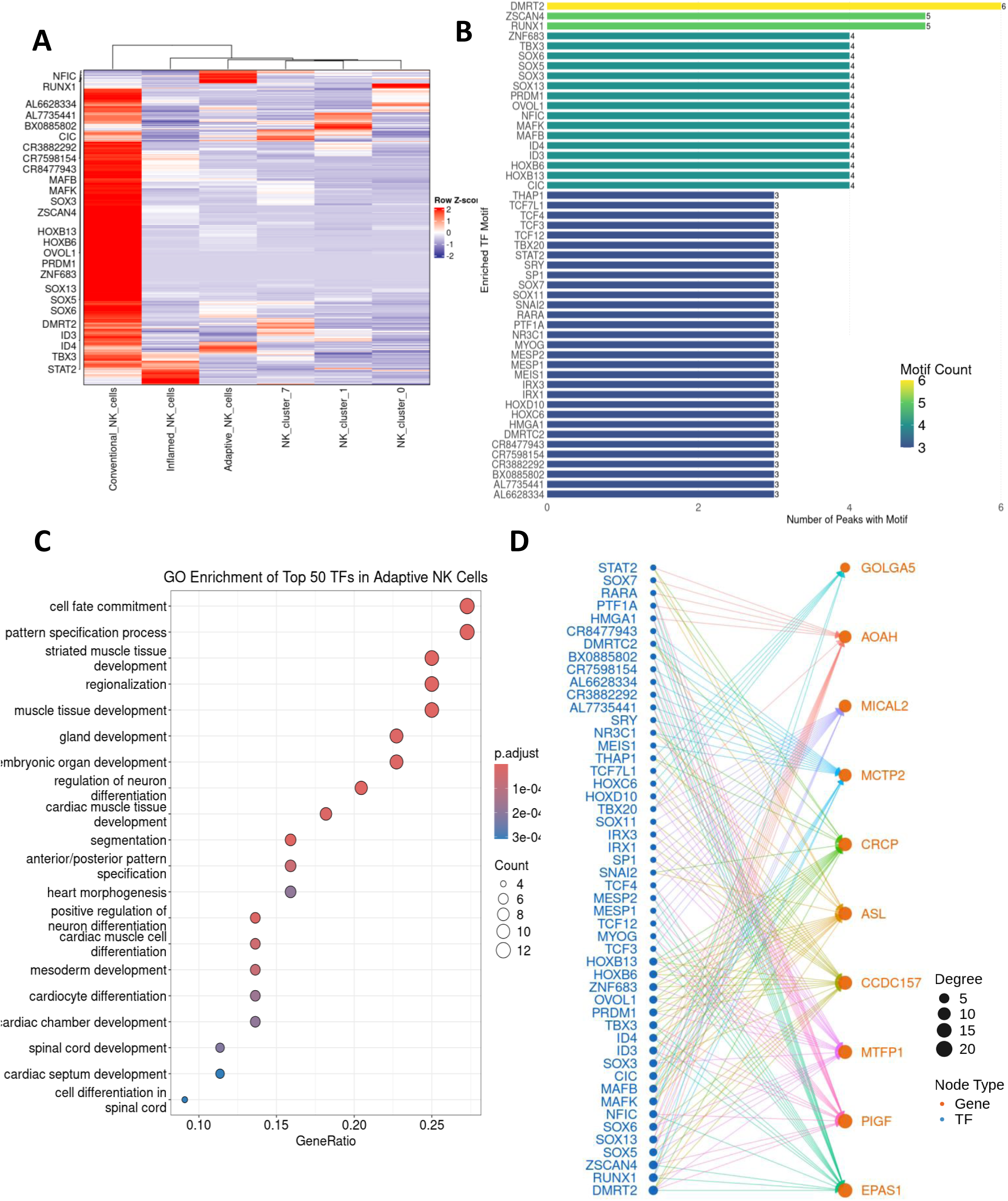
Transcription factor (TF) motif-gene regulatory network enriched in aNK cells. **A.** Row Z-scores of differentially accessible peaks (DARs) with selected adaptive NK cells related TFs are shown **B**. Bar plot showing the number of peaks containing each enriched TF motif. Bars are colored according to motif count, with warmer colors indicating higher counts. **C.** Gene Ontology (GO) enrichment analysis of the top 50 TFs in adaptive NK cells. **D.** The network shows the top 50 enriched transcription factors (TFs, blue) and their predicted target genes (orange)

Gene Ontology (GO) enrichment analysis of the top 50 TFs in aNK cells revealed significant associations with developmental processes such as cell fate commitment, pattern specification, and neuron differentiation-related pathways. These results suggest that aNK cell–specific TFs may engage regulatory networks influencing cell identity and functional specialization beyond conventional immune pathways. However, as the analysis was limited to the top 50 TFs, the breadth of enriched pathways may be underestimated **(Figure 2C)**.

To investigate the downstream impact of these regulatory regions, we next performed peak-to-gene correlation analysis by integrating scATAC-seq accessibility with transcriptomic profiles. This analysis identified ten genes, *CRCP, ASL, MCP2, AOAH, MTFP1, CCDC157, GOLGA5, MICAL2, PIGF*, and *EPAS1,* that exhibited significant positive correlation with DAPs for aNK cells accessibility **(Figure 2D)**. Several of these genes are associated with immunometabolic functions or cellular persistence, including *MTFP1*, involved in mitochondrial dynamics, and *EPAS1*, a hypoxia-responsive transcription factor implicated in tissue adaptation.

We then integrated TF motif and gene linkage results into a TF-gene regulatory network, revealing connections between enriched TFs and their putative downstream targets **(Figure 2D)**. In this network, transcription factor-gene interactions were inferred by combining motif enrichment from scATAC-seq peaks with peak-to-gene correlations from transcriptomic data. The resulting model highlights regulatory relationships between TFs such as *PRDM1*, *STAT2*, *RUNX1*, and SOX-family members and genes involved in immune signaling, mitochondrial function, and cellular persistence, including *CRCP, MTFP1*, and *EPAS1*, supporting a transcriptional program underpinning aNK cell identity within the tumor microenvironment.

### TF regulons were identified using SCENIC analysis based on Smart-seq transcriptomic data obtained from ovarian tumor-infiltrating NK cells

Building on the regulatory insights gained from scATAC-seq data, we next generated our own high-resolution, full-length transcriptomic dataset using Smart-seq, enabling sensitive detection of isoforms, modestly expressed genes, and transcriptional regulators in individual NK cells from human ovarian tumors. We integrated Smart-seq2 and Smart-seq3 datasets and applied batch-effect correction to harmonize the two platforms **(Supplementary Figure 3A).**

For Smart-seq2, tumor tissues from five patients were enzymatically dissociated, and both CD45⁻ and CD45⁺ cells were FACS-sorted into 384-well plates for single-cell sequencing, with the gating strategy shown in **Supplementary Figure 3B**. NK cells were annotated within the dataset. For Smart-seq3, tumor tissues from two patients were processed similarly, and CD3⁻CD56⁺ NK cells were FACS-sorted into 384-well plates for single-cell sequencing, as detailed in **Supplementary Figure 3C**. The overall workflow, including 7 patients (**Figure 3A**). Unsupervised clustering of the integrated resulting transcriptomic data revealed nine distinct NK cell clusters **(Figure 3B)**. Analysis of patient samples revealed variability in NK cell yield across donors. Smart-seq3 generated higher yields than Smart-seq2, primarily due to the direct NK cell sorting strategy (**Supplementary Figure 3D**).

**Figure 3.**
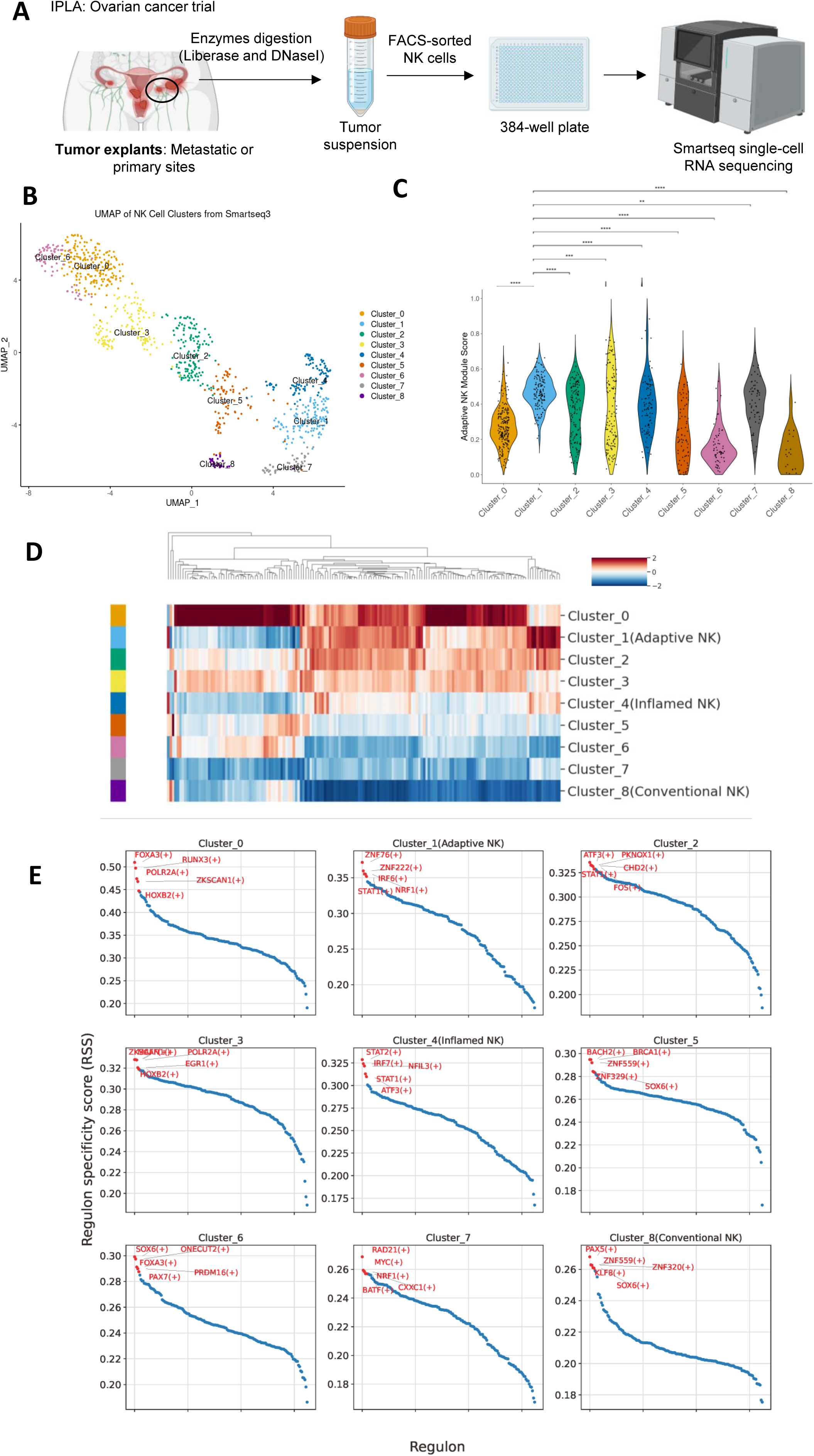
SCENIC identifies cluster-specific transcription factor (TF) regulon activities in aNK cells. **A.** Schematic overview of the Smart-seq workflow applied to tumor-infiltrating cells isolated from ovarian cancer patients.**B.** UMAP visualization of all identified NK cell clusters**. C**. Violin plot displaying the distribution of adaptive NK cell module scores across all clusters, with pairwise Mann-Whitney-Wilcoxon tests performed using Cluster 1 as the reference group. Statistical significance is denoted by: ** *p* < 0.01; *** *p* < 0.001; **** *p* < 0.0001. **D.** Heatmap displaying the regulon specificity scores (RSS) for all identified TF regulons (163) across NK clusters, highlighting their cluster-specific activity. **E.** Ranking plots showing the top 5 TF regulons with the highest

To align our data with previous annotations, we subsequently applied the same annotation reference to identify aNK cells within this dataset. Cluster 1 exhibited a markedly high aNK cell module score, consistent with a transcriptionally distinct aNK subset **(Figure 3C** and **Supplementary Table 3)**. This cluster was therefore classified as the aNK cell population within the Smart-seq dataset. In addition to cluster 1, cluster 8 was classified as cNK cells, while cluster 4 was identified as inflamed NK cells. Examination of the aNK cell cluster (Cluster_1) revealed contributions from multiple patients, with the majority originating from Smart-seq3_ov1 and additional representation from Smart-seq3_ov2 and Smart-seq2_ov1 (Supplementary Figure 3E). These observations indicate that aNK cells are reproducibly captured across both Smart-seq2 and Smart-seq3 platforms, although their relative abundance exhibits inter-patient variability.

To further investigate transcriptional regulators that may drive the aNK cell state, we applied the SCENIC workflow using the pySCENIC implementation, which leverages transcription factor-target gene co-expression and motif enrichment to infer regulatory modules (regulons). This analysis ^[37]^ yielded cluster-specific regulon specificity scores (RSS) across all nine NK cell clusters **(Figure 3D)**. Cluster 1, which had been previously annotated as the aNK cluster, showed marked enrichment for the regulons *ZNF76, ZNF222, IRF6, STAT1, and NRF1***(Figure 3E)**. These TFs represent strong candidates for orchestrating the transcriptional program underpinning aNK cell identity, function, and potential persistence in the TME.

Importantly, this regulatory inference was derived directly from our primary tumor-resident NK cell data, providing novel insights beyond those attainable from public datasets. The use of Smart-seq technology enabled precise mapping of full-length transcripts, improving the resolution of TF regulon activity and enhancing confidence in our identification of key regulatory programs specific to aNK cells in ovarian cancer.

### Integrative analysis of transcriptomic profiles and chromatin accessibility data identified PRDM1 and STAT2 as key transcriptional regulators for aNK cluster

To identify key TFs driving the aNK cell state, we performed a cross-modality integration of chromatin accessibility and transcriptomic regulatory activity. Specifically, we intersected TF regulons inferred by pySCENIC from our Smart-seq3 tumor-infiltrating NK cell intersected the TF regulons inferred by pySCENIC with the top 50 most enriched TFs from the aNK and cNK clusters based on scATAC-seq data. The pySCENIC analysis identified 163 regulons with differential activity (DA) across all NK cell clusters. This integrative approach revealed 21 overlapping TFs shared between SCENIC-inferred regulons and those enriched in scATAC- defined aNK or cNK clusters. Among these, 5 TFs were uniquely associated with the aNK cluster, 13 with the cNK cluster, and 3 TFs, were shared across both datasets **(Figure 4A)**.

**Figure 4.**
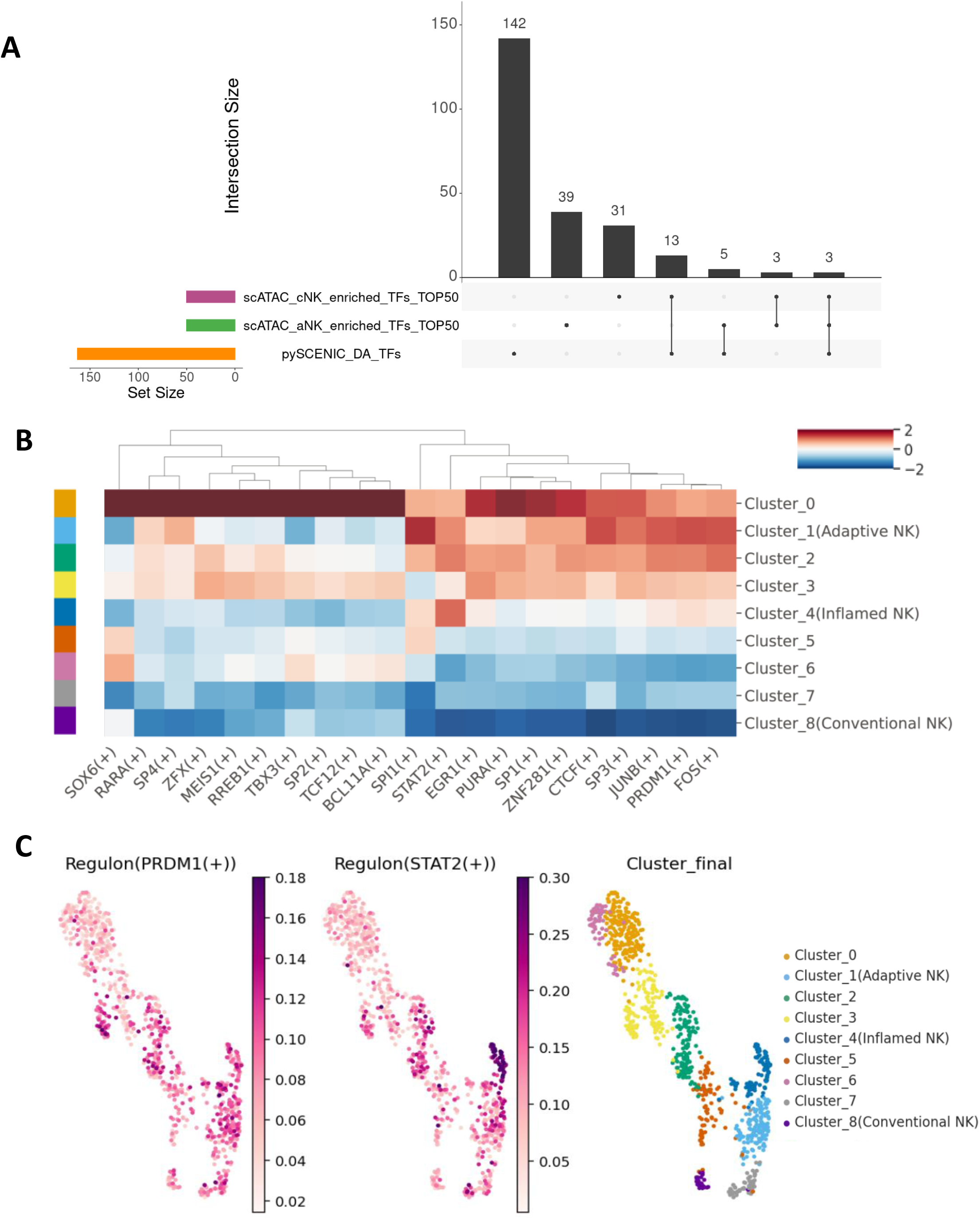
PRDM1 and STAT2 are identified as key regulators for aNK cluster through integrative analysis of transcriptomic and chromatin accessibility data. **A.** Upset plot showing the overlap of 21 transcription factors identified by pySCENIC (Smart-seq3 data) and those associated with adaptive (aNK) and conventional NK (cNK) clusters from scATAC-seq analysis. **B.** Heatmap illustrating the regulon specificity scores (RSS) of the 21 overlapping TF regulons across NK cell clusters, highlighting cluster-specific regulatory activity. **C.** UMAP plot showing the spatial distribution of two shared regulons, PRDM1 and STAT2, both identified in the aNK cluster (Cluster 1) by pySCENIC and in aNK cluster by scATAC-seq.

We then evaluated the RSS of the 8 overlapping TFs between the scATAC-defined aNK cluster and SCENIC-inferred regulons using the pySCENIC results. Among them, *PRDM1* and *STAT2* exhibited the most distinct activity in cluster 1 (the aNK cluster) from Smart-seq sequencing **(Figure 3B)** and were also identified in the aNK cluster based on scATAC-seq data **(Figure 4B, Supplementary Figure 4A and Figure 4B)**, highlighting their potential regulatory roles in aNK cells. Furthermore, we visualized the RSS of *PRDM1* and *STAT2* on UMAP projection, revealing markedly elevated regulon activity for both transcription factors **(Figure 4C)**. These findings, based on full-length single-cell transcriptomics, independently confirm the key regulatory axes inferred from chromatin accessibility and underscore the value of integrative multiomic analysis in defining NK cell states within the TME.

### scATAC regulatory analysis identified *CRCP* and *MTFP1* as potential downstream targets of PRDM1 and STAT2

Having identified PRDM1 and STAT2 as candidate master regulators of the aNK cell transcription program, we next aimed to determine their downstream target genes. To this end, we first constructed a TF-target gene regulatory network based on the 21 TFs identified in **Figure 4A**. This analysis revealed nine putative target genes, including *ASL, CRCP, AOAH, EPAS1, PIGF, CCDC157, MTFP1, MICAL2, and MCTP2* **(Figure 5A)**. We next evaluated the expression of the candidate target genes in Smart-seq-based scRNA-seq data and found that both *CRCP* and *MTFP1* were among the DEGs. Notably, their expression levels were significantly elevated in cluster 1, corresponding to the aNK cell population, compared to other clusters **(Figure 5B and 5D)**. *CRCP* was predicted to be co-regulated by both PRDM1 and STAT2, while *MTFP1* appeared to be primarily driven by PRDM1 **(Figure 5A)**.

**Figure 5.**
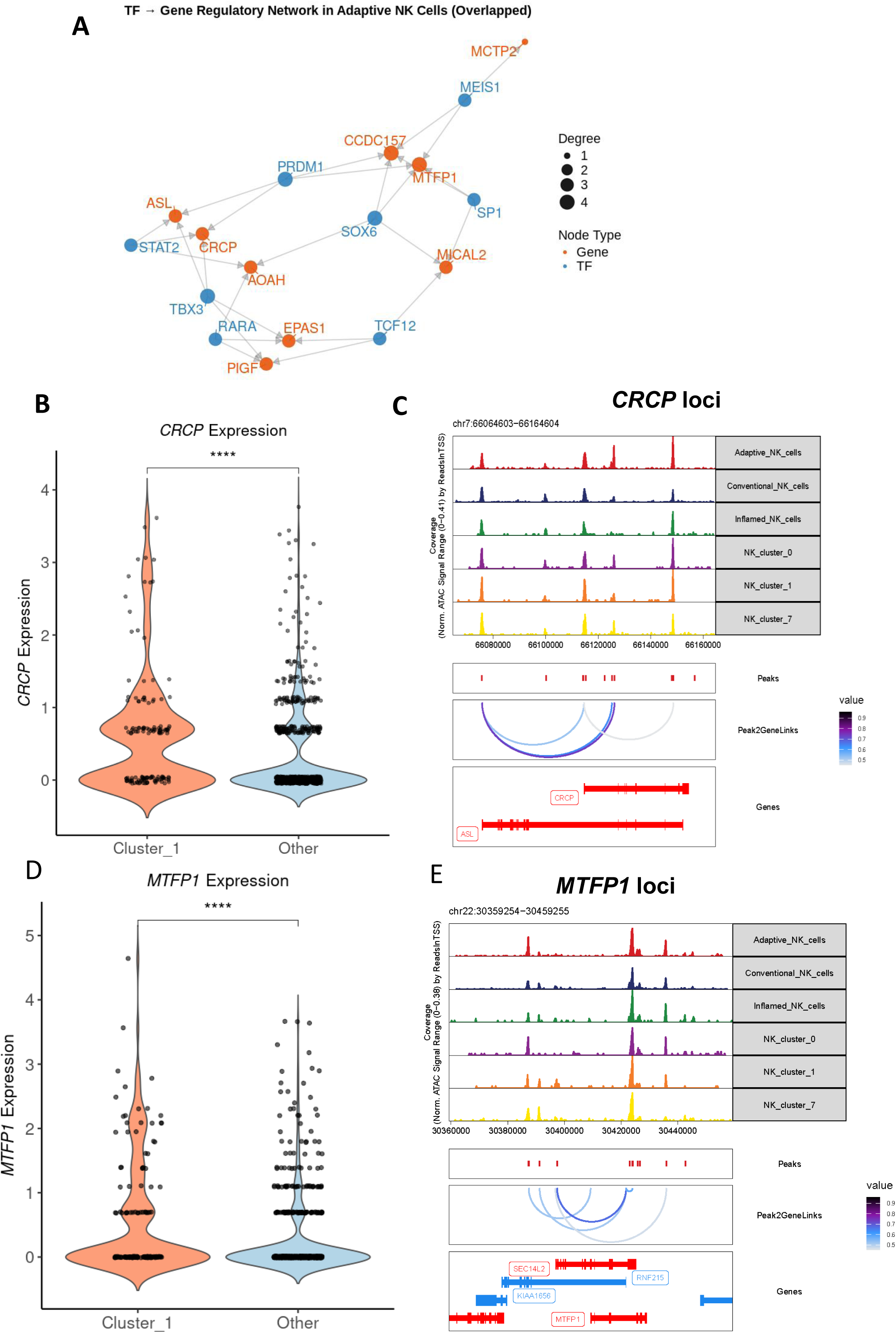
*CRCP* and *MTFP1* are identified as potential downstream targets of PRDM1 and STAT2 through integrated regulatory analysis. **A.** Transcriptional regulatory network of the shared TFs identified in Figure 4A within the aNK cluster-8 TFs, revealing 10 putative target genes based on peak-to-gene correlations. **B.** Violin plot showing high expression of the two TFs(PRDM1 and STAT2) -regulated gene *CRCP* in the aNK cluster (Cluster 1). Mann-Whitney-Wilcoxon test was performed. Statistical significance is denoted by: **** *p* < 0.0001. **C.** Peak-to-gene correlation map at the *CRCP* locus, showing enriched chromatin accessibility and predicted TF-binding motifs specific to the aNK cluster. **D.** Violin plot showing high expression of another PRDM1- and STAT2-regulated gene, *MTFP1*, in the aNK cluster (Cluster 1). Mann-Whitney-Wilcoxon test was performed. Statistical significance is denoted by: **** *p* < 0.0001. **E.** Peak-to-gene correlation map at the *MTFP1* locus, demonstrating strong chromatin accessibility and regulatory motif enrichment in the aNK cluster.

To validate the regulatory potential of these genes, we examined their chromatin accessibility and peak-to-gene connectivity using scATAC-seq data. The genomic loci of *CRCP* and *MTFP1* exhibited clear accessibility in the aNK cluster, with strong peak-to-gene linkages inferred through co-accessibility analysis **(Figure 5C and 5E)**. Importantly, these regulatory regions showed strong peak-to-gene linkage scores (>0.7), suggesting active enhancer-promoter communication. The accessibility of these sites was both cluster-specific and enriched for PRDM1 and STAT2 binding motifs, providing further evidence of direct transcriptional regulation.

Together, these multiomic results support a model in which PRDM1 and STAT2 coordinately regulate the expression of key effector genes, including *CRCP* and *MTFP1*, through aNK- specific accessible chromatin region. This regulatory axis likely contributes to the persistence and functional reprogramming of aNK cells within the TME.

### External validation confirms aNK-associated regulatory program with prognostic relevance across cancer types

To validate the broader relevance of the transcriptional targets identified in our single-cell analysis, we performed independent in silico validation using publicly available bulk RNA-seq datasets from ovarian and breast cancer cohorts. Using the TIMER2.0 platform and applying the CIBERSORT-ABS algorithm ^[38, 39]^, we found a significant positive correlation between *MTFP1*expression and NK cell infiltration in ovarian cancer **(Figure 6A)**. Similarly, using the QUANTISEQ deconvolution method^[40]^, *CRCP* expression also correlated positively with NK cell abundance **(Figure 6B)**. These results suggest that both genes are transcriptionally active in NK-rich tumors and potentially reflect an aNK-related immune activity.

**Figure 6.**
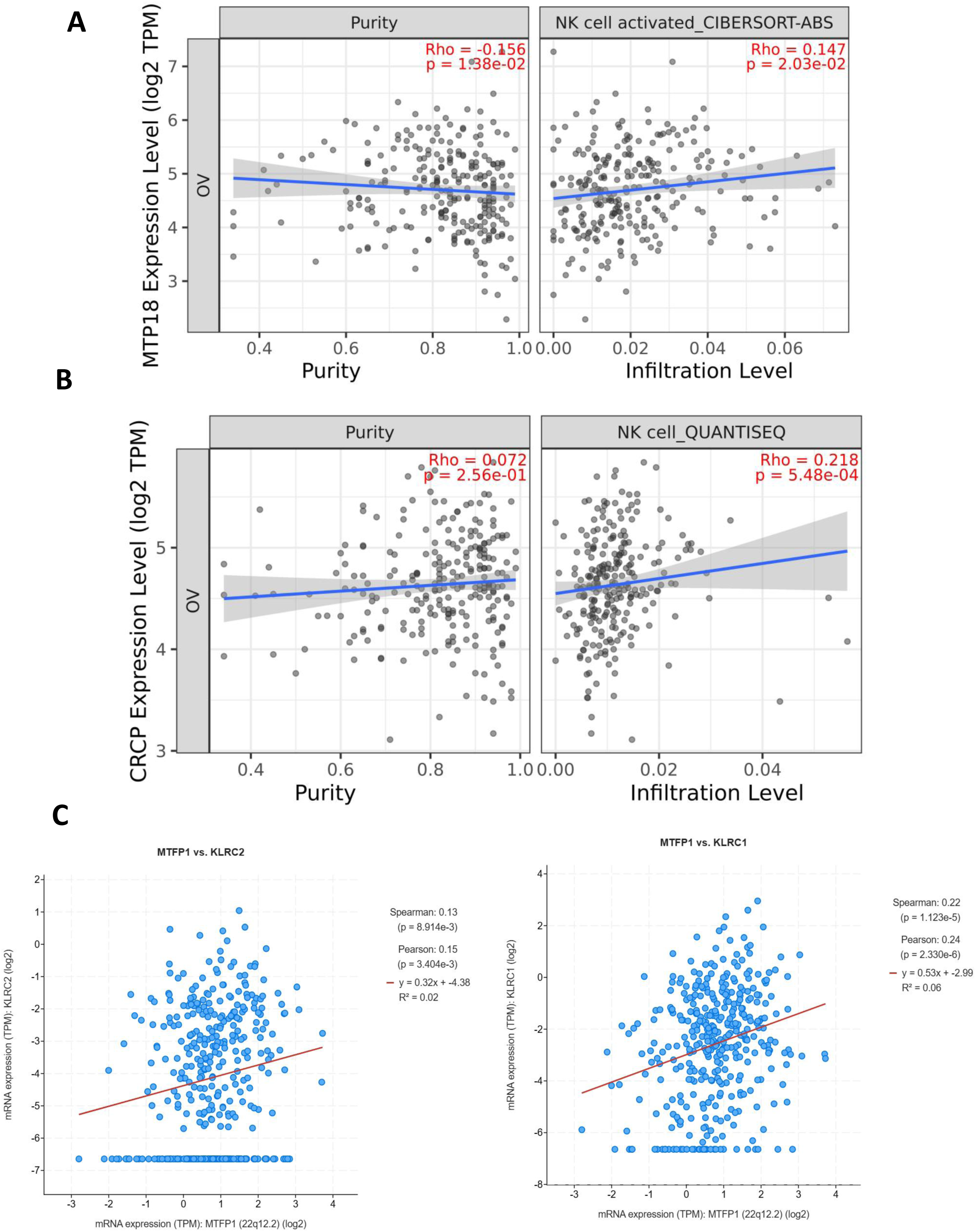
Independent validation of selected target genes. **A.** Correlation map illustrating that expression of *MTFP1* (also known as MTP18) is positively correlated with NK cell infiltration, as determined by CIBERSORT-ABS in ovarian cancer samples. **B.** Correlation map illustrating that expression of *CRCP* shows a positive correlation with NK cell infiltration, as estimated by QUANTISEQ in ovarian tumors**. C.** *MTFP1* expression shows a positive correlation with *KLRC2* and *KLRC1* in HGSOC samples, based on data from the cBio Cancer Genomics Portal.

To assess whether MTFP1 expression reflects broader aNK cell programming, we next analyzed bulk RNA-seq data from 602 HGSOC samples in TCGA via cBio Cancer Genomics Portal^[41]^. We observed that *MTFP1* expression was significantly positively correlated with *KLRC2* and *KLRC*1, which are canonical markers of CMV-induced aNK cells and cytokine-induced memory NK cells, respectively **(Figure 6C)**^[42]^. In contrast, *CRCP* expression showed no correlation with *KLRC2* or *KLRC1* **(Supplementary Figure 5A)**, indicating that *CRCP* may not be directly associated with canonical aNK cell signatures and could be regulated independently of *KLRC2/KLRC1-*related pathways.

To determine whether the aNK-associated transcription factors identified in ovarian cancer are relevant across distinct tumor types, we examined PRDM1 and STAT2 expression in a clinically independent setting using data from the PROMIX (Preoperative Treatment of Breast Cancer with a Combination of Epirubicin, Docetaxel and Bevacizumab) clinical trial^[43]^. This cohort of HER2-negative breast cancer patients receiving neoadjuvant therapy provided a valuable context for testing the generalizability of our findings. In this cohort, *PRDM1* and *STAT2* expression levels were positively correlated with aNK cell gene signatures **(Figure 7A)**, reinforcing the broader applicability of this regulatory program. Moreover, high expression of either TF was associated with improved recurrence-free survival (RFS) in this cohort **(Figure 7B)**, suggesting that the regulatory state we defined for tumor-infiltrating aNK cells carries prognostic relevance in breast cancer. These results extend the importance of PRDM1 and STAT2 beyond ovarian cancer and support their potential as immunogenomic biomarkers with clinical utility.

**Figure 7.**
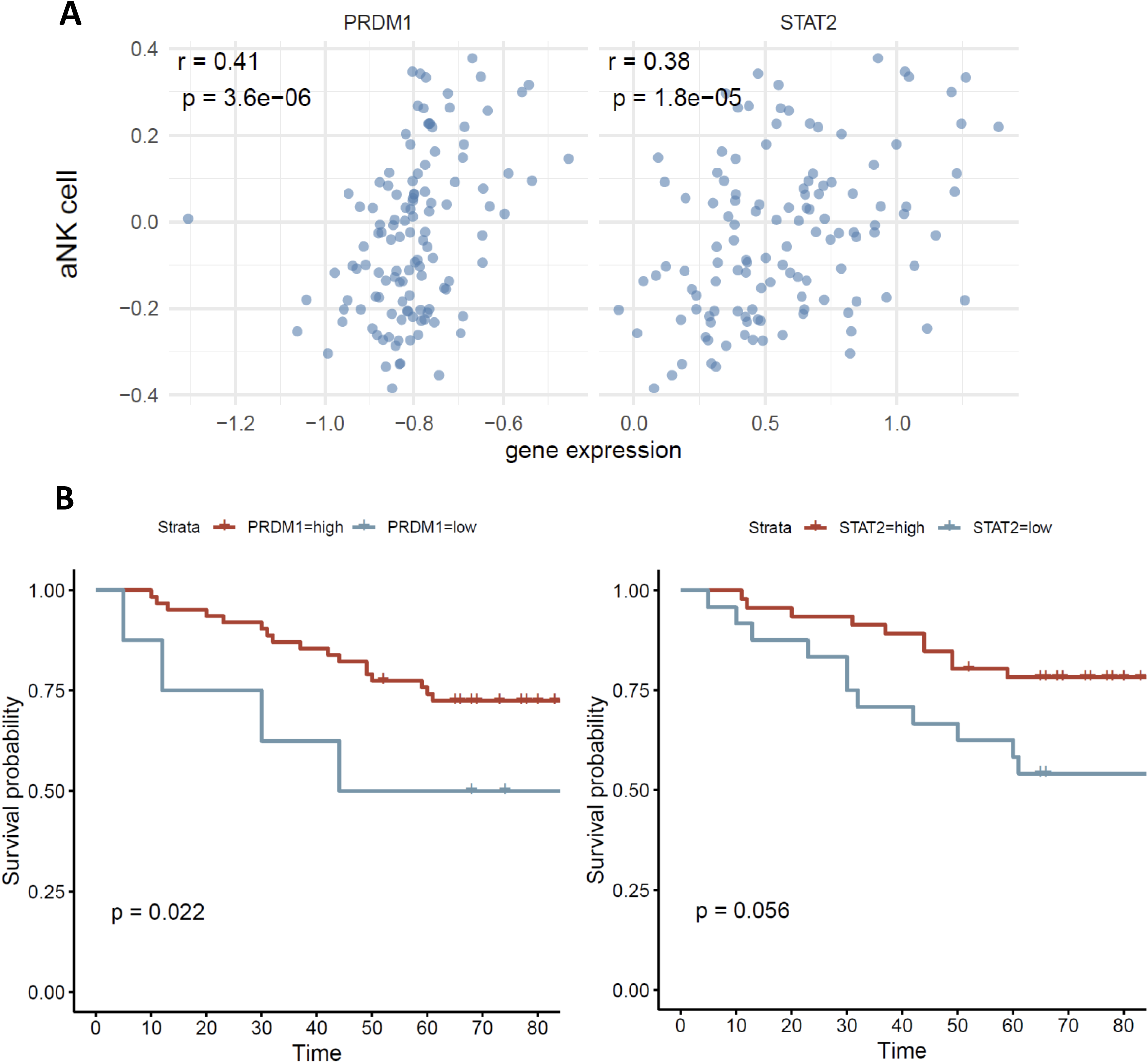
Independent validation of *PRDM1* and *STAT2* as aNK cell–associated transcription factors in the PROMIX clinical trial. **A.** Scatter plots showing the positive correlation between gene expression of *PRDM1* (left) and *STAT2* (right) with aNK cell scores in breast cancer tumor samples from the PROMIX clinical trial. Correlation coefficients (r) and associated p-values are indicated. **B.** Kaplan–Meier survival analysis based on *PRDM1* (left) and *STAT2* (right) expression levels in the PROMIX cohort. Patients with high PRDM1 expression show significantly better survival (p = 0.022), while high STAT2 expression is associated with a trend toward improved survival (p = 0.056).

Together with our earlier analyses, these findings offer cross-cohort, cross-tumor validation of an aNK-like transcriptional state, implicating PRDM1 and STAT2 as key regulators, and supporting MTFP1 and CRCP as downstream effectors associated with immune infiltration and favorable clinical outcomes.

## Discussion

The discovery of aNK cells within solid tumors has challenged the traditional view that NK cells are strictly short-lived, innate effectors lacking memory. While previous studies have described memory-like NK populations following viral infection, particularly HCMV, their existence and functional significance in the TME are only beginning to be understood. Our study expands this emerging paradigm by uncovering a distinct, multiomic transcriptional and epigenetic program that defines tumor-associated aNK cells in ovarian cancer, with direct implications for immunotherapeutic strategies.

Although CMV-driven aNK cells have been extensively characterized, their counterparts within the tumor microenvironment remain insufficiently studied. A major challenge lies in the precise definition of tumor-infiltrating aNK cells. To date, most investigations into aNK biology still rely heavily on several significant genetic markers established in the CMV context such as NKG2C, NKG2A, and FcεRγ,. However, this approach is contentious, as virus infection and the tumor microenvironment differ fundamentally despite both involving antigen presentation and stimulation of aNK cells. Consequently, within tumor-infiltrating NK populations, single featured genes are not sufficient to define aNK cells. Instead, a broader gene signature, originally identified in peripheral aNK cells, is required for accurate characterization ^[14]^. Therefore, in this study, we defined tumor-infiltrating aNK cells using a comprehensive gene signature derived from the DEGs that distinguish peripheral aNK cells from other NK subsets, enabling a more accurate identification of tumor-infiltrating aNK cells beyond reliance on a limited set of featured genes. One of the key findings of this work is the identification of PRDM1 and STAT2 as potential regulators of the aNK cell state within tumors. Although both transcription factors have known roles in immune differentiation, PRDM1 in plasma cell maturation, memory formation, exhaustion regulation, and T/NKT cell functionality, exhaustion^[44–47]^. Moreover, STAT2 plays role in type I and II interferon signaling and early immune memory programming^[18, 48, 49]^. However, their combined involvement in orchestrating a memory-like NK cell state in solid tumors has not previously been reported. Their concurrent identification in both chromatin accessibility (scATAC-seq) and transcriptomic (pySCENIC) analyses strengthens their candidacy as central regulators and highlights the value of integrating single-cell multiomic approaches to dissect immune plasticity.

Importantly, our study identifies CRCP and MTFP1 as putative downstream targets of the PRDM1-STAT2 regulatory axis in aNK cells. Although none of these genes have been clearly linked to NK cell biology, both exhibited selective upregulation and increased chromatin accessibility within the aNK cell population, and their expression correlated positively with NK cell infiltration in tumors, suggesting potential roles in aNK cell persistence or functional adaptation within the TME. CRCP, a component of the Calcitonin Gene-Related Peptide (**CGRP**) receptor complex, may modulate NK cell responsiveness to external cues such as cytokines or neuroimmune signals, potentially contributing to tissue retention and plasticity^[50, 51]^. MTFP1, a key regulator of mitochondrial fission recently shown to drive mitochondrial genetic takeover, maintains the inner membrane integrity^[52, 53]^, may support metabolic reprogramming and longevity in aNK cells, features critical for immune recall and sustained anti-tumor function.

In conclusion, unlike virus-induced aNK cells, which have been relatively well characterized, the environmental cues driving NK cell adaptation in tumors remain largely elusive. Our findings suggest that chronic exposure to tumor-derived cytokines, metabolic stress, and altered antigen landscapes may epigenetically imprint NK cells with a memory-like phenotype, positioning them as long-lived, tissue-adapted effectors with enhanced recall potential.

Clinically, we demonstrate that high expression of PRDM1 and STAT2, identified independently in our data, correlates with the aNK gene signature and improved overall survival in HER2-negative breast cancer patients (PROMIX trial), extending the relevance of our findings beyond ovarian cancer. This cross-tumor validation underscores the prognostic potential of aNK transcriptional programs and supports the utility of PRDM1 and STAT2 as immunogenomic biomarkers.

Methodologically, this study demonstrates the utility of integrating high-sensitivity Smart-seq3 profiling with publicly available multiomic datasets to dissect rare and functionally specialized immune subsets. Smart-seq3 enables near full-length transcript coverage with UMI-based quantification, making it particularly suitable for low-abundance populations such as tumor-infiltrating adaptive NK cells. By combining regulon inference from pySCENIC with motif accessibility and peak-to-gene correlation derived from scATAC-seq, we were able to prioritize transcriptional regulators with multi-layered support. This integrative approach improves confidence in regulator-target relationships that may be difficult to resolve using transcriptomic or epigenomic data alone.

Nonetheless, there are several limitations to consider. First, the regulatory relationships inferred in this study remain correlative and computational; direct validation of PRDM1 and STAT2 binding to *CRCP* and *MTFP1* loci will be essential to confirm direct gene regulation. Second, while our classification of aNK cells is based on well-established gene signatures, the heterogeneity within NK cell subsets may blur functional boundaries, and future studies with longitudinal or perturbation data are needed to clarify lineage trajectories. Finally, although we observed associations between CRCP/MTFP1 expression and NK infiltration in tumors, their functional relevance *in vitro* and *in vivo*, such as effects on persistence, cytotoxicity, or recall, remains to be evaluated.

Taken together, our results propose a new framework for understanding aNK cell function in solid tumors. We show that tumor-infiltrating aNK cells are governed by a transcriptional program likely orchestrated by PRDM1 and STAT2 and mediated by novel effectors such as CRCP and MTFP1. These insights have direct implications for biomarker discovery, therapeutic targeting, and the broader understanding of innate immune memory in cancer.

## Acknowledgments

The authors would like to acknowledge the contributions of the Flow Cytometry Core Facility and the Single Cell Core Facility Flemingsberg (SICOF), financed by the Infrastructure Board at Karolinska Institutet, for providing instrumentation for cell sorting and technical services.

## Author contributions

Y.S. and D.S. conceived the original research idea and study design. Y.S., M.W., S.K., K.W., S.Kh., and O.G. conducted and analyzed experimental data. Y.S., M.W., S.K., S.Kh., R.D and D.S. contributed to the interpretation of the data. S.S. and K.L., Y.S., and O.G., recruited study participants and collected essential human samples. D.S., K.L., R.D., T.F., M.E., and SS provided study materials, reagents, instrumentation, formal analysis or other analysis tools, interpretation, validation, and supervision. D.S., Y.S., M.W., wrote the manuscript. All authors contributed to the final review and editing.

