## Supplementary material for "Single-Cell Integration of Chromatin Accessibility and Transcriptomics Reveals Regulatory Networks in Ovarian Tumor-Infiltrating Adaptive NK Cells": Supp figs and tables

### Supplementary Figure 1

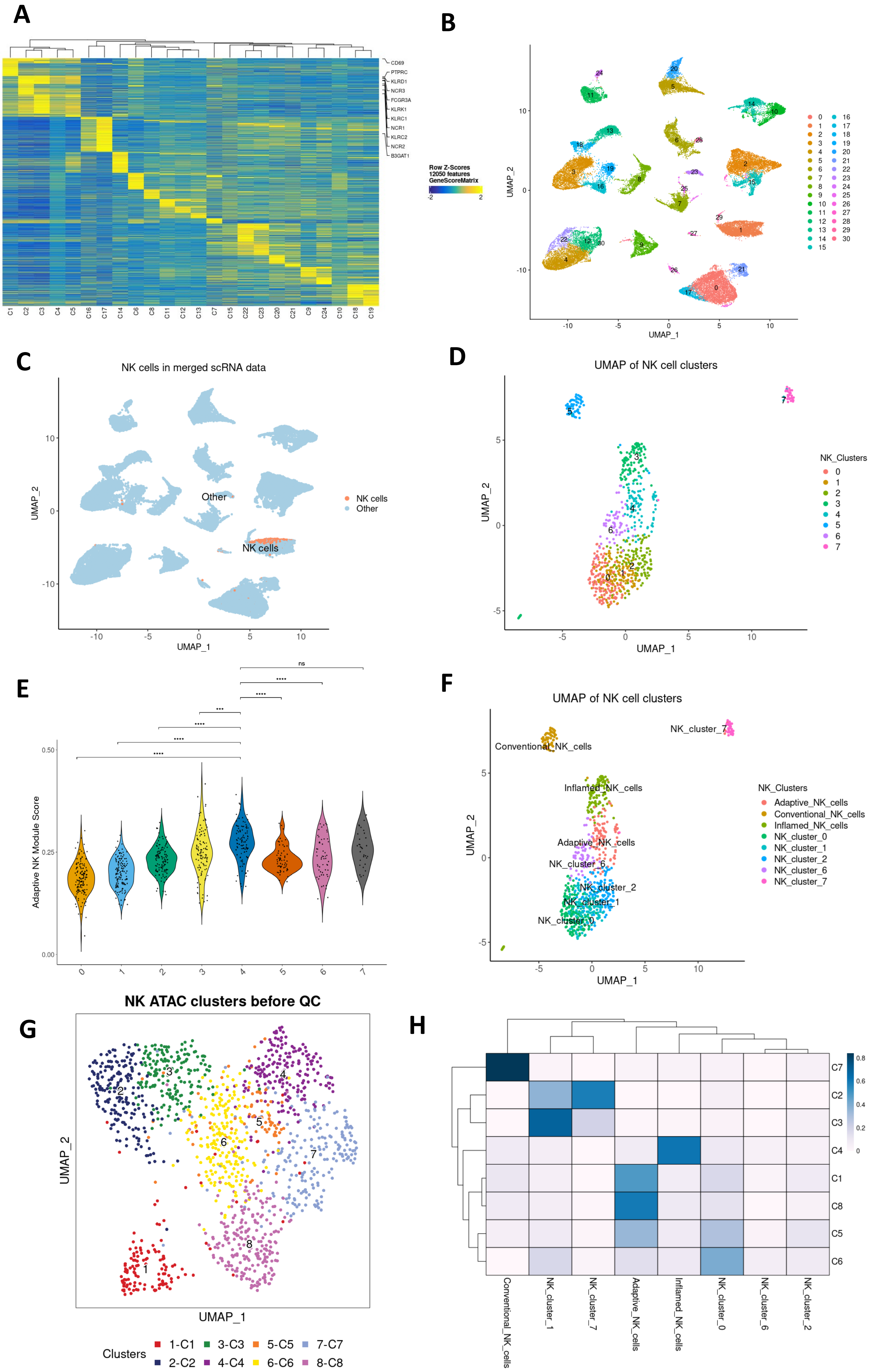


**Supplementary Figure 1. NK cells and aNK cells are distinguished from scRNA sequencing data. A.** Heatmap showing the expression of canonical NK cell marker genes across all ATAC-derived clusters.**B.** UMAP visualization of clusters from scATAC-matched scRNA-seq data. **C.** Annotation of NK cells among the scRNA-seq clusters. **D.** UMAP plot displaying NK cell subclusters derived from the NK population identified in panel C. **E.** Violin plot illustrating the distribution of aNK cell module scores across NK subclusters defined in panel D. Pairwise Mann–Whitney–Wilcoxon tests were performed using Cluster 1 as the reference. Statistical significance is indicated as follows: $**p < 0.01$; $***p < 0.001$; $****p < 0.0001$.**F.** Annotation of NK subclusters based on transcriptional profiles. **G.** UMAP plot displaying NK-specific ATAC clusters before the quality control (QC). **H.** Heatmap showing the correlation between scATAC-defined NK subclusters and scRNA-defined NK subclusters based on multimodal integration.

**Supplementary Figure 2**

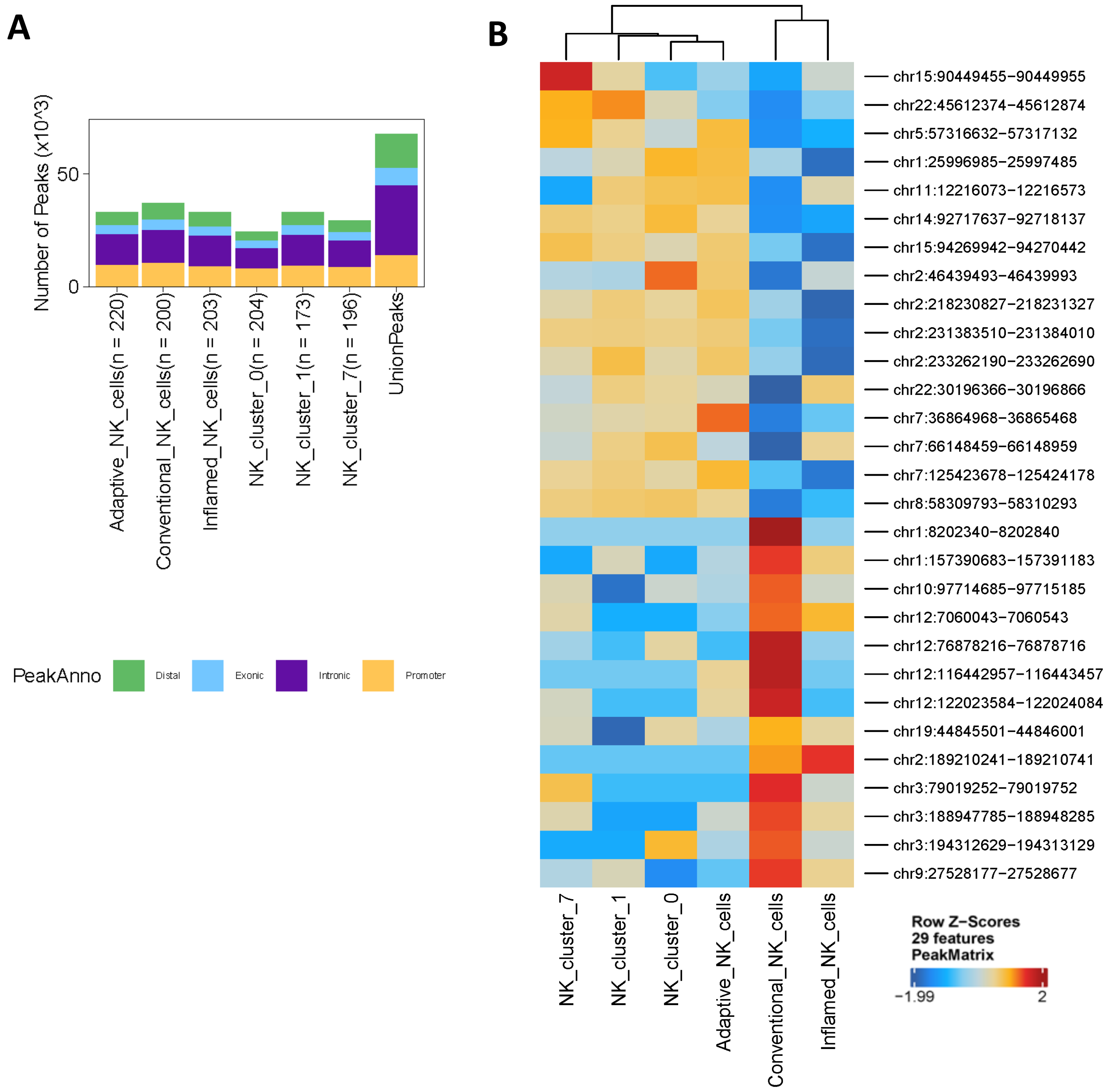


**Supplementary Figure 2. Peaks were identified for aNK cluster from scATAC data. A.** Bar plot showing the number and genomic annotation of accessible chromatin peaks identified in each NK cell subcluster, including adaptive NK cells, conventional NK cells, inflamed NK cells, and other NK clusters. Peaks were annotated based on their genomic location as distal, exonic, intronic, or promoter regions. A union peak set across all NK subsets is shown on the right for reference. **B.**Heatmap illustrating the accessibility patterns of the top 13 aNK-enriched peaks across all NK clusters.

### Supplementary Figure 3

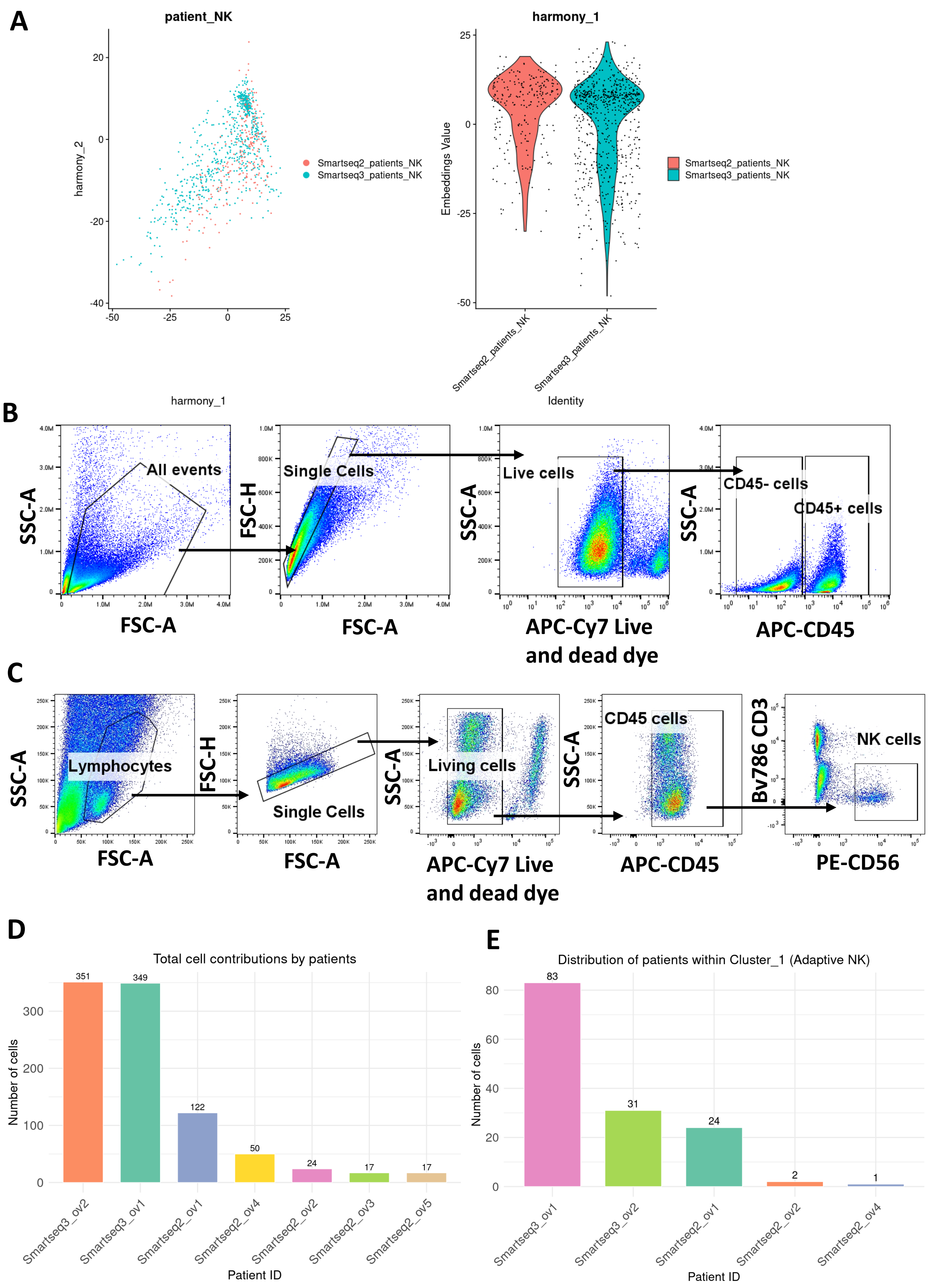


**Supplementary Figure 3. Integration of Smart-seq2 and Smart-seq3 datasets, corresponding gating strategies and distributions of samples.**

**A.** Harmony-corrected UMAP of NK cells from Smart-seq2 (red) and Smart-seq3 (cyan), showing alignment across platforms. Violin plots of Harmony_1 confirm comparable distributions after batch correction.

**B.** Representative flow cytometry gating for Smart-seq2 cells. All events were gated by FSC/SSC, singlets by FSC-H/FSC-A, and live cells by APC-Cy7 viability dye. $CD45^{+}$ leukocytes and $CD45^{-}$ cells were selected.

**C.** Representative flow cytometry gating for Smart-seq3 NK cells. Lymphocytes were gated by FSC/SSC, singlets by FSC-H/FSC-A, and live cells by APC-Cy7 viability dye. $CD45^{+}$ leukocytes were selected, and NK cells defined as $CD3^{-}CD56^{+}$ using Bv786-CD3 and PE-CD56.

**D.** Total NK cell numbers contributed by each patient, showing variation in overall cell yield between Smart-seq2 and Smart-seq3 datasets.

**E.** Distribution of patient-derived NK cells within Cluster_1 (Adaptive NK), highlighting the relative contribution of individual patients to this subset

### Supplementary Figure 4

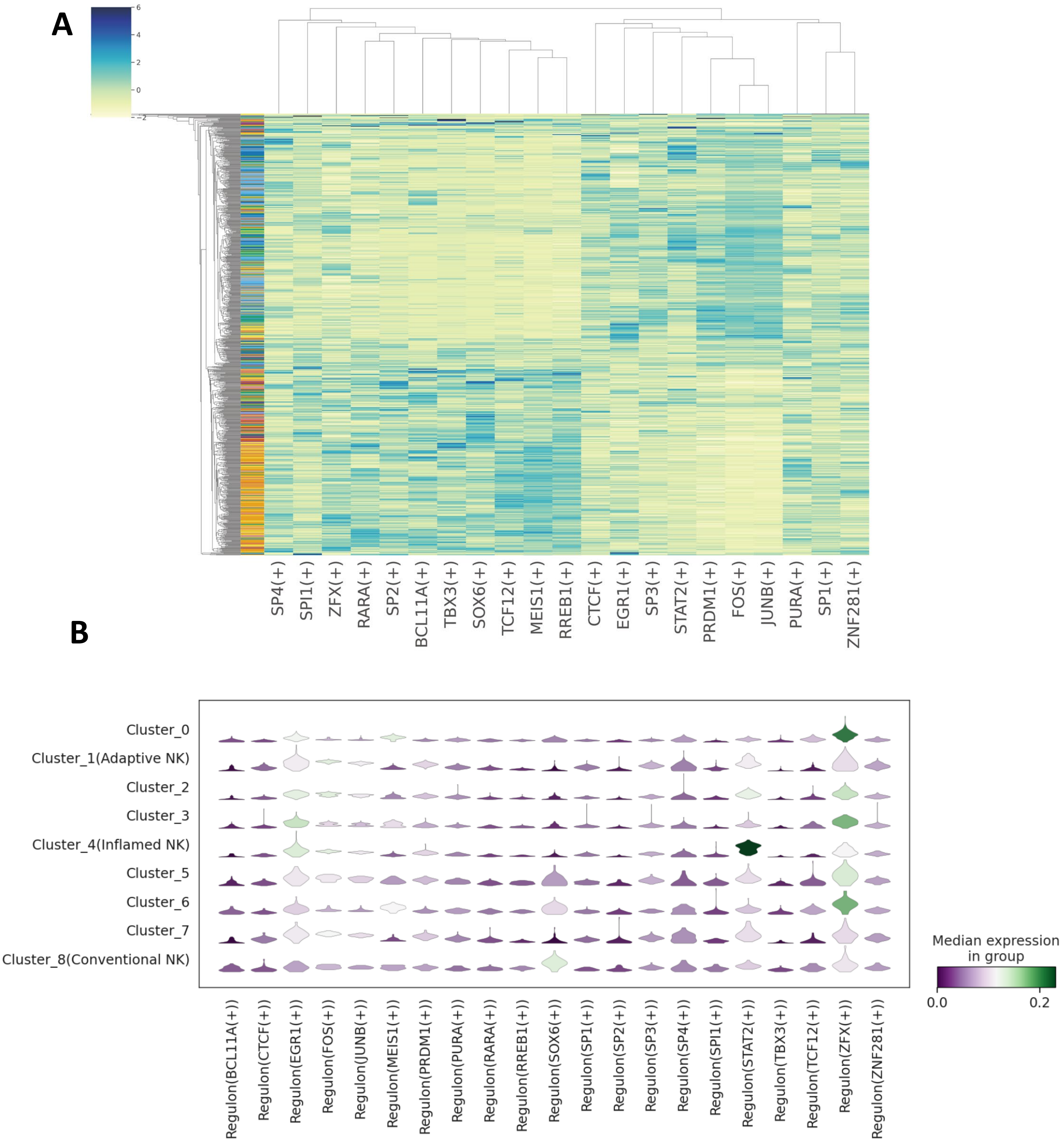


**Supplementary Figure 4. Heatmap and violin plots depicting the regulon specificity scores (RSS) of selected regulons across individual cells, alongside the median expression levels of these regulons across all identified clusters.** **A.** Heatmap displaying the RSS of selected transcription factor regulons across individual cells based on Smart-seq3 data. Each row represents a single cell, and each column corresponds to a specific regulon. **B.** Violin plots showing the distribution of RSS values for the same regulons across all identified clusters. Median RSS levels within each cluster are indicated by color intensity.

### Supplementary Figure 5

**A**

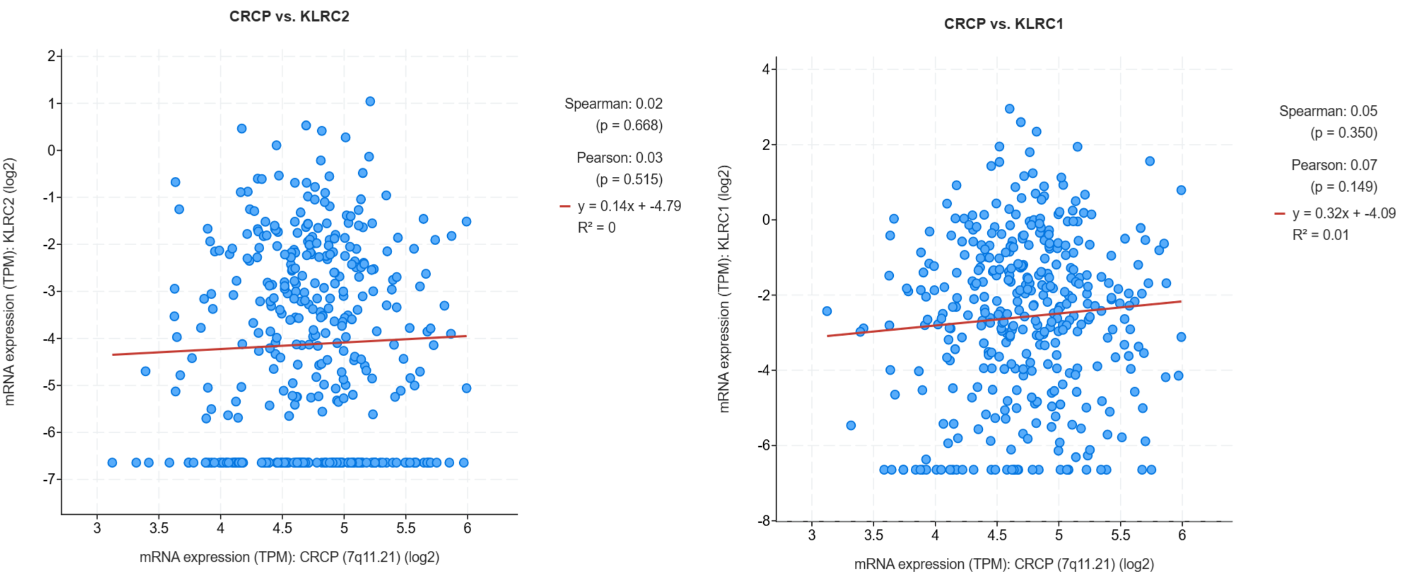


**Supplementary Figure 5. CRCP expression is not correlated with KLRC2 or KLRC1 in HGSOC.** Correlation analysis of CRCP expression with KLRC2 and KLRC1 in high-grade serous ovarian cancer (HGSOC) samples (n = 602, TCGA, accessed via cBio Cancer Genomics Portal) shows no significant association.

**Supplementary Table 1. Gene signatures used for NK subcluster annotation**

| CellType | tissueType | cellName | geneSymbolmore1(Positively expressed) | geneSymbolmore2 (Negatively expressed) |
|---|---|---|---|---|
| 0 | Immune system | Adaptive_NK_merged | FCGR3A | KLRC1,FCER1G,SYK |
| 0 | Immune system | Adaptive_NK_merged | FGFBP2 | KLRC1,FCER1G,SYK |
| 0 | Immune system | Adaptive_NK_merged | KLRF1 | KLRC1,FCER1G,SYK |
| 0 | Immune system | Adaptive_NK_merged | KLF2 | KLRC1,FCER1G,SYK |
| 0 | Immune system | Adaptive_NK_merged | PLAC8 | KLRC1,FCER1G,SYK |
| 0 | Immune system | Adaptive_NK_merged | GZMB | KLRC1,FCER1G,SYK |
| 0 | Immune system | Adaptive_NK_merged | GZMH | KLRC1,FCER1G,SYK |
| 0 | Immune system | Adaptive_NK_merged | CX3CR1 | KLRC1,FCER1G,SYK |
| 0 | Immune system | Adaptive_NK_merged | ARL4C | KLRC1,FCER1G,SYK |
| 0 | Immune system | Adaptive_NK_merged | PRF1 | KLRC1,FCER1G,SYK |
| 0 | Immune system | Adaptive_NK_merged | EFHD2 | KLRC1,FCER1G,SYK |
| 0 | Immune system | Adaptive_NK_merged | CYBA | KLRC1,FCER1G,SYK |
| 0 | Immune system | Adaptive_NK_merged | NKG7 | KLRC1,FCER1G,SYK |
| 0 | Immune system | Adaptive_NK_merged | PLEK | KLRC1,FCER1G,SYK |
| 0 | Immune system | Adaptive_NK_merged | S1PR5 | KLRC1,FCER1G,SYK |
| 0 | Immune system | Adaptive_NK_merged | CST7 | KLRC1,FCER1G,SYK |
| 0 | Immune system | Adaptive_NK_merged | ZEB2 | KLRC1,FCER1G,SYK |
| 0 | Immune system | Adaptive_NK_merged | DTHD1 | KLRC1,FCER1G,SYK |
| 0 | Immune system | Adaptive_NK_merged | SPON2 | KLRC1,FCER1G,SYK |
| 0 | Immune system | Adaptive_NK_merged | ADGRG1 | KLRC1,FCER1G,SYK |
| 0 | Immune system | Adaptive_NK_merged | RIPOR2 | KLRC1,FCER1G,SYK |
| 0 | Immune system | Adaptive_NK_merged | S100A4 | KLRC1,FCER1G,SYK |
| 0 | Immune system | Adaptive_NK_merged | TGFBR3 | KLRC1,FCER1G,SYK |
| 0 | Immune system | Adaptive_NK_merged | PRSS23 | KLRC1,FCER1G,SYK |
| 0 | Immune system | Adaptive_NK_merged | IGFBP7 | KLRC1,FCER1G,SYK |
| 0 | Immune system | Adaptive_NK_merged | GZMM | KLRC1,FCER1G,SYK |
| 0 | Immune system | Adaptive_NK_merged | TTC38 | KLRC1,FCER1G,SYK |
| 0 | Immune system | Adaptive_NK_merged | PFN1 | KLRC1,FCER1G,SYK |
| 0 | Immune system | Adaptive_NK_merged | CD247 | KLRC1,FCER1G,SYK |
| 0 | Immune system | Adaptive_NK_merged | CEP78 | KLRC1,FCER1G,SYK |
| 0 | Immune system | Adaptive_NK_merged | LITAF | KLRC1,FCER1G,SYK |
| 0 | Immune system | Adaptive_NK_merged | ACTB | KLRC1,FCER1G,SYK |
| 0 | Immune system | Adaptive_NK_merged | RAP1B | KLRC1,FCER1G,SYK |
| 0 | Immune system | Adaptive_NK_merged | FLNA | KLRC1,FCER1G,SYK |
| 0 | Immune system | Adaptive_NK_merged | EMP3 | KLRC1,FCER1G,SYK |
| 0 | Immune system | Adaptive_NK_merged | B2M | KLRC1,FCER1G,SYK |
| 0 | Immune system | Adaptive_NK_merged | C12orf75 | KLRC1,FCER1G,SYK |
| 0 | Immune system | Adaptive_NK_merged | CALM1 | KLRC1,FCER1G,SYK |
| 0 | Immune system | Adaptive_NK_merged | KLRD1 | KLRC1,FCER1G,SYK |
| 0 | Immune system | Adaptive_NK_merged | GNG2 | KLRC1,FCER1G,SYK |
| 0 | Immune system | Adaptive_NK_merged | TRBC1 | KLRC1,FCER1G,SYK |
| 0 | Immune system | Adaptive_NK_merged | CCL4L2 | KLRC1,FCER1G,SYK |
| 0 | Immune system | Adaptive_NK_merged | HLA-C | KLRC1,FCER1G,SYK |
| 0 | Immune system | Adaptive_NK_merged | ACTG1 | KLRC1,FCER1G,SYK |
| 0 | Immune system | Adaptive_NK_merged | ANXA1 | KLRC1,FCER1G,SYK |
| 0 | Immune system | Adaptive_NK_merged | C1orf21 | KLRC1,FCER1G,SYK |
| 0 | Immune system | Adaptive_NK_merged | SORL1 | KLRC1,FCER1G,SYK |
| 0 | Immune system | Adaptive_NK_merged | HLA-E | KLRC1,FCER1G,SYK |
| 0 | Immune system | Adaptive_NK_merged | AHNAK | KLRC1,FCER1G,SYK |
| 0 | Immune system | Adaptive_NK_merged | HLA-B | KLRC1,FCER1G,SYK |
| 0 | Immune system | Adaptive_NK_merged | ABHD17A | KLRC1,FCER1G,SYK |
| 0 | Immune system | Adaptive_NK_merged | FGL2 | KLRC1,FCER1G,SYK |

| | | | | |
|---|---|---|---|---|
| 0 | Immune system | Adaptive_NK_merged | PPP2R5C | KLRC1,FCER1G,SYK |
| 0 | Immune system | Adaptive_NK_merged | COL6A2 | KLRC1,FCER1G,SYK |
| 0 | Immune system | Adaptive_NK_merged | TSC22D3 | KLRC1,FCER1G,SYK |
| 0 | Immune system | Adaptive_NK_merged | S100A6 | KLRC1,FCER1G,SYK |
| 0 | Immune system | Adaptive_NK_merged | LYAR | KLRC1,FCER1G,SYK |
| 0 | Immune system | Adaptive_NK_merged | GK5 | KLRC1,FCER1G,SYK |
| 0 | Immune system | Adaptive_NK_merged | TPST2 | KLRC1,FCER1G,SYK |
| 0 | Immune system | Adaptive_NK_merged | LPCAT1 | KLRC1,FCER1G,SYK |
| 0 | Immune system | Adaptive_NK_merged | KIR2DL3 | KLRC1,FCER1G,SYK |
| 0 | Immune system | Adaptive_NK_merged | CFL1 | KLRC1,FCER1G,SYK |
| 0 | Immune system | Adaptive_NK_merged | SH3BP5 | KLRC1,FCER1G,SYK |
| 0 | Immune system | Adaptive_NK_merged | XBP1 | KLRC1,FCER1G,SYK |
| 0 | Immune system | Adaptive_NK_merged | ITGB2 | KLRC1,FCER1G,SYK |
| 0 | Immune system | Adaptive_NK_merged | FCRL6 | KLRC1,FCER1G,SYK |
| 0 | Immune system | Adaptive_NK_merged | PRKCB | KLRC1,FCER1G,SYK |
| 0 | Immune system | Adaptive_NK_merged | MYL12A | KLRC1,FCER1G,SYK |
| 0 | Immune system | Adaptive_NK_merged | HLA-DQA2 | KLRC1,FCER1G,SYK |
| 0 | Immune system | Adaptive_NK_merged | MYBL1 | KLRC1,FCER1G,SYK |
| 0 | Immune system | Adaptive_NK_merged | RAB29 | KLRC1,FCER1G,SYK |
| 0 | Immune system | Adaptive_NK_merged | DSTN | KLRC1,FCER1G,SYK |
| 0 | Immune system | Adaptive_NK_merged | PTGDR | KLRC1,FCER1G,SYK |
| 0 | Immune system | Adaptive_NK_merged | RASGRP2 | KLRC1,FCER1G,SYK |
| 0 | Immune system | Adaptive_NK_merged | MBP | KLRC1,FCER1G,SYK |
| 0 | Immune system | Adaptive_NK_merged | F2R | KLRC1,FCER1G,SYK |
| 0 | Immune system | Adaptive_NK_merged | ARPC2 | KLRC1,FCER1G,SYK |
| 0 | Immune system | Adaptive_NK_merged | SYNE2 | KLRC1,FCER1G,SYK |
| 0 | Immune system | Adaptive_NK_merged | LAIR1 | KLRC1,FCER1G,SYK |
| 0 | Immune system | Adaptive_NK_merged | CCL4 | KLRC1,FCER1G,SYK |
| 0 | Immune system | Adaptive_NK_merged | CD47 | KLRC1,FCER1G,SYK |
| 0 | Immune system | Adaptive_NK_merged | TBX21 | KLRC1,FCER1G,SYK |
| 0 | Immune system | Adaptive_NK_merged | CMC1 | KLRC1,FCER1G,SYK |
| 0 | Immune system | Adaptive_NK_merged | ANXA4 | KLRC1,FCER1G,SYK |
| 0 | Immune system | Adaptive_NK_merged | RGS9 | KLRC1,FCER1G,SYK |
| 0 | Immune system | Adaptive_NK_merged | TMSB10 | KLRC1,FCER1G,SYK |
| 0 | Immune system | Adaptive_NK_merged | PRDM1 | KLRC1,FCER1G,SYK |
| 0 | Immune system | Adaptive_NK_merged | RASA3 | KLRC1,FCER1G,SYK |
| 0 | Immune system | Adaptive_NK_merged | CD53 | KLRC1,FCER1G,SYK |
| 0 | Immune system | Adaptive_NK_merged | SPN | KLRC1,FCER1G,SYK |
| 0 | Immune system | Adaptive_NK_merged | ADRB2 | KLRC1,FCER1G,SYK |
| 0 | Immune system | Adaptive_NK_merged | KLF3 | KLRC1,FCER1G,SYK |
| 0 | Immune system | Adaptive_NK_merged | ADD3 | KLRC1,FCER1G,SYK |
| 0 | Immune system | Adaptive_NK_merged | HLA-F | KLRC1,FCER1G,SYK |
| 0 | Immune system | Adaptive_NK_merged | TFDP2 | KLRC1,FCER1G,SYK |
| 0 | Immune system | Adaptive_NK_merged | SSBP3 | KLRC1,FCER1G,SYK |
| 0 | Immune system | Adaptive_NK_merged | SYNE1 | KLRC1,FCER1G,SYK |
| 0 | Immune system | Adaptive_NK_merged | SYTL3 | KLRC1,FCER1G,SYK |
| 0 | Immune system | Adaptive_NK_merged | GLRX | KLRC1,FCER1G,SYK |
| 0 | Immune system | Adaptive_NK_merged | HLA-DPB1 | KLRC1,FCER1G,SYK |
| 0 | Immune system | Adaptive_NK_merged | RAP2B | KLRC1,FCER1G,SYK |
| 0 | Immune system | Adaptive_NK_merged | UCP2 | KLRC1,FCER1G,SYK |
| 0 | Immune system | Adaptive_NK_merged | TXNIP | KLRC1,FCER1G,SYK |
| 0 | Immune system | Adaptive_NK_merged | VCL | KLRC1,FCER1G,SYK |
| 0 | Immune system | Adaptive_NK_merged | MGAT4A | KLRC1,FCER1G,SYK |
| 0 | Immune system | Adaptive_NK_merged | GZMA | KLRC1,FCER1G,SYK |
| 0 | Immune system | Adaptive_NK_merged | GTF3C1 | KLRC1,FCER1G,SYK |
| 0 | Immune system | Adaptive_NK_merged | PTGER2 | KLRC1,FCER1G,SYK |
| 0 | Immune system | Adaptive_NK_merged | PTPRC | KLRC1,FCER1G,SYK |
| 0 | Immune system | Adaptive_NK_merged | BIN2 | KLRC1,FCER1G,SYK |

| | | | | |
|---|---|---|---|---|
| 0 | Immune system | Adaptive_NK_merged | GIMAP4 | KLRC1,FCER1G,SYK |
| 0 | Immune system | Adaptive_NK_merged | PXN | KLRC1,FCER1G,SYK |
| 0 | Immune system | Adaptive_NK_merged | ABI3 | KLRC1,FCER1G,SYK |
| 0 | Immune system | Adaptive_NK_merged | ADGRE5 | KLRC1,FCER1G,SYK |
| 0 | Immune system | Adaptive_NK_merged | SRGN | KLRC1,FCER1G,SYK |
| 0 | Immune system | Adaptive_NK_merged | HDDC2 | KLRC1,FCER1G,SYK |
| 0 | Immune system | Adaptive_NK_merged | ICAM2 | KLRC1,FCER1G,SYK |
| 0 | Immune system | Adaptive_NK_merged | TPM4 | KLRC1,FCER1G,SYK |
| 0 | Immune system | Adaptive_NK_merged | KLRC2 | KLRC1,FCER1G,SYK |
| 0 | Immune system | Adaptive_NK_merged | CAP1 | KLRC1,FCER1G,SYK |
| 0 | Immune system | Adaptive_NK_merged | LINC00861 | KLRC1,FCER1G,SYK |
| 0 | Immune system | Adaptive_NK_merged | GRAP2 | KLRC1,FCER1G,SYK |
| 0 | Immune system | Adaptive_NK_merged | EIF1 | KLRC1,FCER1G,SYK |
| 0 | Immune system | Adaptive_NK_merged | IFNG | KLRC1,FCER1G,SYK |
| 0 | Immune system | Adaptive_NK_merged | ITGAL | KLRC1,FCER1G,SYK |
| 0 | Immune system | Adaptive_NK_merged | PTPRE | KLRC1,FCER1G,SYK |
| 0 | Immune system | Adaptive_NK_merged | PTGER4 | KLRC1,FCER1G,SYK |
| 0 | Immune system | Adaptive_NK_merged | HLA-DPA1 | KLRC1,FCER1G,SYK |
| 0 | Immune system | Adaptive_NK_merged | CDC42SE1 | KLRC1,FCER1G,SYK |
| 0 | Immune system | Adaptive_NK_merged | MAF | KLRC1,FCER1G,SYK |
| 0 | Immune system | Adaptive_NK_merged | CTBP2 | KLRC1,FCER1G,SYK |
| 0 | Immune system | Adaptive_NK_merged | CDKN2D | KLRC1,FCER1G,SYK |
| 0 | Immune system | Adaptive_NK_merged | CYTH1 | KLRC1,FCER1G,SYK |
| 0 | Immune system | Adaptive_NK_merged | MYO1G | KLRC1,FCER1G,SYK |
| 0 | Immune system | Adaptive_NK_merged | SH3KBP1 | KLRC1,FCER1G,SYK |
| 0 | Immune system | Adaptive_NK_merged | YWHAB | KLRC1,FCER1G,SYK |
| 0 | Immune system | Adaptive_NK_merged | CDC25B | KLRC1,FCER1G,SYK |
| 0 | Immune system | Adaptive_NK_merged | ATM | KLRC1,FCER1G,SYK |
| 0 | Immune system | Adaptive_NK_merged | ACTR3 | KLRC1,FCER1G,SYK |
| 0 | Immune system | Adaptive_NK_merged | RAB9A | KLRC1,FCER1G,SYK |
| 0 | Immune system | Adaptive_NK_merged | RORA | KLRC1,FCER1G,SYK |
| 0 | Immune system | Adaptive_NK_merged | GAB3 | KLRC1,FCER1G,SYK |
| 0 | Immune system | Adaptive_NK_merged | KLF13 | KLRC1,FCER1G,SYK |
| 0 | Immune system | Adaptive_NK_merged | KLF6 | KLRC1,FCER1G,SYK |
| 0 | Immune system | Adaptive_NK_merged | CAST | KLRC1,FCER1G,SYK |
| 0 | Immune system | Adaptive_NK_merged | IQGAP2 | KLRC1,FCER1G,SYK |
| 0 | Immune system | Adaptive_NK_merged | RPL3 | KLRC1,FCER1G,SYK |
| 0 | Immune system | Adaptive_NK_merged | CD320 | KLRC1,FCER1G,SYK |
| 0 | Immune system | Adaptive_NK_merged | IGF2R | KLRC1,FCER1G,SYK |
| 0 | Immune system | Adaptive_NK_merged | IFITM1 | KLRC1,FCER1G,SYK |
| 0 | Immune system | Adaptive_NK_merged | GPSM3 | KLRC1,FCER1G,SYK |
| 0 | Immune system | Adaptive_NK_merged | FTL | KLRC1,FCER1G,SYK |
| 0 | Immune system | Adaptive_NK_merged | RHOG | KLRC1,FCER1G,SYK |
| 0 | Immune system | Adaptive_NK_merged | CD3E | KLRC1,FCER1G,SYK |
| 0 | Immune system | Adaptive_NK_merged | EBP | KLRC1,FCER1G,SYK |
| 0 | Immune system | Adaptive_NK_merged | VASP | KLRC1,FCER1G,SYK |
| 0 | Immune system | Adaptive_NK_merged | ARID5B | KLRC1,FCER1G,SYK |
| 0 | Immune system | Adaptive_NK_merged | SH2D2A | KLRC1,FCER1G,SYK |
| 0 | Immune system | Adaptive_NK_merged | MIAT | KLRC1,FCER1G,SYK |
| 0 | Immune system | Adaptive_NK_merged | PTP4A2 | KLRC1,FCER1G,SYK |
| 0 | Immune system | Adaptive_NK_merged | HSPA8 | KLRC1,FCER1G,SYK |
| 0 | Immune system | Adaptive_NK_merged | IFITM2 | KLRC1,FCER1G,SYK |
| 0 | Immune system | Adaptive_NK_merged | PYHIN1 | KLRC1,FCER1G,SYK |
| 0 | Immune system | Adaptive_NK_merged | NDUFB2 | KLRC1,FCER1G,SYK |
| 0 | Immune system | Adaptive_NK_merged | RPA2 | KLRC1,FCER1G,SYK |
| 0 | Immune system | Adaptive_NK_merged | HIPK2 | KLRC1,FCER1G,SYK |
| 0 | Immune system | Adaptive_NK_merged | SIGIRR | KLRC1,FCER1G,SYK |
| 0 | Immune system | Adaptive_NK_merged | EIF4G3 | KLRC1,FCER1G,SYK |

| | | | | |
|---|---|---|---|---|
| 0 | Immune system | Adaptive_NK_merged | NDUFB7 | KLRC1,FCER1G,SYK |
| 0 | Immune system | Adaptive_NK_merged | APMAP | KLRC1,FCER1G,SYK |
| 0 | Immune system | Adaptive_NK_merged | CD300A | KLRC1,FCER1G,SYK |
| 0 | Immune system | Adaptive_NK_merged | CTSC | KLRC1,FCER1G,SYK |
| 0 | Immune system | Adaptive_NK_merged | CCDC88C | KLRC1,FCER1G,SYK |
| 0 | Immune system | Adaptive_NK_merged | PDIA3 | KLRC1,FCER1G,SYK |
| 0 | Immune system | Adaptive_NK_merged | STK10 | KLRC1,FCER1G,SYK |
| 0 | Immune system | Adaptive_NK_merged | TOB1 | KLRC1,FCER1G,SYK |
| 0 | Immune system | Adaptive_NK_merged | YWHAZ | KLRC1,FCER1G,SYK |
| 0 | Immune system | Adaptive_NK_merged | PPIB | KLRC1,FCER1G,SYK |
| 0 | Immune system | Adaptive_NK_merged | METRNL | KLRC1,FCER1G,SYK |
| 0 | Immune system | Adaptive_NK_merged | HLA-DRB1 | KLRC1,FCER1G,SYK |
| 0 | Immune system | Adaptive_NK_merged | FAM49B | KLRC1,FCER1G,SYK |
| 0 | Immune system | Adaptive_NK_merged | ARPC5 | KLRC1,FCER1G,SYK |
| 0 | Immune system | Adaptive_NK_merged | MYADM | KLRC1,FCER1G,SYK |
| 0 | Immune system | Adaptive_NK_merged | TERF1 | KLRC1,FCER1G,SYK |
| 0 | Immune system | Adaptive_NK_merged | ARHGDIB | KLRC1,FCER1G,SYK |
| 0 | Immune system | Adaptive_NK_merged | PPP1CA | KLRC1,FCER1G,SYK |
| 0 | Immune system | Adaptive_NK_merged | PTPN12 | KLRC1,FCER1G,SYK |
| 0 | Immune system | Adaptive_NK_merged | VAV3 | KLRC1,FCER1G,SYK |
| 0 | Immune system | Adaptive_NK_merged | RTN4 | KLRC1,FCER1G,SYK |
| 0 | Immune system | Adaptive_NK_merged | WDR1 | KLRC1,FCER1G,SYK |
| 0 | Immune system | Adaptive_NK_merged | TLE4 | KLRC1,FCER1G,SYK |
| 0 | Immune system | Adaptive_NK_merged | P4HB | KLRC1,FCER1G,SYK |
| 0 | Immune system | Adaptive_NK_merged | ZBTB38 | KLRC1,FCER1G,SYK |
| 0 | Immune system | Adaptive_NK_merged | CLIC3 | KLRC1,FCER1G,SYK |
| 0 | Immune system | Adaptive_NK_merged | RGS19 | KLRC1,FCER1G,SYK |
| 0 | Immune system | Adaptive_NK_merged | USP28 | KLRC1,FCER1G,SYK |
| 0 | Immune system | Adaptive_NK_merged | FGR | KLRC1,FCER1G,SYK |
| 0 | Immune system | Adaptive_NK_merged | TNFRSF1B | KLRC1,FCER1G,SYK |
| 0 | Immune system | Adaptive_NK_merged | RPL21 | KLRC1,FCER1G,SYK |
| 0 | Immune system | Adaptive_NK_merged | OSTF1 | KLRC1,FCER1G,SYK |
| 0 | Immune system | Adaptive_NK_merged | PRR5L | KLRC1,FCER1G,SYK |
| 0 | Immune system | Adaptive_NK_merged | HSPA5 | KLRC1,FCER1G,SYK |
| 0 | Immune system | Adaptive_NK_merged | TMBIM6 | KLRC1,FCER1G,SYK |
| 0 | Immune system | Adaptive_NK_merged | DBI | KLRC1,FCER1G,SYK |
| 0 | Immune system | Adaptive_NK_merged | TMEM181 | KLRC1,FCER1G,SYK |
| 0 | Immune system | Adaptive_NK_merged | ACTR2 | KLRC1,FCER1G,SYK |
| 0 | Immune system | Adaptive_NK_merged | CORO1A | KLRC1,FCER1G,SYK |
| 0 | Immune system | Adaptive_NK_merged | RAC2 | KLRC1,FCER1G,SYK |
| 0 | Immune system | Adaptive_NK_merged | ARHGAP25 | KLRC1,FCER1G,SYK |
| 0 | Immune system | Adaptive_NK_merged | UPP1 | KLRC1,FCER1G,SYK |
| 0 | Immune system | Adaptive_NK_merged | LYN | KLRC1,FCER1G,SYK |
| 0 | Immune system | Adaptive_NK_merged | TMEM173 | KLRC1,FCER1G,SYK |
| 0 | Immune system | Adaptive_NK_merged | AES | KLRC1,FCER1G,SYK |
| 0 | Immune system | Adaptive_NK_merged | MYO1F | KLRC1,FCER1G,SYK |
| 0 | Immune system | Adaptive_NK_merged | SASH3 | KLRC1,FCER1G,SYK |
| 0 | Immune system | Adaptive_NK_merged | BATF | KLRC1,FCER1G,SYK |
| 0 | Immune system | Adaptive_NK_merged | GIMAP1 | KLRC1,FCER1G,SYK |
| 0 | Immune system | Adaptive_NK_merged | UBE2F | KLRC1,FCER1G,SYK |
| 0 | Immune system | Adaptive_NK_merged | CAPN2 | KLRC1,FCER1G,SYK |
| 0 | Immune system | Adaptive_NK_merged | SLC9A3R1 | KLRC1,FCER1G,SYK |
| 0 | Immune system | Adaptive_NK_merged | SPCS3 | KLRC1,FCER1G,SYK |
| 0 | Immune system | Adaptive_NK_merged | GMFG | KLRC1,FCER1G,SYK |
| 0 | Immune system | Adaptive_NK_merged | SMAP2 | KLRC1,FCER1G,SYK |
| 0 | Immune system | Adaptive_NK_merged | HNRNPF | KLRC1,FCER1G,SYK |
| 0 | Immune system | Adaptive_NK_merged | CD99 | KLRC1,FCER1G,SYK |
| 0 | Immune system | Adaptive_NK_merged | CDC42 | KLRC1,FCER1G,SYK |

| | | | | |
|---|---|---|---|---|
| 0 | Immune system | Adaptive_NK_merged | RHOA | KLRC1,FCER1G,SYK |
| 0 | Immune system | Adaptive_NK_merged | ARPC4 | KLRC1,FCER1G,SYK |
| 0 | Immune system | Adaptive_NK_merged | BIN1 | KLRC1,FCER1G,SYK |
| 0 | Immune system | Adaptive_NK_merged | ORAI1 | KLRC1,FCER1G,SYK |
| 0 | Immune system | Adaptive_NK_merged | DHRS7 | KLRC1,FCER1G,SYK |
| 0 | Immune system | Adaptive_NK_merged | ANXA2 | KLRC1,FCER1G,SYK |
| 0 | Immune system | Adaptive_NK_merged | TPP1 | KLRC1,FCER1G,SYK |
| 0 | Immune system | Adaptive_NK_merged | ATP2B4 | KLRC1,FCER1G,SYK |
| 0 | Immune system | Adaptive_NK_merged | CDC42EP3 | KLRC1,FCER1G,SYK |
| 0 | Immune system | Adaptive_NK_merged | UTRN | KLRC1,FCER1G,SYK |
| 0 | Immune system | Adaptive_NK_merged | GNAI2 | KLRC1,FCER1G,SYK |
| 0 | Immune system | Adaptive_NK_merged | FYN | KLRC1,FCER1G,SYK |
| 0 | Immune system | Adaptive_NK_merged | TES | KLRC1,FCER1G,SYK |
| 0 | Immune system | Adaptive_NK_merged | SDF2L1 | KLRC1,FCER1G,SYK |
| 0 | Immune system | Adaptive_NK_merged | TGFBR1 | KLRC1,FCER1G,SYK |
| 0 | Immune system | Adaptive_NK_merged | RNF19A | KLRC1,FCER1G,SYK |
| 0 | Immune system | Adaptive_NK_merged | KLRG1 | KLRC1,FCER1G,SYK |
| 0 | Immune system | Adaptive_NK_merged | PARP15 | KLRC1,FCER1G,SYK |
| 0 | Immune system | Adaptive_NK_merged | MED15 | KLRC1,FCER1G,SYK |
| 0 | Immune system | Adaptive_NK_merged | SLC15A4 | KLRC1,FCER1G,SYK |
| 0 | Immune system | Adaptive_NK_merged | S1PR4 | KLRC1,FCER1G,SYK |
| 0 | Immune system | Adaptive_NK_merged | LLGL2 | KLRC1,FCER1G,SYK |
| 0 | Immune system | Adaptive_NK_merged | CALR | KLRC1,FCER1G,SYK |
| 0 | Immune system | Adaptive_NK_merged | SFT2D1 | KLRC1,FCER1G,SYK |
| 0 | Immune system | Adaptive_NK_merged | MIEN1 | KLRC1,FCER1G,SYK |
| 0 | Immune system | Adaptive_NK_merged | TUBA4A | KLRC1,FCER1G,SYK |
| 0 | Immune system | Adaptive_NK_merged | NCOA1 | KLRC1,FCER1G,SYK |
| 0 | Immune system | Adaptive_NK_merged | SUN2 | KLRC1,FCER1G,SYK |
| 0 | Immune system | Adaptive_NK_merged | UHMK1 | KLRC1,FCER1G,SYK |
| 0 | Immune system | Adaptive_NK_merged | ANXA6 | KLRC1,FCER1G,SYK |
| 0 | Immune system | Adaptive_NK_merged | MIDN | KLRC1,FCER1G,SYK |
| 0 | Immune system | Adaptive_NK_merged | TNFRSF14 | KLRC1,FCER1G,SYK |
| 0 | Immune system | Adaptive_NK_merged | PTPN18 | KLRC1,FCER1G,SYK |
| 0 | Immune system | Adaptive_NK_merged | GPR65 | KLRC1,FCER1G,SYK |
| 0 | Immune system | Adaptive_NK_merged | AKNA | KLRC1,FCER1G,SYK |
| 0 | Immune system | Adaptive_NK_merged | MANF | KLRC1,FCER1G,SYK |
| 0 | Immune system | Adaptive_NK_merged | LCK | KLRC1,FCER1G,SYK |
| 0 | Immune system | Adaptive_NK_merged | LMAN2 | KLRC1,FCER1G,SYK |
| 0 | Immune system | Adaptive_NK_merged | MOB3A | KLRC1,FCER1G,SYK |
| 0 | Immune system | Adaptive_NK_merged | PRELID1 | KLRC1,FCER1G,SYK |
| 0 | Immune system | Adaptive_NK_merged | CAPZB | KLRC1,FCER1G,SYK |
| 0 | Immune system | Adaptive_NK_merged | CREM | KLRC1,FCER1G,SYK |
| 0 | Immune system | Adaptive_NK_merged | FBXW5 | KLRC1,FCER1G,SYK |
| 0 | Immune system | Adaptive_NK_merged | CDK2AP2 | KLRC1,FCER1G,SYK |
| 0 | Immune system | Adaptive_NK_merged | SH3BP2 | KLRC1,FCER1G,SYK |
| 0 | Immune system | Adaptive_NK_merged | MSN | KLRC1,FCER1G,SYK |
| 0 | Immune system | Adaptive_NK_merged | PPIA | KLRC1,FCER1G,SYK |
| 0 | Immune system | Adaptive_NK_merged | YWHAQ | KLRC1,FCER1G,SYK |
| 0 | Immune system | Adaptive_NK_merged | PRMT2 | KLRC1,FCER1G,SYK |
| 0 | Immune system | Adaptive_NK_merged | ATP1A1 | KLRC1,FCER1G,SYK |
| 0 | Immune system | Adaptive_NK_merged | CLIC1 | KLRC1,FCER1G,SYK |
| 0 | Immune system | Adaptive_NK_merged | STK38 | KLRC1,FCER1G,SYK |
| 0 | Immune system | Adaptive_NK_merged | DIAPH1 | KLRC1,FCER1G,SYK |
| 0 | Immune system | Adaptive_NK_merged | DENND2D | KLRC1,FCER1G,SYK |
| 0 | Immune system | Adaptive_NK_merged | S100A10 | KLRC1,FCER1G,SYK |
| 0 | Immune system | Adaptive_NK_merged | SUPT4H1 | KLRC1,FCER1G,SYK |
| 0 | Immune system | Adaptive_NK_merged | ATP2B1-AS1 | KLRC1,FCER1G,SYK |
| 0 | Immune system | Adaptive_NK_merged | SYNGR1 | KLRC1,FCER1G,SYK |

| | | | | |
|---|---|---|---|---|
| 0 | Immune system | Adaptive_NK_merged | MAPRE2 | KLRC1,FCER1G,SYK |
| 0 | Immune system | Adaptive_NK_merged | HSP90B1 | KLRC1,FCER1G,SYK |
| 0 | Immune system | Adaptive_NK_merged | G6PD | KLRC1,FCER1G,SYK |
| 0 | Immune system | Adaptive_NK_merged | TBCB | KLRC1,FCER1G,SYK |
| 0 | Immune system | Adaptive_NK_merged | TRPV2 | KLRC1,FCER1G,SYK |
| 0 | Immune system | Adaptive_NK_merged | GRK6 | KLRC1,FCER1G,SYK |
| 0 | Immune system | Adaptive_NK_merged | TIMP1 | KLRC1,FCER1G,SYK |
| 0 | Immune system | Adaptive_NK_merged | MTPN | KLRC1,FCER1G,SYK |
| 0 | Immune system | Adaptive_NK_merged | LDLR | KLRC1,FCER1G,SYK |
| 0 | Immune system | Adaptive_NK_merged | ZAP70 | KLRC1,FCER1G,SYK |
| 0 | Immune system | Adaptive_NK_merged | C12orf57 | KLRC1,FCER1G,SYK |
| 0 | Immune system | Adaptive_NK_merged | CISD3 | KLRC1,FCER1G,SYK |
| 0 | Immune system | Adaptive_NK_merged | RNF167 | KLRC1,FCER1G,SYK |
| 0 | Immune system | Adaptive_NK_merged | IVNS1ABP | KLRC1,FCER1G,SYK |
| 0 | Immune system | Adaptive_NK_merged | ARL6IP1 | KLRC1,FCER1G,SYK |
| 0 | Immune system | Adaptive_NK_merged | PPP1R18 | KLRC1,FCER1G,SYK |
| 0 | Immune system | Adaptive_NK_merged | TPM3 | KLRC1,FCER1G,SYK |
| 0 | Immune system | Adaptive_NK_merged | GIMAP7 | KLRC1,FCER1G,SYK |
| 0 | Immune system | Adaptive_NK_merged | SNF8 | KLRC1,FCER1G,SYK |
| 0 | Immune system | Adaptive_NK_merged | SEC11C | KLRC1,FCER1G,SYK |
| 0 | Immune system | Adaptive_NK_merged | LBR | KLRC1,FCER1G,SYK |
| 0 | Immune system | Adaptive_NK_merged | RAB8A | KLRC1,FCER1G,SYK |
| 0 | Immune system | Adaptive_NK_merged | ZNF276 | KLRC1,FCER1G,SYK |
| 0 | Immune system | Adaptive_NK_merged | PIP4K2A | KLRC1,FCER1G,SYK |
| 0 | Immune system | Adaptive_NK_merged | RPS26 | KLRC1,FCER1G,SYK |
| 0 | Immune system | Adaptive_NK_merged | RASSF1 | KLRC1,FCER1G,SYK |
| 0 | Immune system | Adaptive_NK_merged | BCL11B | KLRC1,FCER1G,SYK |
| 0 | Immune system | Adaptive_NK_merged | PCBP1 | KLRC1,FCER1G,SYK |
| 0 | Immune system | Adaptive_NK_merged | FKBP11 | KLRC1,FCER1G,SYK |
| 0 | Immune system | Adaptive_NK_merged | TMEM50A | KLRC1,FCER1G,SYK |
| 0 | Immune system | Adaptive_NK_merged | MYDGF | KLRC1,FCER1G,SYK |
| 0 | Immune system | Adaptive_NK_merged | RNF126 | KLRC1,FCER1G,SYK |
| 0 | Immune system | Adaptive_NK_merged | RRBP1 | KLRC1,FCER1G,SYK |
| 0 | Immune system | Adaptive_NK_merged | PLEKHA1 | KLRC1,FCER1G,SYK |
| 0 | Immune system | Adaptive_NK_merged | SELPLG | KLRC1,FCER1G,SYK |
| 0 | Immune system | Adaptive_NK_merged | SH2D1B | KLRC1,FCER1G,SYK |
| 0 | Immune system | Adaptive_NK_merged | RAB5C | KLRC1,FCER1G,SYK |
| 0 | Immune system | Adaptive_NK_merged | ARRDC3 | KLRC1,FCER1G,SYK |
| 0 | Immune system | Adaptive_NK_merged | ARL6IP5 | KLRC1,FCER1G,SYK |
| 0 | Immune system | Adaptive_NK_merged | TMEM2 | KLRC1,FCER1G,SYK |
| 0 | Immune system | Adaptive_NK_merged | CCDC85B | KLRC1,FCER1G,SYK |
| 0 | Immune system | Adaptive_NK_merged | ARPC1B | KLRC1,FCER1G,SYK |
| 0 | Immune system | Adaptive_NK_merged | SIRT2 | KLRC1,FCER1G,SYK |
| 0 | Immune system | Adaptive_NK_merged | CD226 | KLRC1,FCER1G,SYK |
| 0 | Immune system | Adaptive_NK_merged | LRRFIP1 | KLRC1,FCER1G,SYK |
| 0 | Immune system | Adaptive_NK_merged | RAB10 | KLRC1,FCER1G,SYK |
| 0 | Immune system | Adaptive_NK_merged | ATP1B3 | KLRC1,FCER1G,SYK |
| 0 | Immune system | Adaptive_NK_merged | PDIA6 | KLRC1,FCER1G,SYK |
| 0 | Immune system | Adaptive_NK_merged | SRPK2 | KLRC1,FCER1G,SYK |
| 0 | Immune system | Adaptive_NK_merged | IQGAP1 | KLRC1,FCER1G,SYK |
| 0 | Immune system | Adaptive_NK_merged | PGAM1 | KLRC1,FCER1G,SYK |
| 0 | Immune system | Adaptive_NK_merged | CAPNS1 | KLRC1,FCER1G,SYK |
| 0 | Immune system | Adaptive_NK_merged | TMED9 | KLRC1,FCER1G,SYK |
| 0 | Immune system | Adaptive_NK_merged | MFSD10 | KLRC1,FCER1G,SYK |
| 0 | Immune system | Adaptive_NK_merged | ATP5F1 | KLRC1,FCER1G,SYK |
| 0 | Immune system | Adaptive_NK_merged | DECR1 | KLRC1,FCER1G,SYK |
| 0 | Immune system | Adaptive_NK_merged | PDLIM2 | KLRC1,FCER1G,SYK |
| 0 | Immune system | Adaptive_NK_merged | SERPINB1 | KLRC1,FCER1G,SYK |

| | | | | |
|---|---|---|---|---|
| 0 | Immune system | Adaptive_NK_merged | CCDC82 | KLRC1,FCER1G,SYK |
| 0 | Immune system | Adaptive_NK_merged | CEBPB | KLRC1,FCER1G,SYK |
| 0 | Immune system | Adaptive_NK_merged | ITGB1BP1 | KLRC1,FCER1G,SYK |
| 0 | Immune system | Adaptive_NK_merged | M6PR | KLRC1,FCER1G,SYK |
| 0 | Immune system | Adaptive_NK_merged | TMEM59 | KLRC1,FCER1G,SYK |
| 0 | Immune system | Adaptive_NK_merged | RECQL | KLRC1,FCER1G,SYK |
| 0 | Immune system | Adaptive_NK_merged | RGS14 | KLRC1,FCER1G,SYK |
| 0 | Immune system | Adaptive_NK_merged | PLEKHF1 | KLRC1,FCER1G,SYK |
| 0 | Immune system | Adaptive_NK_merged | NFATC2 | KLRC1,FCER1G,SYK |
| 0 | Immune system | Adaptive_NK_merged | PPP2R5A | KLRC1,FCER1G,SYK |
| 0 | Immune system | Adaptive_NK_merged | SEC61B | KLRC1,FCER1G,SYK |
| 0 | Immune system | Adaptive_NK_merged | SYTL1 | KLRC1,FCER1G,SYK |
| 0 | Immune system | Adaptive_NK_merged | SUB1 | KLRC1,FCER1G,SYK |
| 0 | Immune system | Adaptive_NK_merged | TLN1 | KLRC1,FCER1G,SYK |
| 0 | Immune system | Adaptive_NK_merged | SPCS2 | KLRC1,FCER1G,SYK |
| 0 | Immune system | Adaptive_NK_merged | PDAP1 | KLRC1,FCER1G,SYK |
| 0 | Immune system | Adaptive_NK_merged | VAMP2 | KLRC1,FCER1G,SYK |
| 0 | Immune system | Adaptive_NK_merged | SLAMF7 | KLRC1,FCER1G,SYK |
| 0 | Immune system | Adaptive_NK_merged | CRBN | KLRC1,FCER1G,SYK |
| 0 | Immune system | Adaptive_NK_merged | SELENOT | KLRC1,FCER1G,SYK |
| 0 | Immune system | Adaptive_NK_merged | TECR | KLRC1,FCER1G,SYK |
| 0 | Immune system | Adaptive_NK_merged | RNF149 | KLRC1,FCER1G,SYK |
| 0 | Immune system | Adaptive_NK_merged | SIPA1 | KLRC1,FCER1G,SYK |
| 0 | Immune system | Adaptive_NK_merged | EIF4EBP2 | KLRC1,FCER1G,SYK |
| 0 | Immune system | Adaptive_NK_merged | HSH2D | KLRC1,FCER1G,SYK |
| 0 | Immune system | Adaptive_NK_merged | IMP3 | KLRC1,FCER1G,SYK |
| 0 | Immune system | Adaptive_NK_merged | CIB1 | KLRC1,FCER1G,SYK |
| 0 | Immune system | Adaptive_NK_merged | LRP10 | KLRC1,FCER1G,SYK |
| 0 | Immune system | Adaptive_NK_merged | ARPC5L | KLRC1,FCER1G,SYK |
| 0 | Immune system | Adaptive_NK_merged | RAPGEF1 | KLRC1,FCER1G,SYK |
| 0 | Immune system | Adaptive_NK_merged | POLR3GL | KLRC1,FCER1G,SYK |
| 0 | Immune system | Adaptive_NK_merged | RNF169 | KLRC1,FCER1G,SYK |
| 0 | Immune system | Adaptive_NK_merged | ZFAND6 | KLRC1,FCER1G,SYK |
| 0 | Immune system | Adaptive_NK_merged | ARHGAP30 | KLRC1,FCER1G,SYK |
| 0 | Immune system | Adaptive_NK_merged | STOM | KLRC1,FCER1G,SYK |
| 0 | Immune system | Adaptive_NK_merged | TCF25 | KLRC1,FCER1G,SYK |
| 0 | Immune system | Adaptive_NK_merged | CTDSP1 | KLRC1,FCER1G,SYK |
| 0 | Immune system | Adaptive_NK_merged | PSMB10 | KLRC1,FCER1G,SYK |
| 0 | Immune system | Adaptive_NK_merged | ATOX1 | KLRC1,FCER1G,SYK |
| 0 | Immune system | Adaptive_NK_merged | SYAP1 | KLRC1,FCER1G,SYK |
| 0 | Immune system | Adaptive_NK_merged | CYB561D2 | KLRC1,FCER1G,SYK |
| 0 | Immune system | Adaptive_NK_merged | EIF4B | KLRC1,FCER1G,SYK |
| 0 | Immune system | Adaptive_NK_merged | TAF7 | KLRC1,FCER1G,SYK |
| 0 | Immune system | Adaptive_NK_merged | SRPRA | KLRC1,FCER1G,SYK |
| 0 | Immune system | Adaptive_NK_merged | CANX | KLRC1,FCER1G,SYK |
| 0 | Immune system | Adaptive_NK_merged | RNASEH2C | KLRC1,FCER1G,SYK |
| 0 | Immune system | Adaptive_NK_merged | IDI1 | KLRC1,FCER1G,SYK |
| 0 | Immune system | Adaptive_NK_merged | SSBP4 | KLRC1,FCER1G,SYK |
| 0 | Immune system | Adaptive_NK_merged | C16orf54 | KLRC1,FCER1G,SYK |
| 0 | Immune system | Adaptive_NK_merged | CYFIP2 | KLRC1,FCER1G,SYK |
| 0 | Immune system | Adaptive_NK_merged | RNPEPL1 | KLRC1,FCER1G,SYK |
| 0 | Immune system | Adaptive_NK_merged | ARHGDIA | KLRC1,FCER1G,SYK |
| 0 | Immune system | Adaptive_NK_merged | CELF2 | KLRC1,FCER1G,SYK |
| 0 | Immune system | Adaptive_NK_merged | PMAIP1 | KLRC1,FCER1G,SYK |
| 0 | Immune system | Adaptive_NK_merged | FKBP2 | KLRC1,FCER1G,SYK |
| 0 | Immune system | Adaptive_NK_merged | BANF1 | KLRC1,FCER1G,SYK |
| 0 | Immune system | Adaptive_NK_merged | PAXX | KLRC1,FCER1G,SYK |
| 0 | Immune system | Adaptive_NK_merged | TAOK3 | KLRC1,FCER1G,SYK |

| | | | | |
|---|---|---|---|---|
| 0 | Immune system | Adaptive_NK_merged | CFLAR | KLRC1,FCER1G,SYK |
| 0 | Immune system | Adaptive_NK_merged | KRT10 | KLRC1,FCER1G,SYK |
| 0 | Immune system | Adaptive_NK_merged | SELENOF | KLRC1,FCER1G,SYK |
| 0 | Immune system | Adaptive_NK_merged | HIPK1 | KLRC1,FCER1G,SYK |
| 0 | Immune system | Adaptive_NK_merged | BAZ1A | KLRC1,FCER1G,SYK |
| 0 | Immune system | Adaptive_NK_merged | UBE2Q1 | KLRC1,FCER1G,SYK |
| 0 | Immune system | Adaptive_NK_merged | TRAPPC10 | KLRC1,FCER1G,SYK |
| 0 | Immune system | Adaptive_NK_merged | MORC3 | KLRC1,FCER1G,SYK |
| 0 | Immune system | Adaptive_NK_merged | AKAP13 | KLRC1,FCER1G,SYK |
| 0 | Immune system | Adaptive_NK_merged | PHF20 | KLRC1,FCER1G,SYK |
| 0 | Immune system | Adaptive_NK_merged | EIF3K | KLRC1,FCER1G,SYK |
| 0 | Immune system | Adaptive_NK_merged | WASF2 | KLRC1,FCER1G,SYK |
| 0 | Immune system | Adaptive_NK_merged | ERBIN | KLRC1,FCER1G,SYK |
| 0 | Immune system | Adaptive_NK_merged | BZW1 | KLRC1,FCER1G,SYK |
| 0 | Immune system | Adaptive_NK_merged | LAT | KLRC1,FCER1G,SYK |
| 0 | Immune system | Adaptive_NK_merged | SERPINB6 | KLRC1,FCER1G,SYK |
| 0 | Immune system | Adaptive_NK_merged | DOCK11 | KLRC1,FCER1G,SYK |
| 0 | Immune system | Adaptive_NK_merged | C1QBP | KLRC1,FCER1G,SYK |
| 0 | Immune system | Adaptive_NK_merged | TROVE2 | KLRC1,FCER1G,SYK |
| 0 | Immune system | Adaptive_NK_merged | LINC00869 | KLRC1,FCER1G,SYK |
| 0 | Immune system | Adaptive_NK_merged | SELENOW | KLRC1,FCER1G,SYK |
| 0 | Immune system | Adaptive_NK_merged | ARRB2 | KLRC1,FCER1G,SYK |
| 0 | Immune system | Adaptive_NK_merged | TINF2 | KLRC1,FCER1G,SYK |
| 0 | Immune system | Adaptive_NK_merged | MIB2 | KLRC1,FCER1G,SYK |
| 0 | Immune system | Adaptive_NK_merged | TCP1 | KLRC1,FCER1G,SYK |
| 0 | Immune system | Adaptive_NK_merged | CMPK1 | KLRC1,FCER1G,SYK |
| 0 | Immune system | Adaptive_NK_merged | RNF125 | KLRC1,FCER1G,SYK |
| 0 | Immune system | Adaptive_NK_merged | ZBTB7A | KLRC1,FCER1G,SYK |
| 0 | Immune system | Adaptive_NK_merged | PTPN4 | KLRC1,FCER1G,SYK |
| 0 | Immune system | Adaptive_NK_merged | PRNP | KLRC1,FCER1G,SYK |
| 0 | Immune system | Adaptive_NK_merged | TADA3 | KLRC1,FCER1G,SYK |
| 0 | Immune system | Adaptive_NK_merged | FERMT3 | KLRC1,FCER1G,SYK |
| 0 | Immune system | Adaptive_NK_merged | NCR3 | KLRC1,FCER1G,SYK |
| 0 | Immune system | Adaptive_NK_merged | TMED5 | KLRC1,FCER1G,SYK |
| 0 | Immune system | Adaptive_NK_merged | GLG1 | KLRC1,FCER1G,SYK |
| 0 | Immune system | Adaptive_NK_merged | TMEM9B | KLRC1,FCER1G,SYK |
| 0 | Immune system | Adaptive_NK_merged | COX8A | KLRC1,FCER1G,SYK |
| 0 | Immune system | Adaptive_NK_merged | RAC1 | KLRC1,FCER1G,SYK |
| 0 | Immune system | Adaptive_NK_merged | SRSF9 | KLRC1,FCER1G,SYK |
| 0 | Immune system | Adaptive_NK_merged | TTC16 | KLRC1,FCER1G,SYK |
| 0 | Immune system | Adaptive_NK_merged | CADM1 | KLRC1,FCER1G,SYK |
| 0 | Immune system | Adaptive_NK_merged | SGCD | KLRC1,FCER1G,SYK |
| 0 | Immune system | Adaptive_NK_merged | DCP1B | KLRC1,FCER1G,SYK |
| 0 | Immune system | Adaptive_NK_merged | DRAXIN | KLRC1,FCER1G,SYK |
| 0 | Immune system | Adaptive_NK_merged | CAMK2N1 | KLRC1,FCER1G,SYK |
| 0 | Immune system | Adaptive_NK_merged | LOC283177 | KLRC1,FCER1G,SYK |
| 0 | Immune system | Adaptive_NK_merged | NCAPH | KLRC1,FCER1G,SYK |
| 0 | Immune system | Adaptive_NK_merged | MYO6 | KLRC1,FCER1G,SYK |
| 0 | Immune system | Adaptive_NK_merged | SBK1 | KLRC1,FCER1G,SYK |
| 0 | Immune system | Adaptive_NK_merged | CCL5 | KLRC1,FCER1G,SYK |
| 0 | Immune system | Adaptive_NK_merged | RAB11FIP5 | KLRC1,FCER1G,SYK |
| 0 | Immune system | Adaptive_NK_merged | JAKMIP1 | KLRC1,FCER1G,SYK |
| 0 | Immune system | Adaptive_NK_merged | CORO2A | KLRC1,FCER1G,SYK |
| 0 | Immune system | Adaptive_NK_merged | B3GAT1 | KLRC1,FCER1G,SYK |
| 0 | Immune system | Adaptive_NK_merged | SATB2 | KLRC1,FCER1G,SYK |
| 0 | Immune system | Adaptive_NK_merged | PDGFRB | KLRC1,FCER1G,SYK |
| 0 | Immune system | Adaptive_NK_merged | CD6 | KLRC1,FCER1G,SYK |
| 0 | Immune system | Adaptive_NK_merged | EPB41L4A | KLRC1,FCER1G,SYK |

| | | | | |
|---|---|---|---|---|
| 0 | Immune system | Adaptive_NK_merged | GDPD5 | KLRC1,FCER1G,SYK |
| 0 | Immune system | Adaptive_NK_merged | F8 | KLRC1,FCER1G,SYK |
| 0 | Immune system | Adaptive_NK_merged | LMTK3 | KLRC1,FCER1G,SYK |
| 0 | Immune system | Adaptive_NK_merged | RCAN2 | KLRC1,FCER1G,SYK |
| 0 | Immune system | Adaptive_NK_merged | GOLM1 | KLRC1,FCER1G,SYK |
| 0 | Immune system | Adaptive_NK_merged | ITPRIPL1 | KLRC1,FCER1G,SYK |
| 0 | Immune system | Adaptive_NK_merged | NUAK1 | KLRC1,FCER1G,SYK |
| 0 | Immune system | Adaptive_NK_merged | KLRAP1 | KLRC1,FCER1G,SYK |
| 0 | Immune system | Adaptive_NK_merged | TRG-AS1 | KLRC1,FCER1G,SYK |
| 0 | Immune system | Adaptive_NK_merged | PPFIA3 | KLRC1,FCER1G,SYK |
| 0 | Immune system | Adaptive_NK_merged | WNT10B | KLRC1,FCER1G,SYK |
| 0 | Immune system | Adaptive_NK_merged | TPRG1 | KLRC1,FCER1G,SYK |
| 0 | Immune system | Adaptive_NK_merged | CCDC85C | KLRC1,FCER1G,SYK |
| 0 | Immune system | Adaptive_NK_merged | LINC00944 | KLRC1,FCER1G,SYK |
| 0 | Immune system | Adaptive_NK_merged | NSG1 | KLRC1,FCER1G,SYK |
| 0 | Immune system | Adaptive_NK_merged | CDKN2A | KLRC1,FCER1G,SYK |
| 0 | Immune system | Adaptive_NK_merged | DUSP8 | KLRC1,FCER1G,SYK |
| 0 | Immune system | Adaptive_NK_merged | ACTA2 | KLRC1,FCER1G,SYK |
| 0 | Immune system | Adaptive_NK_merged | CD2 | KLRC1,FCER1G,SYK |
| 0 | Immune system | Adaptive_NK_merged | PTMS | KLRC1,FCER1G,SYK |
| 0 | Immune system | Adaptive_NK_merged | ABCD2 | KLRC1,FCER1G,SYK |
| 0 | Immune system | Adaptive_NK_merged | TP53TG1 | KLRC1,FCER1G,SYK |
| 0 | Immune system | Adaptive_NK_merged | LRRC16B | KLRC1,FCER1G,SYK |
| 0 | Immune system | Adaptive_NK_merged | KIAA1671 | KLRC1,FCER1G,SYK |
| 0 | Immune system | Adaptive_NK_merged | PAK6 | KLRC1,FCER1G,SYK |
| 0 | Immune system | Adaptive_NK_merged | MXRA7 | KLRC1,FCER1G,SYK |
| 0 | Immune system | Adaptive_NK_merged | FAM131B | KLRC1,FCER1G,SYK |
| 0 | Immune system | Adaptive_NK_merged | ATP8B2 | KLRC1,FCER1G,SYK |
| 0 | Immune system | Adaptive_NK_merged | FAM53B | KLRC1,FCER1G,SYK |
| 0 | Immune system | Adaptive_NK_merged | GLB1L2 | KLRC1,FCER1G,SYK |
| 0 | Immune system | Adaptive_NK_merged | CDC14B | KLRC1,FCER1G,SYK |
| 0 | Immune system | Adaptive_NK_merged | PCNXL2 | KLRC1,FCER1G,SYK |
| 0 | Immune system | Adaptive_NK_merged | HOXC5 | KLRC1,FCER1G,SYK |
| 0 | Immune system | Adaptive_NK_merged | MCOLN2 | KLRC1,FCER1G,SYK |
| 0 | Immune system | Adaptive_NK_merged | MVB12B | KLRC1,FCER1G,SYK |
| 0 | Immune system | Adaptive_NK_merged | ZNF365 | KLRC1,FCER1G,SYK |
| 0 | Immune system | Adaptive_NK_merged | SOX13 | KLRC1,FCER1G,SYK |
| 0 | Immune system | Adaptive_NK_merged | GCNT4 | KLRC1,FCER1G,SYK |
| 0 | Immune system | Adaptive_NK_merged | KLRC3 | KLRC1,FCER1G,SYK |
| 0 | Immune system | Adaptive_NK_merged | LAG3 | KLRC1,FCER1G,SYK |
| 0 | Immune system | Adaptive_NK_merged | TMEM255A | KLRC1,FCER1G,SYK |
| 0 | Immune system | Adaptive_NK_merged | GOLGA7B | KLRC1,FCER1G,SYK |
| 0 | Immune system | Adaptive_NK_merged | NINL | KLRC1,FCER1G,SYK |
| 0 | Immune system | Adaptive_NK_merged | DAPK2 | KLRC1,FCER1G,SYK |
| 0 | Immune system | Adaptive_NK_merged | ARHGEF28 | KLRC1,FCER1G,SYK |
| 0 | Immune system | Adaptive_NK_merged | GNAO1 | KLRC1,FCER1G,SYK |
| 0 | Immune system | Adaptive_NK_merged | PBX4 | KLRC1,FCER1G,SYK |
| 0 | Immune system | Adaptive_NK_merged | CRIP1 | KLRC1,FCER1G,SYK |
| 0 | Immune system | Adaptive_NK_merged | EPB41L4A-AS2 | KLRC1,FCER1G,SYK |
| 0 | Immune system | Adaptive_NK_merged | LINC00943 | KLRC1,FCER1G,SYK |
| 0 | Immune system | Adaptive_NK_merged | PATL2 | KLRC1,FCER1G,SYK |
| 0 | Immune system | Adaptive_NK_merged | FKBP1B | KLRC1,FCER1G,SYK |
| 0 | Immune system | Adaptive_NK_merged | MLF1 | KLRC1,FCER1G,SYK |
| 0 | Immune system | Adaptive_NK_merged | EPN2 | KLRC1,FCER1G,SYK |
| 0 | Immune system | Adaptive_NK_merged | KIF5A | KLRC1,FCER1G,SYK |
| 0 | Immune system | Adaptive_NK_merged | KCNA3 | KLRC1,FCER1G,SYK |
| 0 | Immune system | Adaptive_NK_merged | LRFN2 | KLRC1,FCER1G,SYK |
| 0 | Immune system | Adaptive_NK_merged | ISL2 | KLRC1,FCER1G,SYK |

| | | | | |
|---|---|---|---|---|
| 0 | Immune system | Adaptive_NK_merged | LIME1 | KLRC1,FCER1G,SYK |
| 0 | Immune system | Adaptive_NK_merged | GPR153 | KLRC1,FCER1G,SYK |
| 0 | Immune system | Adaptive_NK_merged | KLHL4 | KLRC1,FCER1G,SYK |
| 0 | Immune system | Adaptive_NK_merged | KIAA1324 | KLRC1,FCER1G,SYK |
| 0 | Immune system | Adaptive_NK_merged | VSTM2B | KLRC1,FCER1G,SYK |
| 0 | Immune system | Adaptive_NK_merged | CDYL2 | KLRC1,FCER1G,SYK |
| 0 | Immune system | Adaptive_NK_merged | SLC35G2 | KLRC1,FCER1G,SYK |
| 0 | Immune system | Adaptive_NK_merged | ATL1 | KLRC1,FCER1G,SYK |
| 0 | Immune system | Adaptive_NK_merged | FRMPD3 | KLRC1,FCER1G,SYK |
| 0 | Immune system | Adaptive_NK_merged | CXXC4 | KLRC1,FCER1G,SYK |
| 0 | Immune system | Adaptive_NK_merged | CLTCL1 | KLRC1,FCER1G,SYK |
| 0 | Immune system | Adaptive_NK_merged | FAM167A | KLRC1,FCER1G,SYK |
| 0 | Immune system | Adaptive_NK_merged | PPP2R2B | KLRC1,FCER1G,SYK |
| 0 | Immune system | Adaptive_NK_merged | LRFN3 | KLRC1,FCER1G,SYK |
| 0 | Immune system | Adaptive_NK_merged | HOXC4 | KLRC1,FCER1G,SYK |
| 0 | Immune system | Adaptive_NK_merged | TCERG1L | KLRC1,FCER1G,SYK |
| 0 | Immune system | Adaptive_NK_merged | C1orf177 | KLRC1,FCER1G,SYK |
| 0 | Immune system | Adaptive_NK_merged | TSHZ3 | KLRC1,FCER1G,SYK |
| 0 | Immune system | Adaptive_NK_merged | FOXD1 | KLRC1,FCER1G,SYK |
| 0 | Immune system | Adaptive_NK_merged | MYO3B | KLRC1,FCER1G,SYK |
| 0 | Immune system | Adaptive_NK_merged | TMEM244 | KLRC1,FCER1G,SYK |
| 0 | Immune system | Adaptive_NK_merged | IER5L | KLRC1,FCER1G,SYK |
| 0 | Immune system | Adaptive_NK_merged | RASGEF1A | KLRC1,FCER1G,SYK |
| 0 | Immune system | Adaptive_NK_merged | MID2 | KLRC1,FCER1G,SYK |
| 0 | Immune system | Adaptive_NK_merged | CD3D | KLRC1,FCER1G,SYK |
| 0 | Immune system | Adaptive_NK_merged | SELM | KLRC1,FCER1G,SYK |
| 0 | Immune system | Adaptive_NK_merged | TLR3 | KLRC1,FCER1G,SYK |
| 0 | Immune system | Adaptive_NK_merged | RAB6B | KLRC1,FCER1G,SYK |
| 0 | Immune system | Adaptive_NK_merged | LRRC75A | KLRC1,FCER1G,SYK |
| 0 | Immune system | Adaptive_NK_merged | MIR4435-2HG | KLRC1,FCER1G,SYK |
| 0 | Immune system | Adaptive_NK_merged | KLRC4 | KLRC1,FCER1G,SYK |
| 0 | Immune system | Adaptive_NK_merged | LTBP4 | KLRC1,FCER1G,SYK |
| 0 | Immune system | Adaptive_NK_merged | PERP | KLRC1,FCER1G,SYK |
| 0 | Immune system | Adaptive_NK_merged | SCN8A | KLRC1,FCER1G,SYK |
| 0 | Immune system | Adaptive_NK_merged | DUSP19 | KLRC1,FCER1G,SYK |
| 0 | Immune system | Adaptive_NK_merged | EDNRB-AS1 | KLRC1,FCER1G,SYK |
| 0 | Immune system | Adaptive_NK_merged | CRYBB3 | KLRC1,FCER1G,SYK |
| 0 | Immune system | Adaptive_NK_merged | SCD5 | KLRC1,FCER1G,SYK |
| 0 | Immune system | Adaptive_NK_merged | LOC102724094 | KLRC1,FCER1G,SYK |
| 0 | Immune system | Adaptive_NK_merged | ELOVL4 | KLRC1,FCER1G,SYK |
| 0 | Immune system | Adaptive_NK_merged | SPATA6L | KLRC1,FCER1G,SYK |
| 0 | Immune system | Adaptive_NK_merged | TF | KLRC1,FCER1G,SYK |
| 0 | Immune system | Adaptive_NK_merged | EFNA5 | KLRC1,FCER1G,SYK |
| 0 | Immune system | Adaptive_NK_merged | SLC45A1 | KLRC1,FCER1G,SYK |
| 0 | Immune system | Adaptive_NK_merged | CDKN2B-AS1 | KLRC1,FCER1G,SYK |
| 0 | Immune system | Adaptive_NK_merged | OTOF | KLRC1,FCER1G,SYK |
| 0 | Immune system | Adaptive_NK_merged | C1orf61 | KLRC1,FCER1G,SYK |
| 0 | Immune system | Adaptive_NK_merged | CHRNE | KLRC1,FCER1G,SYK |
| 0 | Immune system | Adaptive_NK_merged | TKTL1 | KLRC1,FCER1G,SYK |
| 0 | Immune system | Adaptive_NK_merged | TSPAN2 | KLRC1,FCER1G,SYK |
| 0 | Immune system | Adaptive_NK_merged | LOC101928988 | KLRC1,FCER1G,SYK |
| 0 | Immune system | Adaptive_NK_merged | PARD6G | KLRC1,FCER1G,SYK |
| 0 | Immune system | Adaptive_NK_merged | STXBP6 | KLRC1,FCER1G,SYK |
| 0 | Immune system | Adaptive_NK_merged | TRPC3 | KLRC1,FCER1G,SYK |
| 0 | Immune system | Adaptive_NK_merged | JAKMIP2 | KLRC1,FCER1G,SYK |
| 0 | Immune system | Adaptive_NK_merged | HEY2 | KLRC1,FCER1G,SYK |
| 0 | Immune system | Adaptive_NK_merged | TRIM46 | KLRC1,FCER1G,SYK |
| 0 | Immune system | Adaptive_NK_merged | SNPH | KLRC1,FCER1G,SYK |

| | | | | |
|---|---|---|---|---|
| 0 | Immune system | Adaptive_NK_merged | HRASLS5 | KLRC1,FCER1G,SYK |
| 0 | Immune system | Adaptive_NK_merged | SLC14A2 | KLRC1,FCER1G,SYK |
| 0 | Immune system | Adaptive_NK_merged | KRTAP5-AS1 | KLRC1,FCER1G,SYK |
| 0 | Immune system | Adaptive_NK_merged | WWTR1 | KLRC1,FCER1G,SYK |
| 0 | Immune system | Adaptive_NK_merged | SPTBN5 | KLRC1,FCER1G,SYK |
| 0 | Immune system | Adaptive_NK_merged | KIR2DL2 | KLRC1,FCER1G,SYK |
| 0 | Immune system | Adaptive_NK_merged | SLC14A1 | KLRC1,FCER1G,SYK |
| 0 | Immune system | Adaptive_NK_merged | C16orf45 | KLRC1,FCER1G,SYK |
| 0 | Immune system | Adaptive_NK_merged | TJP3 | KLRC1,FCER1G,SYK |
| 0 | Immune system | Adaptive_NK_merged | ATP1A3 | KLRC1,FCER1G,SYK |
| 0 | Immune system | Adaptive_NK_merged | ST8SIA1 | KLRC1,FCER1G,SYK |
| 0 | Immune system | Adaptive_NK_merged | AGAP1 | KLRC1,FCER1G,SYK |
| 0 | Immune system | Adaptive_NK_merged | KCCAT198 | KLRC1,FCER1G,SYK |
| 0 | Immune system | Adaptive_NK_merged | PTPRM | KLRC1,FCER1G,SYK |
| 0 | Immune system | Adaptive_NK_merged | NFIA | KLRC1,FCER1G,SYK |
| 0 | Immune system | Adaptive_NK_merged | DGKH | KLRC1,FCER1G,SYK |
| 0 | Immune system | Adaptive_NK_merged | GREB1 | KLRC1,FCER1G,SYK |
| 0 | Immune system | Adaptive_NK_merged | LINC00565 | KLRC1,FCER1G,SYK |
| 0 | Immune system | Adaptive_NK_merged | NUGGC | KLRC1,FCER1G,SYK |
| 0 | Immune system | Adaptive_NK_merged | TUBB4A | KLRC1,FCER1G,SYK |
| 0 | Immune system | Adaptive_NK_merged | IL5RA | KLRC1,FCER1G,SYK |
| 0 | Immune system | Adaptive_NK_merged | SGCB | KLRC1,FCER1G,SYK |
| 0 | Immune system | Adaptive_NK_merged | NPR3 | KLRC1,FCER1G,SYK |
| 0 | Immune system | Adaptive_NK_merged | DENND2C | KLRC1,FCER1G,SYK |
| 0 | Immune system | Adaptive_NK_merged | CD52 | KLRC1,FCER1G,SYK |
| 0 | Immune system | Adaptive_NK_merged | CDC14C | KLRC1,FCER1G,SYK |
| 0 | Immune system | Adaptive_NK_merged | MUC3A | KLRC1,FCER1G,SYK |
| 0 | Immune system | Adaptive_NK_merged | ERRFI1 | KLRC1,FCER1G,SYK |
| 0 | Immune system | Adaptive_NK_merged | LILRB1 | KLRC1,FCER1G,SYK |
| 0 | Immune system | Adaptive_NK_merged | IMPG1 | KLRC1,FCER1G,SYK |
| 0 | Immune system | Adaptive_NK_merged | GSC | KLRC1,FCER1G,SYK |
| 0 | Immune system | Adaptive_NK_merged | SCUBE3 | KLRC1,FCER1G,SYK |
| 0 | Immune system | Adaptive_NK_merged | WNT1 | KLRC1,FCER1G,SYK |
| 0 | Immune system | Adaptive_NK_merged | EPHX2 | KLRC1,FCER1G,SYK |
| 0 | Immune system | Adaptive_NK_merged | MAMLD1 | KLRC1,FCER1G,SYK |
| 0 | Immune system | Adaptive_NK_merged | NACAD | KLRC1,FCER1G,SYK |
| 0 | Immune system | Adaptive_NK_merged | LAMB1 | KLRC1,FCER1G,SYK |
| 0 | Immune system | Adaptive_NK_merged | CHSY3 | KLRC1,FCER1G,SYK |
| 0 | Immune system | Adaptive_NK_merged | SHB | KLRC1,FCER1G,SYK |
| 0 | Immune system | Adaptive_NK_merged | ADAMTSL5 | KLRC1,FCER1G,SYK |
| 0 | Immune system | Adaptive_NK_merged | LOC101929719 | KLRC1,FCER1G,SYK |
| 0 | Immune system | Adaptive_NK_merged | TRPC1 | KLRC1,FCER1G,SYK |
| 0 | Immune system | Adaptive_NK_merged | IL32 | KLRC1,FCER1G,SYK |
| 0 | Immune system | Adaptive_NK_merged | ZFP36 | KLRC1,FCER1G,SYK |
| 0 | Immune system | Adaptive_NK_merged | WDR74 | KLRC1,FCER1G,SYK |
| 0 | Immune system | Adaptive_NK_merged | CD160 | KLRC1,FCER1G,SYK |
| 0 | Immune system | Adaptive_NK_merged | XCL2 | KLRC1,FCER1G,SYK |
| 0 | Immune system | Adaptive_NK_merged | ID2 | KLRC1,FCER1G,SYK |
| 0 | Immune system | Adaptive_NK_merged | IER2 | KLRC1,FCER1G,SYK |
| 0 | Immune system | Adaptive_NK_merged | CD7 | KLRC1,FCER1G,SYK |
| 0 | Immune system | Adaptive_NK_merged | SNORD3A | KLRC1,FCER1G,SYK |
| 0 | Immune system | Adaptive_NK_merged | CCL3 | KLRC1,FCER1G,SYK |
| 0 | Immune system | Adaptive_NK_merged | TMEM107 | KLRC1,FCER1G,SYK |
| 0 | Immune system | Adaptive_NK_merged | FOS | KLRC1,FCER1G,SYK |
| 0 | Immune system | Adaptive_NK_merged | XCL1 | KLRC1,FCER1G,SYK |
| 0 | Immune system | Adaptive_NK_merged | SNORD3D | KLRC1,FCER1G,SYK |
| 0 | Immune system | Adaptive_NK_merged | MAP3K8 | KLRC1,FCER1G,SYK |
| 0 | Immune system | Adaptive_NK_merged | JUN | KLRC1,FCER1G,SYK |

| | | | | |
|---|---|---|---|---|
| 0 | Immune system | Adaptive_NK_merged | PPP1R15A | KLRC1,FCER1G,SYK |
| 0 | Immune system | Adaptive_NK_merged | VIM | KLRC1,FCER1G,SYK |
| 0 | Immune system | Adaptive_NK_merged | RNU12 | KLRC1,FCER1G,SYK |
| 0 | Immune system | Adaptive_NK_merged | SNORD3B-1 | KLRC1,FCER1G,SYK |
| 0 | Immune system | Adaptive_NK_merged | IL2RB | KLRC1,FCER1G,SYK |
| 0 | Immune system | Adaptive_NK_merged | TMIGD2 | KLRC1,FCER1G,SYK |
| 0 | Immune system | Adaptive_NK_merged | RP11-386I14.4 | KLRC1,FCER1G,SYK |
| 0 | Immune system | Adaptive_NK_merged | GZMK | KLRC1,FCER1G,SYK |
| 0 | Immune system | Adaptive_NK_merged | LTB | KLRC1,FCER1G,SYK |
| 1 | Immune system | Conventional NK | TCF4 | KLRC2 |
| 1 | Immune system | Conventional NK | IL18RAP | KLRC2 |
| 1 | Immune system | Conventional NK | CD79A | KLRC2 |
| 1 | Immune system | Conventional NK | CABLES1 | KLRC2 |
| 1 | Immune system | Conventional NK | KIT | KLRC2 |
| 1 | Immune system | Conventional NK | ZBTB16 | KLRC2 |
| 1 | Immune system | Conventional NK | POU2AF1 | KLRC2 |
| 1 | Immune system | Conventional NK | ORAI2 | KLRC2 |
| 1 | Immune system | Conventional NK | CYSLTR1 | KLRC2 |
| 1 | Immune system | Conventional NK | RUNX2 | KLRC2 |
| 1 | Immune system | Conventional NK | P2RX1 | KLRC2 |
| 1 | Immune system | Conventional NK | RTKN | KLRC2 |
| 1 | Immune system | Conventional NK | KLRC1 | KLRC2 |
| 1 | Immune system | Conventional NK | SCRN1 | KLRC2 |
| 1 | Immune system | Conventional NK | MFGE8 | KLRC2 |
| 1 | Immune system | Conventional NK | BLNK | KLRC2 |
| 1 | Immune system | Conventional NK | CD83 | KLRC2 |
| 1 | Immune system | Conventional NK | NAPSB | KLRC2 |
| 1 | Immune system | Conventional NK | GSN | KLRC2 |
| 1 | Immune system | Conventional NK | LINC00926 | KLRC2 |
| 1 | Immune system | Conventional NK | ADGRG3 | KLRC2 |
| 1 | Immune system | Conventional NK | IGF2BP2 | KLRC2 |
| 1 | Immune system | Conventional NK | APP | KLRC2 |
| 1 | Immune system | Conventional NK | FCRLA | KLRC2 |
| 1 | Immune system | Conventional NK | JCHAIN | KLRC2 |
| 1 | Immune system | Conventional NK | MEF2C | KLRC2 |
| 1 | Immune system | Conventional NK | SLC9A7 | KLRC2 |
| 1 | Immune system | Conventional NK | GNG7 | KLRC2 |
| 1 | Immune system | Conventional NK | FHL3 | KLRC2 |
| 1 | Immune system | Conventional NK | LGALS9B | KLRC2 |
| 1 | Immune system | Conventional NK | PTK2 | KLRC2 |
| 1 | Immune system | Conventional NK | BANK1 | KLRC2 |
| 1 | Immune system | Conventional NK | HVCN1 | KLRC2 |
| 1 | Immune system | Conventional NK | TNFRSF13C | KLRC2 |
| 1 | Immune system | Conventional NK | HDAC9 | KLRC2 |
| 1 | Immune system | Conventional NK | CD24 | KLRC2 |
| 1 | Immune system | Conventional NK | BTBD3 | KLRC2 |
| 1 | Immune system | Conventional NK | BAALC | KLRC2 |
| 1 | Immune system | Conventional NK | BTK | KLRC2 |
| 1 | Immune system | Conventional NK | MIR378I | KLRC2 |
| 1 | Immune system | Conventional NK | GAB1 | KLRC2 |
| 1 | Immune system | Conventional NK | FAM129C | KLRC2 |
| 1 | Immune system | Conventional NK | GPR55 | KLRC2 |
| 1 | Immune system | Conventional NK | TRIB2 | KLRC2 |
| 1 | Immune system | Conventional NK | TLR10 | KLRC2 |
| 1 | Immune system | Conventional NK | GYLTL1B | KLRC2 |
| 1 | Immune system | Conventional NK | CD79B | KLRC2 |
| 1 | Immune system | Conventional NK | DAPP1 | KLRC2 |
| 1 | Immune system | Conventional NK | ZNF418 | KLRC2 |

| | | | | |
|---|---|---|---|---|
| 1 | Immune system | Conventional NK | SOCS6 | KLRC2 |
| 1 | Immune system | Conventional NK | KLHL6 | KLRC2 |
| 1 | Immune system | Conventional NK | CD19 | KLRC2 |
| 1 | Immune system | Conventional NK | LINC00996 | KLRC2 |
| 1 | Immune system | Conventional NK | BLK | KLRC2 |
| 1 | Immune system | Conventional NK | PDE7B | KLRC2 |
| 1 | Immune system | Conventional NK | MS4A1 | KLRC2 |
| 1 | Immune system | Conventional NK | STAP1 | KLRC2 |
| 1 | Immune system | Conventional NK | KIAA0754 | KLRC2 |
| 1 | Immune system | Conventional NK | TOX2 | KLRC2 |
| 1 | Immune system | Conventional NK | TCL1A | KLRC2 |
| 1 | Immune system | Conventional NK | MIR4539 | KLRC2 |
| 1 | Immune system | Conventional NK | TTC9 | KLRC2 |
| 1 | Immune system | Conventional NK | LTB | KLRC2 |
| 1 | Immune system | Conventional NK | NCF4 | KLRC2 |
| 1 | Immune system | Conventional NK | CD200 | KLRC2 |
| 1 | Immune system | Conventional NK | FCRL2 | KLRC2 |
| 1 | Immune system | Conventional NK | AEBP1 | KLRC2 |
| 1 | Immune system | Conventional NK | SPIB | KLRC2 |
| 1 | Immune system | Conventional NK | INF2 | KLRC2 |
| 1 | Immune system | Conventional NK | ITM2C | KLRC2 |
| 1 | Immune system | Conventional NK | MICU3 | KLRC2 |
| 1 | Immune system | Conventional NK | AFF3 | KLRC2 |
| 1 | Immune system | Conventional NK | CD22 | KLRC2 |
| 1 | Immune system | Conventional NK | ZNF704 | KLRC2 |
| 1 | Immune system | Conventional NK | MIR4537 | KLRC2 |
| 1 | Immune system | Conventional NK | EBF1 | KLRC2 |
| 1 | Immune system | Conventional NK | DERL3 | KLRC2 |
| 1 | Immune system | Conventional NK | IL1R1 | KLRC2 |
| 1 | Immune system | Conventional NK | IRS2 | KLRC2 |
| 1 | Immune system | Conventional NK | IL12RB2 | KLRC2 |
| 1 | Immune system | Conventional NK | CPNE5 | KLRC2 |
| 1 | Immune system | Conventional NK | ZNF135 | KLRC2 |
| 1 | Immune system | Conventional NK | GAPT | KLRC2 |
| 1 | Immune system | Conventional NK | CCR7 | KLRC2 |
| 1 | Immune system | Conventional NK | CALCRL | KLRC2 |
| 1 | Immune system | Conventional NK | SPINT2 | KLRC2 |
| 1 | Immune system | Conventional NK | SNN | KLRC2 |
| 1 | Immune system | Conventional NK | RFX2 | KLRC2 |
| 1 | Immune system | Conventional NK | MAML3 | KLRC2 |
| 1 | Immune system | Conventional NK | FZD6 | KLRC2 |
| 1 | Immune system | Conventional NK | PVR | KLRC2 |
| 1 | Immune system | Conventional NK | FAM111B | KLRC2 |
| 1 | Immune system | Conventional NK | TSPAN13 | KLRC2 |
| 1 | Immune system | Conventional NK | ICOSLG | KLRC2 |
| 1 | Immune system | Conventional NK | C6orf25 | KLRC2 |
| 1 | Immune system | Conventional NK | GRASP | KLRC2 |
| 1 | Immune system | Conventional NK | FCER1G | KLRC2 |
| 1 | Immune system | Conventional NK | BCL7A | KLRC2 |
| 1 | Immune system | Conventional NK | LY86 | KLRC2 |
| 1 | Immune system | Conventional NK | LAD1 | KLRC2 |
| 1 | Immune system | Conventional NK | BTLA | KLRC2 |
| 1 | Immune system | Conventional NK | CDK14 | KLRC2 |
| 1 | Immune system | Conventional NK | IL18 | KLRC2 |
| 1 | Immune system | Conventional NK | KCNN4 | KLRC2 |
| 1 | Immune system | Conventional NK | RGS1 | KLRC2 |
| 1 | Immune system | Conventional NK | AATK | KLRC2 |
| 1 | Immune system | Conventional NK | EBI3 | KLRC2 |

| | | | | |
|---|---|---|---|---|
| 1 | Immune system | Conventional NK | PHACTR1 | KLRC2 |
| 1 | Immune system | Conventional NK | IGFBP4 | KLRC2 |
| 1 | Immune system | Conventional NK | TGM2 | KLRC2 |
| 1 | Immune system | Conventional NK | ADAM28 | KLRC2 |
| 1 | Immune system | Conventional NK | LAPTM4B | KLRC2 |
| 1 | Immune system | Conventional NK | POU2F2 | KLRC2 |
| 1 | Immune system | Conventional NK | ANKRD50 | KLRC2 |
| 1 | Immune system | Conventional NK | KIAA0125 | KLRC2 |
| 1 | Immune system | Conventional NK | JUP | KLRC2 |
| 1 | Immune system | Conventional NK | ZNF154 | KLRC2 |
| 1 | Immune system | Conventional NK | PLD4 | KLRC2 |
| 1 | Immune system | Conventional NK | ANKRD33B | KLRC2 |
| 1 | Immune system | Conventional NK | MYO1E | KLRC2 |
| 1 | Immune system | Conventional NK | GUCY1A3 | KLRC2 |
| 1 | Immune system | Conventional NK | TIE1 | KLRC2 |
| 1 | Immune system | Conventional NK | RAB31 | KLRC2 |
| 1 | Immune system | Conventional NK | HLA-DOA | KLRC2 |
| 1 | Immune system | Conventional NK | C19orf38 | KLRC2 |
| 1 | Immune system | Conventional NK | SERPINE2 | KLRC2 |
| 1 | Immune system | Conventional NK | TRIB1 | KLRC2 |
| 1 | Immune system | Conventional NK | ZNF629 | KLRC2 |
| 1 | Immune system | Conventional NK | BMP2 | KLRC2 |
| 1 | Immune system | Conventional NK | SNX8 | KLRC2 |
| 1 | Immune system | Conventional NK | ALDH2 | KLRC2 |
| 1 | Immune system | Conventional NK | LDB2 | KLRC2 |
| 1 | Immune system | Conventional NK | LRRK2 | KLRC2 |
| 1 | Immune system | Conventional NK | RALGPS2 | KLRC2 |
| 1 | Immune system | Conventional NK | PAX5 | KLRC2 |
| 1 | Immune system | Conventional NK | FES | KLRC2 |
| 1 | Immune system | Conventional NK | WDFY4 | KLRC2 |
| 1 | Immune system | Conventional NK | CHML | KLRC2 |
| 1 | Immune system | Conventional NK | ZNF814 | KLRC2 |
| 1 | Immune system | Conventional NK | CLEC17A | KLRC2 |
| 1 | Immune system | Conventional NK | TNFRSF25 | KLRC2 |
| 1 | Immune system | Conventional NK | KIAA0226L | KLRC2 |
| 1 | Immune system | Conventional NK | FLNB | KLRC2 |
| 1 | Immune system | Conventional NK | CDCA7L | KLRC2 |
| 1 | Immune system | Conventional NK | OSBPL10 | KLRC2 |
| 1 | Immune system | Conventional NK | TBC1D9 | KLRC2 |
| 1 | Immune system | Conventional NK | TTC28 | KLRC2 |
| 1 | Immune system | Conventional NK | SCML1 | KLRC2 |
| 1 | Immune system | Conventional NK | SLC4A10 | KLRC2 |
| 1 | Immune system | Conventional NK | FAM63A | KLRC2 |
| 1 | Immune system | Conventional NK | ALPK1 | KLRC2 |
| 1 | Immune system | Conventional NK | JADE3 | KLRC2 |
| 1 | Immune system | Conventional NK | CLIC4 | KLRC2 |
| 1 | Immune system | Conventional NK | TNFSF11 | KLRC2 |
| 1 | Immune system | Conventional NK | TNFRSF11A | KLRC2 |
| 1 | Immune system | Conventional NK | TSPAN33 | KLRC2 |
| 1 | Immune system | Conventional NK | XYLT1 | KLRC2 |
| 1 | Immune system | Conventional NK | CCR6 | KLRC2 |
| 1 | Immune system | Conventional NK | LGALS3BP | KLRC2 |
| 1 | Immune system | Conventional NK | DPP4 | KLRC2 |
| 1 | Immune system | Conventional NK | GNB4 | KLRC2 |
| 1 | Immune system | Conventional NK | LGALS9C | KLRC2 |
| 1 | Immune system | Conventional NK | PDE6G | KLRC2 |
| 1 | Immune system | Conventional NK | CD180 | KLRC2 |
| 1 | Immune system | Conventional NK | CCL22 | KLRC2 |

| | | | | |
|---|---|---|---|---|
| 1 | Immune system | Conventional NK | SRGAP3 | KLRC2 |
| 1 | Immune system | Conventional NK | LRRK1 | KLRC2 |
| 1 | Immune system | Conventional NK | KIF7 | KLRC2 |
| 1 | Immune system | Conventional NK | AHR | KLRC2 |
| 1 | Immune system | Conventional NK | RENBP | KLRC2 |
| 1 | Immune system | Conventional NK | ITGAE | KLRC2 |
| 1 | Immune system | Conventional NK | CR2 | KLRC2 |
| 1 | Immune system | Conventional NK | HLA-DRA | KLRC2 |
| 1 | Immune system | Conventional NK | GEN1 | KLRC2 |
| 1 | Immune system | Conventional NK | COL9A3 | KLRC2 |
| 1 | Immune system | Conventional NK | ZSCAN18 | KLRC2 |
| 1 | Immune system | Conventional NK | TBC1D8 | KLRC2 |
| 1 | Immune system | Conventional NK | TESPA1 | KLRC2 |
| 1 | Immune system | Conventional NK | SLC7A11 | KLRC2 |
| 1 | Immune system | Conventional NK | KRT73 | KLRC2 |
| 1 | Immune system | Conventional NK | TNFAIP8 | KLRC2 |
| 1 | Immune system | Conventional NK | CXorf21 | KLRC2 |
| 1 | Immune system | Conventional NK | PECAM1 | KLRC2 |
| 1 | Immune system | Conventional NK | ZNF469 | KLRC2 |
| 1 | Immune system | Conventional NK | GSG2 | KLRC2 |
| 1 | Immune system | Conventional NK | RASSF2 | KLRC2 |
| 1 | Immune system | Conventional NK | ZNF667-AS1 | KLRC2 |
| 1 | Immune system | Conventional NK | DLL1 | KLRC2 |
| 1 | Immune system | Conventional NK | CNN3 | KLRC2 |
| 1 | Immune system | Conventional NK | RNF144B | KLRC2 |
| 1 | Immune system | Conventional NK | CERS6 | KLRC2 |
| 1 | Immune system | Conventional NK | MMRN1 | KLRC2 |
| 1 | Immune system | Conventional NK | KCNC3 | KLRC2 |
| 1 | Immune system | Conventional NK | TFPI | KLRC2 |
| 1 | Immune system | Conventional NK | USP6NL | KLRC2 |
| 1 | Immune system | Conventional NK | HLX | KLRC2 |
| 1 | Immune system | Conventional NK | ZBTB18 | KLRC2 |
| 1 | Immune system | Conventional NK | CLMN | KLRC2 |
| 1 | Immune system | Conventional NK | ALCAM | KLRC2 |
| 1 | Immune system | Conventional NK | TRAF4 | KLRC2 |
| 1 | Immune system | Conventional NK | KCTD3 | KLRC2 |
| 1 | Immune system | Conventional NK | RNF217 | KLRC2 |
| 1 | Immune system | Conventional NK | C16orf74 | KLRC2 |
| 1 | Immune system | Conventional NK | SWAP70 | KLRC2 |
| 1 | Immune system | Conventional NK | MGLL | KLRC2 |
| 1 | Immune system | Conventional NK | TNFRSF13B | KLRC2 |
| 1 | Immune system | Conventional NK | LINC01215 | KLRC2 |
| 1 | Immune system | Conventional NK | CD40 | KLRC2 |
| 1 | Immune system | Conventional NK | GNG11 | KLRC2 |
| 1 | Immune system | Conventional NK | HLA-DMB | KLRC2 |
| 1 | Immune system | Conventional NK | KRT72 | KLRC2 |
| 1 | Immune system | Conventional NK | MS4A7 | KLRC2 |
| 1 | Immune system | Conventional NK | IFNGR2 | KLRC2 |
| 1 | Immune system | Conventional NK | RUSC2 | KLRC2 |
| 1 | Immune system | Conventional NK | ADRBK2 | KLRC2 |
| 1 | Immune system | Conventional NK | C1orf162 | KLRC2 |
| 1 | Immune system | Conventional NK | SPTSSB | KLRC2 |
| 1 | Immune system | Conventional NK | PLXNB2 | KLRC2 |
| 1 | Immune system | Conventional NK | LRRC32 | KLRC2 |
| 1 | Immune system | Conventional NK | ID3 | KLRC2 |
| 1 | Immune system | Conventional NK | B3GALNT1 | KLRC2 |
| 1 | Immune system | Conventional NK | SPTLC3 | KLRC2 |
| 1 | Immune system | Conventional NK | DENND5B | KLRC2 |

| | | | | |
|---|---|---|---|---|
| 1 | Immune system | Conventional NK | GPR183 | KLRC2 |
| 1 | Immune system | Conventional NK | SLC2A5 | KLRC2 |
| 1 | Immune system | Conventional NK | WNT10A | KLRC2 |
| 1 | Immune system | Conventional NK | CD1C | KLRC2 |
| 1 | Immune system | Conventional NK | PTK7 | KLRC2 |
| 1 | Immune system | Conventional NK | PTGIR | KLRC2 |
| 1 | Immune system | Conventional NK | SIT1 | KLRC2 |
| 1 | Immune system | Conventional NK | EMILIN2 | KLRC2 |
| 1 | Immune system | Conventional NK | BCL11A | KLRC2 |
| 1 | Immune system | Conventional NK | KYNU | KLRC2 |
| 1 | Immune system | Conventional NK | ARNTL2 | KLRC2 |
| 1 | Immune system | Conventional NK | BASP1 | KLRC2 |
| 1 | Immune system | Conventional NK | ADAMTS7 | KLRC2 |
| 1 | Immune system | Conventional NK | SH3PXD2A | KLRC2 |
| 1 | Immune system | Conventional NK | MACROD2 | KLRC2 |
| 1 | Immune system | Conventional NK | VPREB3 | KLRC2 |
| 1 | Immune system | Conventional NK | NFIX | KLRC2 |
| 1 | Immune system | Conventional NK | MARCKS | KLRC2 |
| 1 | Immune system | Conventional NK | SNCA | KLRC2 |
| 1 | Immune system | Conventional NK | FGD2 | KLRC2 |
| 1 | Immune system | Conventional NK | LAMC1 | KLRC2 |
| 1 | Immune system | Conventional NK | THBS1 | KLRC2 |
| 1 | Immune system | Conventional NK | ELL3 | KLRC2 |
| 1 | Immune system | Conventional NK | SIK1 | KLRC2 |
| 1 | Immune system | Conventional NK | 3-Mar | KLRC2 |
| 1 | Immune system | Conventional NK | ZNF285 | KLRC2 |
| 1 | Immune system | Conventional NK | LOC151174 | KLRC2 |
| 1 | Immune system | Conventional NK | ZIK1 | KLRC2 |
| 1 | Immune system | Conventional NK | RMI2 | KLRC2 |
| 1 | Immune system | Conventional NK | ZNF415 | KLRC2 |
| 1 | Immune system | Conventional NK | SEMA7A | KLRC2 |
| 1 | Immune system | Conventional NK | MARCKSL1 | KLRC2 |
| 1 | Immune system | Conventional NK | CD27 | KLRC2 |
| 1 | Immune system | Conventional NK | PTGS1 | KLRC2 |
| 1 | Immune system | Conventional NK | NEIL1 | KLRC2 |
| 1 | Immune system | Conventional NK | CXCR5 | KLRC2 |
| 1 | Immune system | Conventional NK | MIR4538 | KLRC2 |
| 1 | Immune system | Conventional NK | CBFA2T3 | KLRC2 |
| 1 | Immune system | Conventional NK | LINC00494 | KLRC2 |
| 1 | Immune system | Conventional NK | DTX4 | KLRC2 |
| 1 | Immune system | Conventional NK | FSCN1 | KLRC2 |
| 1 | Immune system | Conventional NK | PYCR1 | KLRC2 |
| 1 | Immune system | Conventional NK | TIMP2 | KLRC2 |
| 1 | Immune system | Conventional NK | PKIG | KLRC2 |
| 1 | Immune system | Conventional NK | TRAF3IP2 | KLRC2 |
| 1 | Immune system | Conventional NK | ADAM19 | KLRC2 |
| 1 | Immune system | Conventional NK | GPR15 | KLRC2 |
| 1 | Immune system | Conventional NK | LOC100129034 | KLRC2 |
| 1 | Immune system | Conventional NK | TREML2 | KLRC2 |
| 1 | Immune system | Conventional NK | CCDC106 | KLRC2 |
| 1 | Immune system | Conventional NK | CTSH | KLRC2 |
| 1 | Immune system | Conventional NK | HOXA3 | KLRC2 |
| 1 | Immune system | Conventional NK | GUCY1B3 | KLRC2 |
| 1 | Immune system | Conventional NK | ADGRA2 | KLRC2 |
| 1 | Immune system | Conventional NK | COL19A1 | KLRC2 |
| 1 | Immune system | Conventional NK | NDST1 | KLRC2 |
| 1 | Immune system | Conventional NK | PTPRO | KLRC2 |
| 1 | Immune system | Conventional NK | LOC100506258 | KLRC2 |

| | | | | |
|---|---|---|---|---|
| 1 | Immune system | Conventional NK | BTBD11 | KLRC2 |
| 1 | Immune system | Conventional NK | ZNF223 | KLRC2 |
| 1 | Immune system | Conventional NK | MB21D2 | KLRC2 |
| 1 | Immune system | Conventional NK | RGS18 | KLRC2 |
| 1 | Immune system | Conventional NK | UBE2E2 | KLRC2 |
| 1 | Immune system | Conventional NK | PAG1 | KLRC2 |
| 1 | Immune system | Conventional NK | CTTNBP2NL | KLRC2 |
| 1 | Immune system | Conventional NK | SPRED2 | KLRC2 |
| 1 | Immune system | Conventional NK | MCTP1 | KLRC2 |
| 1 | Immune system | Conventional NK | SLAMF1 | KLRC2 |
| 1 | Immune system | Conventional NK | GRAP | KLRC2 |
| 1 | Immune system | Conventional NK | LRRC43 | KLRC2 |
| 1 | Immune system | Conventional NK | NEK8 | KLRC2 |
| 1 | Immune system | Conventional NK | AMICA1 | KLRC2 |
| 1 | Immune system | Conventional NK | ATP8B4 | KLRC2 |
| 1 | Immune system | Conventional NK | PCDH9 | KLRC2 |
| 1 | Immune system | Conventional NK | SOX4 | KLRC2 |
| 1 | Immune system | Conventional NK | GPX7 | KLRC2 |
| 1 | Immune system | Conventional NK | ZNF772 | KLRC2 |
| 1 | Immune system | Conventional NK | MGAT3 | KLRC2 |
| 1 | Immune system | Conventional NK | EREG | KLRC2 |
| 1 | Immune system | Conventional NK | SOGA1 | KLRC2 |
| 1 | Immune system | Conventional NK | IRAK3 | KLRC2 |
| 1 | Immune system | Conventional NK | HSBP1L1 | KLRC2 |
| 1 | Immune system | Conventional NK | BEND4 | KLRC2 |
| 1 | Immune system | Conventional NK | WFDC21P | KLRC2 |
| 1 | Immune system | Conventional NK | KMO | KLRC2 |
| 1 | Immune system | Conventional NK | FCRL1 | KLRC2 |
| 1 | Immune system | Conventional NK | LOC102724323 | KLRC2 |
| 1 | Immune system | Conventional NK | ZNF835 | KLRC2 |
| 1 | Immune system | Conventional NK | PLEKHO1 | KLRC2 |
| 1 | Immune system | Conventional NK | OCRL | KLRC2 |
| 1 | Immune system | Conventional NK | PPBP | KLRC2 |
| 1 | Immune system | Conventional NK | PAWR | KLRC2 |
| 1 | Immune system | Conventional NK | FSD1 | KLRC2 |
| 1 | Immune system | Conventional NK | CD300C | KLRC2 |
| 1 | Immune system | Conventional NK | ZNF404 | KLRC2 |
| 1 | Immune system | Conventional NK | ZNF608 | KLRC2 |
| 1 | Immune system | Conventional NK | NOXA1 | KLRC2 |
| 1 | Immune system | Conventional NK | IRAK2 | KLRC2 |
| 1 | Immune system | Conventional NK | ZDHHC14 | KLRC2 |
| 1 | Immune system | Conventional NK | MIR631 | KLRC2 |
| 1 | Immune system | Conventional NK | LCN10 | KLRC2 |
| 1 | Immune system | Conventional NK | 1-Mar | KLRC2 |
| 1 | Immune system | Conventional NK | C1orf186 | KLRC2 |
| 1 | Immune system | Conventional NK | ARSJ | KLRC2 |
| 1 | Immune system | Conventional NK | P2RY1 | KLRC2 |
| 1 | Immune system | Conventional NK | PPP1R26 | KLRC2 |
| 1 | Immune system | Conventional NK | IL6R | KLRC2 |
| 1 | Immune system | Conventional NK | RAB32 | KLRC2 |
| 1 | Immune system | Conventional NK | ZNF660 | KLRC2 |
| 1 | Immune system | Conventional NK | TPTEP1 | KLRC2 |
| 1 | Immune system | Conventional NK | ZNF256 | KLRC2 |
| 1 | Immune system | Conventional NK | NEK6 | KLRC2 |
| 1 | Immune system | Conventional NK | LRP5 | KLRC2 |
| 1 | Immune system | Conventional NK | ATP6V0A1 | KLRC2 |
| 1 | Immune system | Conventional NK | C9orf91 | KLRC2 |
| 1 | Immune system | Conventional NK | PSD3 | KLRC2 |

| | | | | |
|---|---|---|---|---|
| 1 | Immune system | Conventional NK | IL13RA1 | KLRC2 |
| 1 | Immune system | Conventional NK | SLCO4A1 | KLRC2 |
| 1 | Immune system | Conventional NK | LOC389641 | KLRC2 |
| 1 | Immune system | Conventional NK | RERE | KLRC2 |
| 1 | Immune system | Conventional NK | ALOX5 | KLRC2 |
| 1 | Immune system | Conventional NK | ZNF385A | KLRC2 |
| 1 | Immune system | Conventional NK | GNG8 | KLRC2 |
| 1 | Immune system | Conventional NK | TTYH3 | KLRC2 |
| 1 | Immune system | Conventional NK | CDCA7 | KLRC2 |
| 1 | Immune system | Conventional NK | RGMB | KLRC2 |
| 1 | Immune system | Conventional NK | TBC1D12 | KLRC2 |
| 1 | Immune system | Conventional NK | CELSR1 | KLRC2 |
| 1 | Immune system | Conventional NK | MAFG | KLRC2 |
| 1 | Immune system | Conventional NK | SORT1 | KLRC2 |
| 1 | Immune system | Conventional NK | AMZ1 | KLRC2 |
| 1 | Immune system | Conventional NK | SQLE | KLRC2 |
| 1 | Immune system | Conventional NK | FCER2 | KLRC2 |
| 1 | Immune system | Conventional NK | COTL1 | KLRC2 |
| 1 | Immune system | Conventional NK | SPI1 | KLRC2 |
| 1 | Immune system | Conventional NK | CEBPD | KLRC2 |
| 1 | Immune system | Conventional NK | P2RY14 | KLRC2 |
| 1 | Immune system | Conventional NK | CPNE2 | KLRC2 |
| 1 | Immune system | Conventional NK | HLA-DRB1 | KLRC2 |
| 1 | Immune system | Conventional NK | MPEG1 | KLRC2 |
| 1 | Immune system | Conventional NK | TRIQK | KLRC2 |
| 1 | Immune system | Conventional NK | TUBB6 | KLRC2 |
| 1 | Immune system | Conventional NK | SYPL1 | KLRC2 |
| 1 | Immune system | Conventional NK | LINC01480 | KLRC2 |
| 1 | Immune system | Conventional NK | ZNF516 | KLRC2 |
| 1 | Immune system | Conventional NK | F13A1 | KLRC2 |
| 1 | Immune system | Conventional NK | GNA15 | KLRC2 |
| 1 | Immune system | Conventional NK | SIGLEC6 | KLRC2 |
| 1 | Immune system | Conventional NK | ATF7IP2 | KLRC2 |
| 1 | Immune system | Conventional NK | DPPA4 | KLRC2 |
| 1 | Immune system | Conventional NK | CDC42BPB | KLRC2 |
| 1 | Immune system | Conventional NK | COL9A2 | KLRC2 |
| 1 | Immune system | Conventional NK | DCANP1 | KLRC2 |
| 1 | Immune system | Conventional NK | MZB1 | KLRC2 |
| 1 | Immune system | Conventional NK | RGS16 | KLRC2 |
| 1 | Immune system | Conventional NK | CSF2RB | KLRC2 |
| 1 | Immune system | Conventional NK | CYFIP1 | KLRC2 |
| 1 | Immune system | Conventional NK | RBM47 | KLRC2 |
| 1 | Immune system | Conventional NK | LHFPL2 | KLRC2 |
| 1 | Immune system | Conventional NK | LOC100506985 | KLRC2 |
| 1 | Immune system | Conventional NK | HLA-DQA1 | KLRC2 |
| 1 | Immune system | Conventional NK | CFP | KLRC2 |
| 1 | Immune system | Conventional NK | SLC37A2 | KLRC2 |
| 1 | Immune system | Conventional NK | PVT1 | KLRC2 |
| 1 | Immune system | Conventional NK | TP53I3 | KLRC2 |
| 1 | Immune system | Conventional NK | MYC | KLRC2 |
| 1 | Immune system | Conventional NK | ARMC9 | KLRC2 |
| 1 | Immune system | Conventional NK | DST | KLRC2 |
| 1 | Immune system | Conventional NK | KIF13A | KLRC2 |
| 1 | Immune system | Conventional NK | LOC646762 | KLRC2 |
| 1 | Immune system | Conventional NK | P2RX5 | KLRC2 |
| 1 | Immune system | Conventional NK | ADGRE2 | KLRC2 |
| 1 | Immune system | Conventional NK | BHLHE41 | KLRC2 |
| 1 | Immune system | Conventional NK | GIPC3 | KLRC2 |

| | | | | |
|---|---|---|---|---|
| 1 | Immune system | Conventional NK | ZNF667 | KLRC2 |
| 1 | Immune system | Conventional NK | STARD4-AS1 | KLRC2 |
| 1 | Immune system | Conventional NK | ADGRA3 | KLRC2 |
| 1 | Immune system | Conventional NK | HCK | KLRC2 |
| 1 | Immune system | Conventional NK | CD55 | KLRC2 |
| 1 | Immune system | Conventional NK | FAM69B | KLRC2 |
| 1 | Immune system | Conventional NK | MPZL2 | KLRC2 |
| 1 | Immune system | Conventional NK | BCAR3 | KLRC2 |
| 1 | Immune system | Conventional NK | CARD6 | KLRC2 |
| 1 | Immune system | Conventional NK | PRICKLE1 | KLRC2 |
| 1 | Immune system | Conventional NK | RAB13 | KLRC2 |
| 1 | Immune system | Conventional NK | BEX4 | KLRC2 |
| 1 | Immune system | Conventional NK | NGFRAP1 | KLRC2 |
| 1 | Immune system | Conventional NK | GAS6 | KLRC2 |
| 1 | Immune system | Conventional NK | DSE | KLRC2 |
| 1 | Immune system | Conventional NK | RLTPR | KLRC2 |
| 1 | Immune system | Conventional NK | CEACAM1 | KLRC2 |
| 1 | Immune system | Conventional NK | HHEX | KLRC2 |
| 1 | Immune system | Conventional NK | RASAL1 | KLRC2 |
| 1 | Immune system | Conventional NK | GNAQ | KLRC2 |
| 1 | Immune system | Conventional NK | ACKR2 | KLRC2 |
| 1 | Immune system | Conventional NK | MAP10 | KLRC2 |
| 1 | Immune system | Conventional NK | PTGFRN | KLRC2 |
| 1 | Immune system | Conventional NK | SMOX | KLRC2 |
| 1 | Immune system | Conventional NK | VMO1 | KLRC2 |
| 1 | Immune system | Conventional NK | DOK3 | KLRC2 |
| 1 | Immune system | Conventional NK | DUSP6 | KLRC2 |
| 1 | Immune system | Conventional NK | NR4A3 | KLRC2 |
| 1 | Immune system | Conventional NK | SRC | KLRC2 |
| 1 | Immune system | Conventional NK | CD300LF | KLRC2 |
| 1 | Immune system | Conventional NK | PF4 | KLRC2 |
| 1 | Immune system | Conventional NK | SYNGR2 | KLRC2 |
| 1 | Immune system | Conventional NK | GATM | KLRC2 |
| 1 | Immune system | Conventional NK | CYGB | KLRC2 |
| 1 | Immune system | Conventional NK | CAPG | KLRC2 |
| 1 | Immune system | Conventional NK | CLIC2 | KLRC2 |
| 1 | Immune system | Conventional NK | ZFHX3 | KLRC2 |
| 1 | Immune system | Conventional NK | KLF4 | KLRC2 |
| 1 | Immune system | Conventional NK | C1orf220 | KLRC2 |
| 1 | Immune system | Conventional NK | NUDT10 | KLRC2 |
| 1 | Immune system | Conventional NK | OSM | KLRC2 |
| 1 | Immune system | Conventional NK | RBBP8 | KLRC2 |
| 1 | Immune system | Conventional NK | LOC101929698 | KLRC2 |
| 1 | Immune system | Conventional NK | LPP-AS2 | KLRC2 |
| 1 | Immune system | Conventional NK | DAB2 | KLRC2 |
| 1 | Immune system | Conventional NK | GZMK | KLRC2 |
| 1 | Immune system | Conventional NK | LOC388813 | KLRC2 |
| 1 | Immune system | Conventional NK | GPX1 | KLRC2 |
| 1 | Immune system | Conventional NK | NFKBIZ | KLRC2 |
| 1 | Immune system | Conventional NK | ZNF300 | KLRC2 |
| 1 | Immune system | Conventional NK | LINC00865 | KLRC2 |
| 1 | Immune system | Conventional NK | WASF1 | KLRC2 |
| 1 | Immune system | Conventional NK | SCARF1 | KLRC2 |
| 1 | Immune system | Conventional NK | TLR9 | KLRC2 |
| 1 | Immune system | Conventional NK | NR4A1 | KLRC2 |
| 1 | Immune system | Conventional NK | UBR5-AS1 | KLRC2 |
| 1 | Immune system | Conventional NK | MS4A14 | KLRC2 |
| 1 | Immune system | Conventional NK | ARL11 | KLRC2 |

| | | | | |
|---|---|---|---|---|
| 1 | Immune system | Conventional NK | OPN3 | KLRC2 |
| 1 | Immune system | Conventional NK | CCRL2 | KLRC2 |
| 1 | Immune system | Conventional NK | AHRR | KLRC2 |
| 1 | Immune system | Conventional NK | KCTD12 | KLRC2 |
| 1 | Immune system | Conventional NK | CD5 | KLRC2 |
| 1 | Immune system | Conventional NK | GSTM2 | KLRC2 |
| 1 | Immune system | Conventional NK | ZNF366 | KLRC2 |
| 1 | Immune system | Conventional NK | SH3BP4 | KLRC2 |
| 1 | Immune system | Conventional NK | LRRC25 | KLRC2 |
| 1 | Immune system | Conventional NK | FAM132B | KLRC2 |
| 1 | Immune system | Conventional NK | FLT3 | KLRC2 |
| 1 | Immune system | Conventional NK | HTR3A | KLRC2 |
| 1 | Immune system | Conventional NK | UBD | KLRC2 |
| 1 | Immune system | Conventional NK | A2M | KLRC2 |
| 1 | Immune system | Conventional NK | GPR82 | KLRC2 |
| 1 | Immune system | Conventional NK | ZFP28 | KLRC2 |
| 1 | Immune system | Conventional NK | ST14 | KLRC2 |
| 1 | Immune system | Conventional NK | LYZ | KLRC2 |
| 1 | Immune system | Conventional NK | PLEKHG7 | KLRC2 |
| 1 | Immune system | Conventional NK | SLC7A7 | KLRC2 |
| 1 | Immune system | Conventional NK | RCAN3 | KLRC2 |
| 1 | Immune system | Conventional NK | LOC100128398 | KLRC2 |
| 1 | Immune system | Conventional NK | SIRPA | KLRC2 |
| 1 | Immune system | Conventional NK | PLAU | KLRC2 |
| 1 | Immune system | Conventional NK | LYL1 | KLRC2 |
| 1 | Immune system | Conventional NK | MICAL2 | KLRC2 |
| 1 | Immune system | Conventional NK | RAMP1 | KLRC2 |
| 1 | Immune system | Conventional NK | INPP5A | KLRC2 |
| 1 | Immune system | Conventional NK | NT5DC2 | KLRC2 |
| 1 | Immune system | Conventional NK | GNG4 | KLRC2 |
| 1 | Immune system | Conventional NK | ZNF662 | KLRC2 |
| 1 | Immune system | Conventional NK | CDCP1 | KLRC2 |
| 1 | Immune system | Conventional NK | ZNF334 | KLRC2 |
| 1 | Immune system | Conventional NK | LOC440896 | KLRC2 |
| 1 | Immune system | Conventional NK | AMPD3 | KLRC2 |
| 1 | Immune system | Conventional NK | IGLL5 | KLRC2 |
| 1 | Immune system | Conventional NK | CYP27A1 | KLRC2 |
| 1 | Immune system | Conventional NK | LILRA4 | KLRC2 |
| 1 | Immune system | Conventional NK | LAMA5 | KLRC2 |
| 1 | Immune system | Conventional NK | DIRAS1 | KLRC2 |
| 1 | Immune system | Conventional NK | CTSZ | KLRC2 |
| 1 | Immune system | Conventional NK | NSUN7 | KLRC2 |
| 1 | Immune system | Conventional NK | ACTN1 | KLRC2 |
| 1 | Immune system | Conventional NK | PPFIBP2 | KLRC2 |
| 1 | Immune system | Conventional NK | FAM177B | KLRC2 |
| 1 | Immune system | Conventional NK | LINC01547 | KLRC2 |
| 1 | Immune system | Conventional NK | TIAM1 | KLRC2 |
| 1 | Immune system | Conventional NK | NCCRP1 | KLRC2 |
| 1 | Immune system | Conventional NK | CD86 | KLRC2 |
| 1 | Immune system | Conventional NK | COL4A4 | KLRC2 |
| 1 | Immune system | Conventional NK | ARMCX2 | KLRC2 |
| 1 | Immune system | Conventional NK | MFSD2A | KLRC2 |
| 1 | Immune system | Conventional NK | SLC2A10 | KLRC2 |
| 1 | Immune system | Conventional NK | MIR4772 | KLRC2 |
| 1 | Immune system | Conventional NK | PXDC1 | KLRC2 |
| 1 | Immune system | Conventional NK | PFKFB2 | KLRC2 |
| 1 | Immune system | Conventional NK | CKAP4 | KLRC2 |
| 1 | Immune system | Conventional NK | FAM20C | KLRC2 |

| | | | | |
|---|---|---|---|---|
| 1 | Immune system | Conventional NK | MIR600HG | KLRC2 |
| 1 | Immune system | Conventional NK | ARRDC4 | KLRC2 |
| 1 | Immune system | Conventional NK | MPZL3 | KLRC2 |
| 1 | Immune system | Conventional NK | CD109 | KLRC2 |
| 1 | Immune system | Conventional NK | TNFSF4 | KLRC2 |
| 1 | Immune system | Conventional NK | ARHGAP32 | KLRC2 |
| 1 | Immune system | Conventional NK | CCNJL | KLRC2 |
| 1 | Immune system | Conventional NK | NREP | KLRC2 |
| 1 | Immune system | Conventional NK | METTL1 | KLRC2 |
| 1 | Immune system | Conventional NK | ARHGEF40 | KLRC2 |
| 1 | Immune system | Conventional NK | ABCB4 | KLRC2 |
| 1 | Immune system | Conventional NK | BCAT1 | KLRC2 |
| 1 | Immune system | Conventional NK | KCTD15 | KLRC2 |
| 1 | Immune system | Conventional NK | SATB1-AS1 | KLRC2 |
| 1 | Immune system | Conventional NK | ENTPD1 | KLRC2 |
| 1 | Immune system | Conventional NK | MEX3B | KLRC2 |
| 1 | Immune system | Conventional NK | ENPP3 | KLRC2 |
| 1 | Immune system | Conventional NK | ARHGAP31 | KLRC2 |
| 1 | Immune system | Conventional NK | ADAMTS1 | KLRC2 |
| 1 | Immune system | Conventional NK | MMD | KLRC2 |
| 1 | Immune system | Conventional NK | CEACAM19 | KLRC2 |
| 1 | Immune system | Conventional NK | TVP23A | KLRC2 |
| 1 | Immune system | Conventional NK | FAM110C | KLRC2 |
| 1 | Immune system | Conventional NK | ADCY6 | KLRC2 |
| 1 | Immune system | Conventional NK | CHPT1 | KLRC2 |
| 1 | Immune system | Conventional NK | NRGN | KLRC2 |
| 1 | Immune system | Conventional NK | CYP2S1 | KLRC2 |
| 1 | Immune system | Conventional NK | MICAL3 | KLRC2 |
| 1 | Immune system | Conventional NK | TPST1 | KLRC2 |
| 1 | Immune system | Conventional NK | TGM5 | KLRC2 |
| 1 | Immune system | Conventional NK | SAMD4A | KLRC2 |
| 1 | Immune system | Conventional NK | TNS3 | KLRC2 |
| 1 | Immune system | Conventional NK | CXXC5 | KLRC2 |
| 1 | Immune system | Conventional NK | ME1 | KLRC2 |
| 1 | Immune system | Conventional NK | CLEC12A | KLRC2 |
| 1 | Immune system | Conventional NK | TBC1D4 | KLRC2 |
| 1 | Immune system | Conventional NK | TNFRSF21 | KLRC2 |
| 1 | Immune system | Conventional NK | KLF8 | KLRC2 |
| 1 | Immune system | Conventional NK | PID1 | KLRC2 |
| 1 | Immune system | Conventional NK | RIN2 | KLRC2 |
| 1 | Immune system | Conventional NK | SMAD1 | KLRC2 |
| 1 | Immune system | Conventional NK | CNKSR2 | KLRC2 |
| 1 | Immune system | Conventional NK | FCRL4 | KLRC2 |
| 1 | Immune system | Conventional NK | ALPL | KLRC2 |
| 1 | Immune system | Conventional NK | ZNF677 | KLRC2 |
| 1 | Immune system | Conventional NK | TLR4 | KLRC2 |
| 1 | Immune system | Conventional NK | COL23A1 | KLRC2 |
| 1 | Immune system | Conventional NK | EFHC2 | KLRC2 |
| 1 | Immune system | Conventional NK | NRARP | KLRC2 |
| 1 | Immune system | Conventional NK | TMIGD2 | KLRC2 |
| 1 | Immune system | Conventional NK | CEP170B | KLRC2 |
| 1 | Immune system | Conventional NK | TAF4B | KLRC2 |
| 1 | Immune system | Conventional NK | PNOC | KLRC2 |
| 1 | Immune system | Conventional NK | AKAP2 | KLRC2 |
| 1 | Immune system | Conventional NK | CARD9 | KLRC2 |
| 1 | Immune system | Conventional NK | ATP6V1C2 | KLRC2 |
| 1 | Immune system | Conventional NK | AKT1S1 | KLRC2 |
| 1 | Immune system | Conventional NK | EEPD1 | KLRC2 |

| | | | | |
|---|---|---|---|---|
| 1 | Immune system | Conventional NK | CST3 | KLRC2 |
| 1 | Immune system | Conventional NK | TBXA2R | KLRC2 |
| 1 | Immune system | Conventional NK | LOC613266 | KLRC2 |
| 1 | Immune system | Conventional NK | AMOTL1 | KLRC2 |
| 1 | Immune system | Conventional NK | ECHDC3 | KLRC2 |
| 1 | Immune system | Conventional NK | DOCK4 | KLRC2 |
| 1 | Immune system | Conventional NK | ZMYND15 | KLRC2 |
| 1 | Immune system | Conventional NK | ARHGEF10L | KLRC2 |
| 1 | Immune system | Conventional NK | TTTY14 | KLRC2 |
| 1 | Immune system | Conventional NK | FNBP1L | KLRC2 |
| 1 | Immune system | Conventional NK | THBD | KLRC2 |
| 1 | Immune system | Conventional NK | CYBB | KLRC2 |
| 1 | Immune system | Conventional NK | VAV2 | KLRC2 |
| 1 | Immune system | Conventional NK | NLRP3 | KLRC2 |
| 1 | Immune system | Conventional NK | BEST1 | KLRC2 |
| 1 | Immune system | Conventional NK | PRKAR2B | KLRC2 |
| 1 | Immune system | Conventional NK | LTBP3 | KLRC2 |
| 1 | Immune system | Conventional NK | PVRL2 | KLRC2 |
| 1 | Immune system | Conventional NK | PADI2 | KLRC2 |
| 1 | Immune system | Conventional NK | HSP90AB1 | KLRC2 |
| 1 | Immune system | Conventional NK | PLXNA4 | KLRC2 |
| 1 | Immune system | Conventional NK | MYO1C | KLRC2 |
| 1 | Immune system | Conventional NK | DYSF | KLRC2 |
| 1 | Immune system | Conventional NK | SCARB1 | KLRC2 |
| 1 | Immune system | Conventional NK | EPB41L2 | KLRC2 |
| 1 | Immune system | Conventional NK | MYL9 | KLRC2 |
| 1 | Immune system | Conventional NK | LTBR | KLRC2 |
| 1 | Immune system | Conventional NK | MYO1B | KLRC2 |
| 1 | Immune system | Conventional NK | CD93 | KLRC2 |
| 1 | Immune system | Conventional NK | SLCO5A1 | KLRC2 |
| 1 | Immune system | Conventional NK | IL23A | KLRC2 |
| 1 | Immune system | Conventional NK | DENND6B | KLRC2 |
| 1 | Immune system | Conventional NK | NRP1 | KLRC2 |
| 1 | Immune system | Conventional NK | RAPGEF5 | KLRC2 |
| 1 | Immune system | Conventional NK | ZNF471 | KLRC2 |
| 1 | Immune system | Conventional NK | PIP5K1B | KLRC2 |
| 1 | Immune system | Conventional NK | BACH2 | KLRC2 |
| 1 | Immune system | Conventional NK | NLN | KLRC2 |
| 1 | Immune system | Conventional NK | KLK1 | KLRC2 |
| 1 | Immune system | Conventional NK | JAG1 | KLRC2 |
| 1 | Immune system | Conventional NK | LY96 | KLRC2 |
| 1 | Immune system | Conventional NK | SPATA6 | KLRC2 |
| 1 | Immune system | Conventional NK | ATF5 | KLRC2 |
| 1 | Immune system | Conventional NK | IRF8 | KLRC2 |
| 1 | Immune system | Conventional NK | GAS6-AS1 | KLRC2 |
| 1 | Immune system | Conventional NK | IZUMO4 | KLRC2 |
| 1 | Immune system | Conventional NK | IL1A | KLRC2 |
| 1 | Immune system | Conventional NK | P2RY6 | KLRC2 |
| 1 | Immune system | Conventional NK | UACA | KLRC2 |
| 1 | Immune system | Conventional NK | PDIA5 | KLRC2 |
| 1 | Immune system | Conventional NK | ABTB2 | KLRC2 |
| 1 | Immune system | Conventional NK | DCHS1 | KLRC2 |
| 1 | Immune system | Conventional NK | VDR | KLRC2 |
| 1 | Immune system | Conventional NK | CD80 | KLRC2 |
| 1 | Immune system | Conventional NK | GPR157 | KLRC2 |
| 1 | Immune system | Conventional NK | CXCL16 | KLRC2 |
| 1 | Immune system | Conventional NK | CD1D | KLRC2 |
| 1 | Immune system | Conventional NK | FMNL2 | KLRC2 |

| | | | | |
|---|---|---|---|---|
| 1 | Immune system | Conventional NK | BACE2 | KLRC2 |
| 1 | Immune system | Conventional NK | ZBTB32 | KLRC2 |
| 1 | Immune system | Conventional NK | SDPR | KLRC2 |
| 1 | Immune system | Conventional NK | HAPLN3 | KLRC2 |
| 1 | Immune system | Conventional NK | PRAM1 | KLRC2 |
| 1 | Immune system | Conventional NK | H1F0 | KLRC2 |
| 1 | Immune system | Conventional NK | PAICS | KLRC2 |
| 1 | Immune system | Conventional NK | LINC00899 | KLRC2 |
| 1 | Immune system | Conventional NK | MED12L | KLRC2 |
| 1 | Immune system | Conventional NK | GBGT1 | KLRC2 |
| 1 | Immune system | Conventional NK | WNT16 | KLRC2 |
| 1 | Immune system | Conventional NK | SALL4 | KLRC2 |
| 1 | Immune system | Conventional NK | BAHCC1 | KLRC2 |
| 1 | Immune system | Conventional NK | CLCN4 | KLRC2 |
| 1 | Immune system | Conventional NK | TGFA | KLRC2 |
| 1 | Immune system | Conventional NK | SHF | KLRC2 |
| 1 | Immune system | Conventional NK | HOMER3 | KLRC2 |
| 1 | Immune system | Conventional NK | UNC93B1 | KLRC2 |
| 1 | Immune system | Conventional NK | GYPE | KLRC2 |
| 1 | Immune system | Conventional NK | PRELID2 | KLRC2 |
| 1 | Immune system | Conventional NK | GPLD1 | KLRC2 |
| 1 | Immune system | Conventional NK | XXYLT1-AS2 | KLRC2 |
| 1 | Immune system | Conventional NK | GPR34 | KLRC2 |
| 1 | Immune system | Conventional NK | SHTN1 | KLRC2 |
| 1 | Immune system | Conventional NK | NLRC4 | KLRC2 |
| 1 | Immune system | Conventional NK | CSTA | KLRC2 |
| 1 | Immune system | Conventional NK | ANXA9 | KLRC2 |
| 1 | Immune system | Conventional NK | JDP2 | KLRC2 |
| 1 | Immune system | Conventional NK | FCGRT | KLRC2 |
| 1 | Immune system | Conventional NK | PLXDC2 | KLRC2 |
| 1 | Immune system | Conventional NK | NME1 | KLRC2 |
| 1 | Immune system | Conventional NK | SMPDL3B | KLRC2 |
| 1 | Immune system | Conventional NK | PRCD | KLRC2 |
| 1 | Immune system | Conventional NK | PDGFC | KLRC2 |
| 1 | Immune system | Conventional NK | IFI30 | KLRC2 |
| 1 | Immune system | Conventional NK | UST | KLRC2 |
| 1 | Immune system | Conventional NK | PACSIN1 | KLRC2 |
| 1 | Immune system | Conventional NK | SNX9 | KLRC2 |
| 1 | Immune system | Conventional NK | SLC41A2 | KLRC2 |
| 1 | Immune system | Conventional NK | GLT1D1 | KLRC2 |
| 1 | Immune system | Conventional NK | LOC102724552 | KLRC2 |
| 1 | Immune system | Conventional NK | KRT81 | KLRC2 |
| 1 | Immune system | Conventional NK | GADD45G | KLRC2 |
| 1 | Immune system | Conventional NK | LMO2 | KLRC2 |
| 1 | Immune system | Conventional NK | SLC40A1 | KLRC2 |
| 1 | Immune system | Conventional NK | GPR162 | KLRC2 |
| 1 | Immune system | Conventional NK | APBB2 | KLRC2 |
| 1 | Immune system | Conventional NK | LRRC75B | KLRC2 |
| 1 | Immune system | Conventional NK | COL4A3 | KLRC2 |
| 1 | Immune system | Conventional NK | ESPL1 | KLRC2 |
| 1 | Immune system | Conventional NK | RRAD | KLRC2 |
| 1 | Immune system | Conventional NK | EMID1 | KLRC2 |
| 1 | Immune system | Conventional NK | NET1 | KLRC2 |
| 1 | Immune system | Conventional NK | WNT3 | KLRC2 |
| 1 | Immune system | Conventional NK | HSF5 | KLRC2 |
| 1 | Immune system | Conventional NK | MRC2 | KLRC2 |
| 1 | Immune system | Conventional NK | SPARC | KLRC2 |
| 1 | Immune system | Conventional NK | SNX22 | KLRC2 |

| | | | | |
|---|---|---|---|---|
| 1 | Immune system | Conventional NK | C4orf32 | KLRC2 |
| 1 | Immune system | Conventional NK | FGD4 | KLRC2 |
| 1 | Immune system | Conventional NK | HOMER2 | KLRC2 |
| 1 | Immune system | Conventional NK | RIN1 | KLRC2 |
| 1 | Immune system | Conventional NK | CRYBG3 | KLRC2 |
| 1 | Immune system | Conventional NK | P2RX5-TAX1BP3 | KLRC2 |
| 1 | Immune system | Conventional NK | CPNE7 | KLRC2 |
| 1 | Immune system | Conventional NK | TSPAN18 | KLRC2 |
| 1 | Immune system | Conventional NK | LINC01535 | KLRC2 |
| 1 | Immune system | Conventional NK | NT5E | KLRC2 |
| 1 | Immune system | Conventional NK | NEDD4L | KLRC2 |
| 1 | Immune system | Conventional NK | SLC11A1 | KLRC2 |
| 1 | Immune system | Conventional NK | SELL | KLRC2 |
| 1 | Immune system | Conventional NK | CD200R1 | KLRC2 |
| 1 | Immune system | Conventional NK | TRIP10 | KLRC2 |
| 1 | Immune system | Conventional NK | REPS2 | KLRC2 |
| 1 | Immune system | Conventional NK | SCIMP | KLRC2 |
| 1 | Immune system | Conventional NK | ACP5 | KLRC2 |
| 1 | Immune system | Conventional NK | LPCAT2 | KLRC2 |
| 1 | Immune system | Conventional NK | PDE9A | KLRC2 |
| 1 | Immune system | Conventional NK | INSR | KLRC2 |
| 1 | Immune system | Conventional NK | L3MBTL4 | KLRC2 |
| 1 | Immune system | Conventional NK | PKIB | KLRC2 |
| 1 | Immune system | Conventional NK | MGAT5B | KLRC2 |
| 1 | Immune system | Conventional NK | MGC70870 | KLRC2 |
| 2 | Immune system | Transitional NK | FGFBP2 | |
| 2 | Immune system | Transitional NK | GZMB | |
| 2 | Immune system | Transitional NK | GZMK | |
| 2 | Immune system | Transitional NK | FCGR3A | |
| 2 | Immune system | Transitional NK | SPON2 | |
| 2 | Immune system | Transitional NK | S100A4 | |
| 2 | Immune system | Transitional NK | B2M | |
| 2 | Immune system | Transitional NK | GZMH | |
| 2 | Immune system | Transitional NK | NKG7 | |
| 2 | Immune system | Transitional NK | LGALS1 | |
| 2 | Immune system | Transitional NK | CST7 | |
| 2 | Immune system | Transitional NK | MYOM2 | |
| 2 | Immune system | Transitional NK | PRSS23 | |
| 2 | Immune system | Transitional NK | CMC1 | |
| 2 | Immune system | Transitional NK | S100A6 | |
| 2 | Immune system | Transitional NK | XCL1 | |
| 2 | Immune system | Transitional NK | HLA-C | |
| 2 | Immune system | Transitional NK | CYBA | |
| 2 | Immune system | Transitional NK | PRF1 | |
| 2 | Immune system | Transitional NK | PABPC1 | |
| 2 | Immune system | Transitional NK | CD44 | |
| 2 | Immune system | Transitional NK | XCL2 | |
| 2 | Immune system | Transitional NK | ARPC2 | |
| 2 | Immune system | Transitional NK | PIK3R1 | |
| 2 | Immune system | Transitional NK | CCL4 | |
| 2 | Immune system | Transitional NK | SH3BGRL3 | |
| 2 | Immune system | Transitional NK | CD27 | |
| 2 | Immune system | Transitional NK | ITGB2 | |
| 2 | Immune system | Transitional NK | CD99 | |
| 2 | Immune system | Transitional NK | FCRL6 | |
| 2 | Immune system | Transitional NK | RP11-277L2.4 | |
| 2 | Immune system | Transitional NK | GNLY | |
| 2 | Immune system | Transitional NK | GPR56 | |

| | | | |
|---|---|---|---|
| 2 | Immune system | Transitional NK | ACTB |
| 2 | Immune system | Transitional NK | CLDND1 |
| 2 | Immune system | Transitional NK | LAIR2 |
| 2 | Immune system | Transitional NK | AKR1C3 |
| 2 | Immune system | Transitional NK | CD52 |
| 2 | Immune system | Transitional NK | IFRD1 |
| 2 | Immune system | Transitional NK | CX3CR1 |
| 2 | Immune system | Transitional NK | IGFBP7 |
| 2 | Immune system | Transitional NK | ANXA1 |
| 2 | Immune system | Transitional NK | DBI |
| 2 | Immune system | Transitional NK | ZEB2 |
| 2 | Immune system | Transitional NK | ITGB7 |
| 2 | Immune system | Transitional NK | TMEM173 |
| 2 | Immune system | Transitional NK | SPTSSB |
| 2 | Immune system | Transitional NK | EMP3 |
| 2 | Immune system | Transitional NK | EIF3G |
| 2 | Immune system | Transitional NK | TCF7 |
| 2 | Immune system | Transitional NK | CTSD |
| 2 | Immune system | Transitional NK | CD247 |
| 2 | Immune system | Transitional NK | SPN |
| 2 | Immune system | Transitional NK | DSTN |
| 2 | Immune system | Transitional NK | UCP2 |
| 2 | Immune system | Transitional NK | TTC38 |
| 2 | Immune system | Transitional NK | GZMA |
| 2 | Immune system | Transitional NK | KLRC1 |
| 2 | Immune system | Transitional NK | PTGDS |
| 2 | Immune system | Transitional NK | AES |
| 2 | Immune system | Transitional NK | GTF3C1 |
| 2 | Immune system | Transitional NK | CAP1 |
| 2 | Immune system | Transitional NK | PDCD4 |
| 2 | Immune system | Transitional NK | TIMP1 |
| 2 | Immune system | Transitional NK | CAPN12 |
| 2 | Immune system | Transitional NK | SH2D1A |
| 2 | Immune system | Transitional NK | SYNGR1 |
| 2 | Immune system | Transitional NK | DHRS7 |
| 2 | Immune system | Transitional NK | FLNA |
| 2 | Immune system | Transitional NK | ABHD17A |
| 2 | Immune system | Transitional NK | SSBP3 |
| 2 | Immune system | Transitional NK | SCP2 |
| 2 | Immune system | Transitional NK | CD74 |
| 2 | Immune system | Transitional NK | LAIR1 |
| 2 | Immune system | Transitional NK | CNN2 |
| 2 | Immune system | Transitional NK | IL2RB |
| 2 | Immune system | Transitional NK | TMEM107 |
| 2 | Immune system | Transitional NK | CCL5 |
| 2 | Immune system | Transitional NK | CCL3 |
| 2 | Immune system | Transitional NK | IFIT2 |
| 2 | Immune system | Transitional NK | WDR74 |
| 2 | Immune system | Transitional NK | C12orf57 |
| 2 | Immune system | Transitional NK | C20orf24 |
| 3 | Immune system | Inflamed NK | KLRB1 |
| 3 | Immune system | Inflamed NK | PMAIP1 |
| 3 | Immune system | Inflamed NK | IFIT3 |
| 3 | Immune system | Inflamed NK | IFIT2 |
| 3 | Immune system | Inflamed NK | CCL3 |
| 3 | Immune system | Inflamed NK | CCL3L1 |
| 3 | Immune system | Inflamed NK | CCL3L3 |
| 3 | Immune system | Inflamed NK | TNF |

| | | | |
|---|---|---|---|
| 3 | Immune system | Inflamed NK | NFKBIZ |
| 3 | Immune system | Inflamed NK | CCL4 |
| 3 | Immune system | Inflamed NK | OASL |
| 3 | Immune system | Inflamed NK | ISG15 |
| 3 | Immune system | Inflamed NK | GCA |
| 3 | Immune system | Inflamed NK | PPP1R15A |
| 3 | Immune system | Inflamed NK | CCL4L2 |
| 3 | Immune system | Inflamed NK | IFIT1 |
| 3 | Immune system | Inflamed NK | RPS4Y1 |
| 3 | Immune system | Inflamed NK | LTA |
| 3 | Immune system | Inflamed NK | TNFAIP3 |
| 3 | Immune system | Inflamed NK | FOS |
| 3 | Immune system | Inflamed NK | CCL4L1 |
| 3 | Immune system | Inflamed NK | ZC3HAV1 |
| 3 | Immune system | Inflamed NK | CD69 |
| 3 | Immune system | Inflamed NK | AC069363.1 |
| 3 | Immune system | Inflamed NK | LAX1 |
| 3 | Immune system | Inflamed NK | HELB |
| 3 | Immune system | Inflamed NK | YBX1 |
| 3 | Immune system | Inflamed NK | ETV3 |
| 3 | Immune system | Inflamed NK | CCL5 |
| 3 | Immune system | Inflamed NK | RP1-40E16.12 |
| 3 | Immune system | Inflamed NK | MYL12A |
| 3 | Immune system | Inflamed NK | EHD4 |
| 3 | Immune system | Inflamed NK | PTGER4 |
| 3 | Immune system | Inflamed NK | APOL6 |
| 3 | Immune system | Inflamed NK | KLRB1 |
| 3 | Immune system | Inflamed NK | FTH1 |
| 3 | Immune system | Inflamed NK | XIST |
| 3 | Immune system | Inflamed NK | NANS |
| 3 | Immune system | Inflamed NK | ZEB2-AS1 |
| 3 | Immune system | Inflamed NK | TMSB4X |
| 3 | Immune system | Inflamed NK | CD7 |
| 3 | Immune system | Inflamed NK | CDKN1A |
| 3 | Immune system | Inflamed NK | DDX3Y |
| 3 | Immune system | Inflamed NK | ARHGAP11B |
| 3 | Immune system | Inflamed NK | ZBP1 |
| 3 | Immune system | Inflamed NK | NR4A2 |
| 3 | Immune system | Inflamed NK | MT2A |
| 3 | Immune system | Inflamed NK | TTTY15 |
| 3 | Immune system | Inflamed NK | ACTB |
| 3 | Immune system | Inflamed NK | HERC5 |
| 3 | Immune system | Inflamed NK | PRF1 |
| 3 | Immune system | Inflamed NK | C20orf24 |
| 3 | Immune system | Inflamed NK | DDIT4 |
| 3 | Immune system | Inflamed NK | LINC-PINT |
| 3 | Immune system | Inflamed NK | ANKRD28 |
| 3 | Immune system | Inflamed NK | FGR |
| 3 | Immune system | Inflamed NK | TNIP1 |
| 3 | Immune system | Inflamed NK | MYL12B |
| 3 | Immune system | Inflamed NK | RBM39 |
| 3 | Immune system | Inflamed NK | RGS1 |
| 3 | Immune system | Inflamed NK | TXNIP |
| 3 | Immune system | Inflamed NK | GZMK |
| 3 | Immune system | Inflamed NK | FCER1G |
| 3 | Immune system | Inflamed NK | EFHD2 |
| 3 | Immune system | Inflamed NK | ZBTB20 |
| 3 | Immune system | Inflamed NK | RAC2 |

| | | | |
|---|---|---|---|
| 3 | Immune system | Inflamed NK | SBF2 |
| 3 | Immune system | Inflamed NK | CTSW |
| 3 | Immune system | Inflamed NK | CORO1A |
| 3 | Immune system | Inflamed NK | GZMA |
| 3 | Immune system | Inflamed NK | CLIC1 |
| 3 | Immune system | Inflamed NK | BTG1 |
| 3 | Immune system | Inflamed NK | IGFBP7 |
| 3 | Immune system | Inflamed NK | SFT2D2 |
| 3 | Immune system | Inflamed NK | CLEC2B |
| 3 | Immune system | Inflamed NK | CMC1 |
| 3 | Immune system | Inflamed NK | ARPC5 |
| 3 | Immune system | Inflamed NK | CLIC3 |
| 3 | Immune system | Inflamed NK | CFL1 |
| 3 | Immune system | Inflamed NK | CTD-2037K23.2 |
| 3 | Immune system | Inflamed NK | CD160 |
| 3 | Immune system | Inflamed NK | KLF9 |
| 3 | Immune system | Inflamed NK | BTG2 |
| 3 | Immune system | Inflamed NK | ZEB2 |
| 3 | Immune system | Inflamed NK | RHOB |
| 3 | Immune system | Inflamed NK | PPP2R5C |
| 3 | Immune system | Inflamed NK | MIA3 |
| 3 | Immune system | Inflamed NK | ANXA2 |
| 3 | Immune system | Inflamed NK | ELF1 |
| 3 | Immune system | Inflamed NK | C1orf162 |
| 3 | Immune system | Inflamed NK | PRDM1 |
| 4 | Immune system | Terminal NK | WDR74 |
| 4 | Immune system | Terminal NK | RPS27 |
| 4 | Immune system | Terminal NK | SNORD3A |
| 4 | Immune system | Terminal NK | TMEM107 |
| 4 | Immune system | Terminal NK | RPS29 |
| 4 | Immune system | Terminal NK | RPLP2 |
| 4 | Immune system | Terminal NK | RPL23A |
| 4 | Immune system | Terminal NK | RPL34 |
| 4 | Immune system | Terminal NK | RPS14 |
| 4 | Immune system | Terminal NK | RPL39 |
| 4 | Immune system | Terminal NK | SNORD3D |
| 4 | Immune system | Terminal NK | RPL21 |
| 4 | Immune system | Terminal NK | RPS18 |
| 4 | Immune system | Terminal NK | RPS25 |
| 4 | Immune system | Terminal NK | RPS12 |
| 4 | Immune system | Terminal NK | LINC00910 |
| 4 | Immune system | Terminal NK | RPL32 |
| 4 | Immune system | Terminal NK | RPL13 |
| 4 | Immune system | Terminal NK | RPS3 |
| 4 | Immune system | Terminal NK | RPS28 |
| 4 | Immune system | Terminal NK | RNU12 |
| 4 | Immune system | Terminal NK | RPL18A |
| 4 | Immune system | Terminal NK | SNORD3B-2 |
| 4 | Immune system | Terminal NK | RPS21 |
| 4 | Immune system | Terminal NK | RPL13A |
| 4 | Immune system | Terminal NK | HIST1H1E |
| 4 | Immune system | Terminal NK | C12orf57 |
| 4 | Immune system | Terminal NK | SPON2 |
| 4 | Immune system | Terminal NK | SNORD3B-1 |
| 4 | Immune system | Terminal NK | RPS2 |
| 4 | Immune system | Terminal NK | CLIC3 |
| 4 | Immune system | Terminal NK | RPS10 |
| 4 | Immune system | Terminal NK | FGFBP2 |

| | | | |
|---|---|---|---|
| 4 | Immune system | Terminal NK | RPL8 |
| 4 | Immune system | Terminal NK | GZMB |
| 4 | Immune system | Terminal NK | PRF1 |
| 4 | Immune system | Terminal NK | RP11-386I14.4 |
| 4 | Immune system | Terminal NK | FCGR3A |
| 4 | Immune system | Terminal NK | CD247 |
| 4 | Immune system | Terminal NK | CD160 |
| 4 | Immune system | Terminal NK | PTCH2 |
| 4 | Immune system | Terminal NK | RPL36A |
| 4 | Immune system | Terminal NK | FCER1G |
| 4 | Immune system | Terminal NK | CD38 |
| 4 | Immune system | Terminal NK | GZMK |
| 4 | Immune system | Terminal NK | RNU11 |
| 4 | Immune system | Terminal NK | HIST1H1D |
| 4 | Immune system | Terminal NK | NFKBIA |
| 4 | Immune system | Terminal NK | HIST2H2BE |
| 4 | Immune system | Terminal NK | MYL12A |
| 4 | Immune system | Terminal NK | RAC2 |
| 4 | Immune system | Terminal NK | PLAC8 |
| 4 | Immune system | Terminal NK | ACTB |
| 4 | Immune system | Terminal NK | RAP1B |
| 4 | Immune system | Terminal NK | NEAT1 |
| 4 | Immune system | Terminal NK | FGR |
| 4 | Immune system | Terminal NK | CD44 |
| 4 | Immune system | Terminal NK | HIST1H2BD |
| 4 | Immune system | Terminal NK | HIST1H1C |
| 4 | Immune system | Terminal NK | EVL |
| 4 | Immune system | Terminal NK | PTPRC |
| 4 | Immune system | Terminal NK | GZMA |
| 4 | Immune system | Terminal NK | HNRNPA2B1 |
| 4 | Immune system | Terminal NK | S1PR5 |
| 4 | Immune system | Terminal NK | PTPRCAP |
| 4 | Immune system | Terminal NK | GNG2 |
| 4 | Immune system | Terminal NK | IFRD1 |
| 4 | Immune system | Terminal NK | APMAP |
| 4 | Immune system | Terminal NK | ARPC5 |
| 4 | Immune system | Terminal NK | LIMD2 |
| 4 | Immune system | Terminal NK | HIST1H2BK |
| 4 | Immune system | Terminal NK | PRSS23 |
| 4 | Immune system | Terminal NK | UCP2 |
| 4 | Immune system | Terminal NK | CD52 |
| 4 | Immune system | Terminal NK | ZEB2 |
| 4 | Immune system | Terminal NK | MYOM2 |
| 4 | Immune system | Terminal NK | ANKRD20A4 |
| 4 | Immune system | Terminal NK | GPR56 |
| 4 | Immune system | Terminal NK | HNRNPK |
| 4 | Immune system | Terminal NK | CEP78 |
| 4 | Immune system | Terminal NK | FCRL6 |
| 4 | Immune system | Terminal NK | SELPLG |
| 4 | Immune system | Terminal NK | TTC38 |
| 4 | Immune system | Terminal NK | CX3CR1 |
| 4 | Immune system | Terminal NK | ANXA6 |
| 4 | Immune system | Terminal NK | DDX17 |
| 4 | Immune system | Terminal NK | CORO1A |
| 4 | Immune system | Terminal NK | PRKCH |
| 4 | Immune system | Terminal NK | AKR1C3 |
| 4 | Immune system | Terminal NK | PSMB9 |
| 4 | Immune system | Terminal NK | ADD3 |

| | | | |
|---|---|---|---|
| 4 | Immune system | Terminal NK | ARHGAP30 |
| 4 | Immune system | Terminal NK | PTPN6 |
| 4 | Immune system | Terminal NK | SNHG8 |
| 4 | Immune system | Terminal NK | CD3E |
| 4 | Immune system | Terminal NK | PLEK |
| 4 | Immune system | Terminal NK | SLFN5 |
| 4 | Immune system | Terminal NK | LAIR2 |
| 4 | Immune system | Terminal NK | DHRS7 |
| 4 | Immune system | Terminal NK | ACTR2 |
| 4 | Immune system | Terminal NK | OSTF1 |
| 4 | Immune system | Terminal NK | ALOX5AP |
| 4 | Immune system | Terminal NK | PMAIP1 |
| 4 | Immune system | Terminal NK | TNFRSF18 |
| 4 | Immune system | Terminal NK | TRAF3IP3 |
| 4 | Immune system | Terminal NK | NR4A2 |
| 4 | Immune system | Terminal NK | SAMD3 |
| 4 | Immune system | Terminal NK | IFI16 |
| 4 | Immune system | Terminal NK | PPP1R18 |
| 4 | Immune system | Terminal NK | XCL1 |
| 4 | Immune system | Terminal NK | FOS |
| 4 | Immune system | Terminal NK | DYNLL1 |
| 4 | Immune system | Terminal NK | XIST |
| 4 | Immune system | Terminal NK | LTB |
| 4 | Immune system | Terminal NK | NFKBIZ |

**Supplementary Table 2. NK Cell type annotation for scRNA clusters matched with scATAC-seq data**

| cluster | type | scores | ncells |
|---|---|---|---|
| 6 | Unknown | -5.326831035 | 57 |
| 2 | Unknown | -5.418221973 | 122 |
| 3 | Inflamed NK | 94.51435097 | 105 |
| 4 | Adaptive_NK_merged | 44.05848478 | 100 |
| 1 | Unknown | -34.45454893 | 138 |
| 7 | Transitional NK | 15.08796968 | 32 |
| 5 | Conventional NK | 512.9596661 | 62 |
| 0 | Unknown | -47.10830658 | 144 |

**Supplementary Table 3. NK Cell type annotation for scRNA clusters from Smartseq3 data**

| cluster | type | scores | ncells |
|---|---|---|---|
| Cluster_4 | Inflamed NK | 267.2537189 | 93 |
| Cluster_5 | Unknown | 16.2169182 | 81 |
| Cluster_7 | Adaptive_NK_merged | 188.3393251 | 59 |
| Cluster_1 | Adaptive_NK_merged | 545.0795913 | 141 |
| Cluster_3 | Terminal NK | 58.93839745 | 129 |
| Cluster_2 | Inflamed NK | 189.6161225 | 139 |
| Cluster_8 | Conventional NK | 88.19891363 | 28 |
| Cluster_0 | Unknown | -151.6160277 | 196 |
| Cluster_6 | Unknown | 7.00672643 | 64 |
